# Architectures and biochemical activities of Mtl1-Red1 MTREC helicase complexes

**DOI:** 10.64898/2026.08.18.745603

**Authors:** Lucas D. Repeta, Christopher D. Lima

**Affiliations:** Tri-Institutional PhD Program in Chemical Biology, Memorial Sloan Kettering Cancer Center, New York, NY; Structural Biology Program, Sloan Kettering Institute, New York, NY; Howard Hughes Medical Institute, New York, NY

## Abstract

RNA surveillance and decay is carried out in part by helicase containing complexes that identify, capture and sometimes modify RNA before delivering it to the RNA exosome complex for processing or degradation. The MTREC core complex includes a Mtr4-like protein (Mtl1) helicase and Red1 that works with other cofactors and the RNA exosome in Schizosaccharomyces pombe to degrade nuclear transcripts in processes that can result in formation of facultative heterochromatin. The activities of Mtl1 remain uncharacterized as do contributions of Red1 to Mtl1 within MTREC. Here, we reconstitute the MTREC core complex, resolve structures by cryo-electron microscopy, and compare MTREC activities to S. pombe Mtr4 and Mtl1. We show that Mtl1 is more active relative to MTREC and Mtr4, that MTREC binds RNA better than Mtl1, and that Red1 includes an autoinhibitory coiled-coil domain that dimerizes MTREC and contacts the Mtl1 RecA domains to disrupt its ATPase active site. Together, these data suggest that Red1 may endow MTREC to bind RNA while slowing translocation so that it remains associated with RNA long enough to chaperone it to the RNA exosome for processing or degradation.

## Introduction

Surveillance systems monitor RNA during and after synthesis to regulate the lifetime of RNA or degrade it to prevent accumulation of aberrant or misprocessed RNAs that could interfere with cellular processes ^1–5^. One such eukaryotic pathway includes factors that engage RNA substrates at their 3’ ends to prepare them for delivery to the highly conserved RNA exosome, a multi-subunit exoribonuclease complex that serves as a focal point for cytoplasmic and nuclear RNA surveillance ^6–11^. Prior to degradation, RNA targets are identified and delivered to the RNA exosome by several helicase-containing complexes that include the Mtr4 RNA helicase in addition to specialized pathway-specific RNA-binding protein cofactors that endow the complexes with the ability to identify and hold onto RNA substrates long enough to facilitate processing by the exosome^6,12–20^.

The spectrum of exosome cofactors and the RNAs they regulate is emerging, but structures and mechanisms for many of these upstream complexes remain less clear. Further, the number and variety of helicase-containing complexes that modulate RNA surveillance varies across eukaryotic species. In the budding yeast *Saccharomyces cerevisiae* where the RNA exosome was first discovered ^21^, the Trf4/5-Air1/2-Mtr4 (TRAMP) complex contributes to nuclear decay through post-transcriptional modification of RNA via its non-templated poly(A) polymerase activity ^22–26^. The Mtr4 helicase subunit of TRAMP interacts with the RNA exosome via contacts to exosome co-factors that help dock the helicase onto the exosome where it facilitates delivery of the substrate for processing or degradation ^10,25,27^. In the cytoplasm, exosome-mediated mRNA surveillance includes the ribosome-associated SKI complex composed of the Mtr4-related Ski2 helicase, the tetratricopeptide repeat (TPR) protein Ski3, and two subunits of the WD repeat protein Ski8 which targets the cytoplasmic form of the RNA exosome to mRNAs, miRNAs, and viral transcripts ^17,28,29^.

While the TRAMP and the SKI complex are conserved in human and the fission yeast *Schizosaccharomyces pombe*, TRAMP activities appear restricted to the nucleolus where it participates in the processing or degradation of RNA Pol I transcripts such as rRNA and snoRNAs ^14,30–32^. In these organisms, nuclear quality control is carried out by other helicase containing complexes, including the nuclear exosome targeting (NEXT) complex and the poly(A) exosome targeting (PAXT) connection ^14,18,19,33–37^ in humans and the Mtl1-Red1 core (MTREC) complex in fission yeast ^15,16,31^. Structural and biochemical characterization of NEXT revealed a ternary complex between the helicase MTR4, the zinc finger protein ZCCHC8, and the RRM-containing protein RBM7, which forms a stable homodimer mediated by ZCCHC8 ^13,35,38^. Available structures and biochemistry show that ZCCHC8 covers MTR4 surfaces that would be required for RNA translocation and cofactor-dependent docking of MTR4 onto the exosome core ^13,35,38^, suggesting a hierarchical remodeling of protein surfaces that would be required to coordinate RNA capture, delivery and translocation.

Unlike NEXT, much less is known about the architectures and activities of the PAXT connection ^18,19,34,39,40^ or MTREC ^15,16^ although both include scaffolding subunits that facilitate interactions with nascent ribonucleoprotein particles via contacts to nuclear cap binding proteins, Ars2, and the nuclear poly(A) binding protein among others. The PAXT connection core complex includes the MTR4 helicase and the C3H1 zinc-finger protein ZFC3H1 ^18^ while the MTREC core complex includes the Mtr4-like protein 1 helicase (Mtl1) and the C3H1 zinc-finger protein Red1 ^15,16,31^. While Mtl1 is highly related to Mtr4, Red1 shares only limited similarity to the PAXT subunit ZFC3H1 ^15,18,31,37^. Similar to human PAXT connection, MTREC is predicted to dynamically associate with RNA protein complexes including the 5’ cap binding complex-Ars2 (CBCA), the poly(A) binding protein Pab2, the zinc finger protein Red5, and the RRM-containing protein Rmn1 ^15,16,41–47^. Unlike PAXT, MTREC also binds the fission yeast specific proteins Mmi1 and Iss10 which, along with Erh1, help target MTREC to hexanucleotide determinant of selective removal (DSR) sequences in meiosis related mRNAs and certain ncRNAs ^48–52^. MTREC has also been implicated in post-transcriptional gene silencing (PTGS) pathways ^15,16,31,44,53^ where it is proposed to collaborate with the RNA-induced transcriptional silencing (RITS) complex to locate de-repressed regions of the genome, degrade spurious RNAs, and re-establish facultative heterochromatin ^54,55^. The connection between MTREC and PTGS pathways was the first indication that the exosome was directly involved in this type of gene regulation, and recent evidence has emerged suggesting that this connection may be conserved in higher eukaryotes including humans ^16,56–58^.

Recent efforts to characterize MTREC provided information on protein-protein interactions that organize the complex and classes of RNAs sensitive to deletions and mutations of MTREC genes ^59–61^. While some structural information was reported for hybrid systems ^59,60^, characterization of intact *S. pombe* MTREC core complexes at the biochemical and structural level is not yet available. Here, we isolate intact MTREC core complexes composed of Mtl1 and Red1 and characterize MTREC RNA binding and strand displacement activities relative to Mtl1 or Mtr4. We determine three single particle cryo-electron reconstructions including MTREC dimer structures that reveal interactions between Mtl1 and Red1 that contribute to RNA binding, ATPase activity and helicase function.

## RESULTS

### Biochemical comparison of Mtr4, Mtl1 and MTREC from fission yeast

Mtr4 and Mtl1 are reported to have little overlap in their cellular activities and appear to interact exclusively with TRAMP and MTREC associated proteins, respectively ^15,16,31^. Prior work revealed that Mtr4 proteins from fission yeast, budding yeast, and humans exhibit weak strand displacement or helicase activities that can be stimulated by interactions with exosome cofactors or cofactor complexes to efficiently translocate RNA substrates ^10,13,35^. While Mtr4 and Mtl1 share 55% sequence identity and are predicted to adopt similar overall folds, current AlphaFold models of Mtl1 do not reveal structural differences that would predict biochemical differences between these two helicases. Furthermore, Red1 was proposed as a scaffold within MTREC, but it remains unclear if it contributes to Mtl1 enzymatic activity ^16,59,60^. To determine if Mtl1 displays biochemical properties that distinguish it from Mtr4 or MTREC, full-length Mtr4, Mtl1, and MTREC were purified (Fig. 1a, Fig. S1) and assessed in biochemical assays for RNA binding and strand displacement (Fig. 1b-g, Fig. S2, Tables S1, S2).

**Figure 1.**
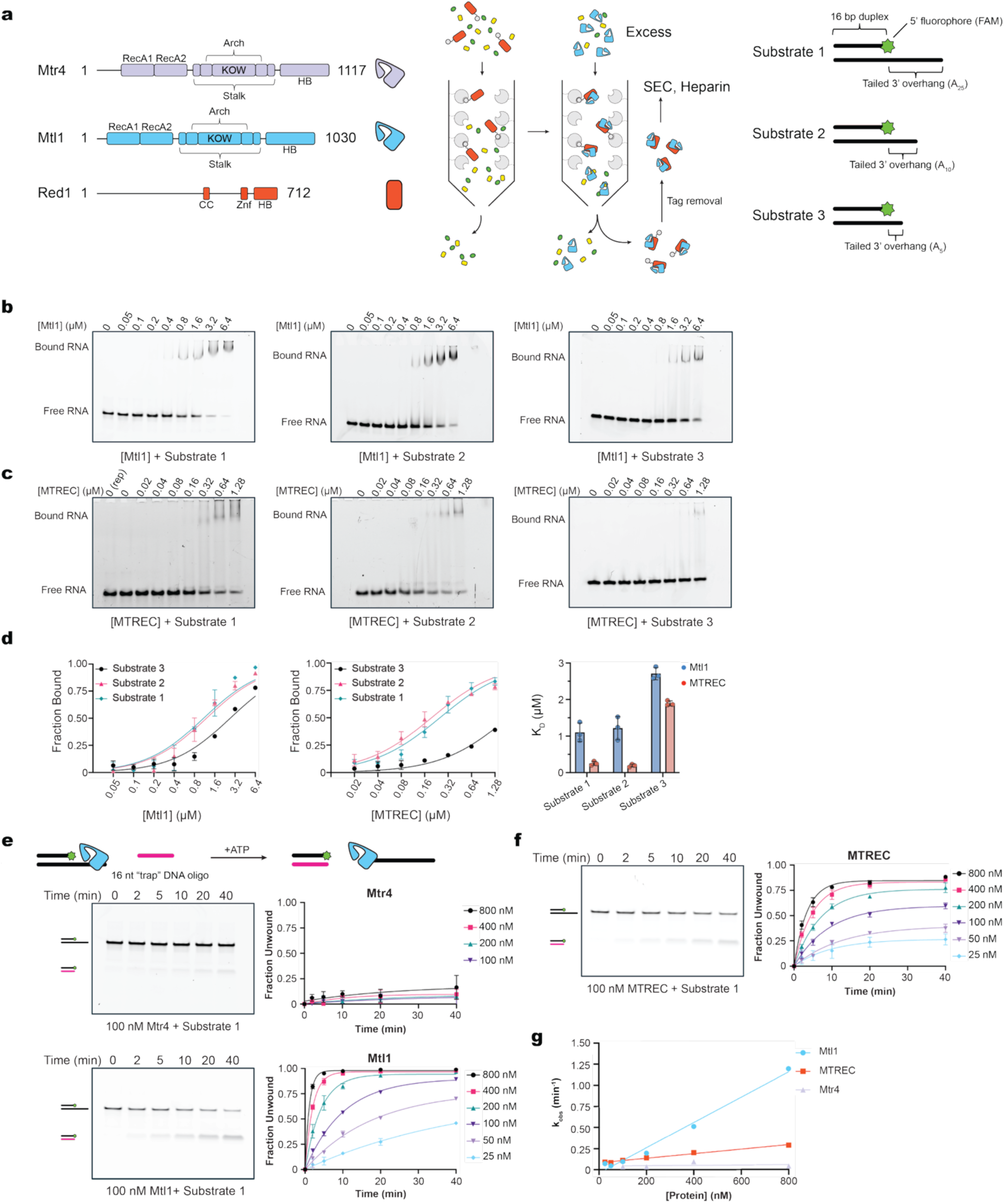
Biochemical activities of Mtr4, Mtl1 and MTREC from fission yeast. **(a)** Domain schematics for Mtr4, Mtl1, and Red1 (left). Cartoon depiction of methods used for MTREC complex formation by sequential binding of GST-tagged Red1 to glutathione resin followed by incubation with Mtl1 (middle). Cartoon schematic of RNA substrates used in binding and strand displacement assays (right). The 16 nt 5’ 6-carboxyfluorescein (FAM) labeled strand is annealed to a complementary RNA strand that also includes a 3’ poly(A) overhang of 5, 10, or 25 nt. **(b)** Representative gels for electrophoretic mobility shift assays (EMSAs) of Mtl1 with each RNA substrate. Positions of free and shifted RNA are labeled. **(c)** EMSAs of MTREC with each RNA substrate. **(d)** Graphs depicting quantification of Mtl1 and MTREC binding across concentrations of protein for each RNA substrate as well as calculated dissociation constants. **(e)** Cartoon depiction of strand displacement assay and representative gels of Mtl1 and Mtr4 with the 25 nt poly(A) overhang substrate 1. Bands corresponding to substrate and product RNA:DNA hybrid are displayed with a cartoon. **(f)** Strand displacement assay of MTREC with substrate 1. **(g)** Plot of the observed rate constants against the concentration of protein used in the strand displacement assays. Error bars represent the standard error of the mean (SEM) from 3 technical replicates as calculated by GraphPad Prism.

### MTREC binding to RNA requires a 3’ single stranded end of more than 5 nucleotides

Three distinct RNA substrates were generated to determine a suitable RNA substrate for comparative analysis between Mtr4, Mtl1 and MTREC. These RNA substrates included a 16 bp duplex with a 3’ poly(A) single stranded overhang of 5, 10, or 25 nucleotides (nt) (Fig. 1a). Binding assays were performed by incubating purified Mtl1 or MTREC with the RNA substrates in the presence of a nonhydrolyzable ATP analog AMPPNP. Consistent with structures and known biochemical properties for Mtr4 which prefers overhangs of at least 7 nucleotides^10,20^, Mtl1 exhibits weaker affinity for a 3’ 5 nt poly(A) overhang substrate with a K_D_ of approximately 2.7 μM ± 0.19 µM compared to substrates containing a single stranded 3’ overhang of 10 nt (1.2 μM ± 0.14 µM) or of 25 nt (1.1 μM ± 0.13 µM) (Fig. 1b,d; Fig. S2a, Table S1). MTREC bound poorly to the RNA substrate with a 5 nt 3’ overhang resulting in an estimated K_D_ of 1.9 μM ± 0.01 µM. Similar to Mtl1, MTREC preferred binding to RNA substrates with 10 or 25 nt poly(A) single stranded overhangs, however MTREC bound these substrates 5-fold better than Mtl1 with K_D_ values estimated at 0.191 ± 0.013 µM and 0.246 ± 0.017 µM, respectively (Fig. 1c,d; Fig. S2b, Table S1). In the absence of known Red1 RNA binding domains, it is unclear if increased affinity observed for MTREC is due to altering how Mtl1 engages RNA or if Red1 directly interacts with the RNA. Regardless, these data suggest that Red1 alters interactions with RNA in the context of MTREC in addition to its previously reported scaffolding function. These data also suggest that an RNA substrate with a 25 nt single stranded overhang is sufficient for comparing Mtr4, Mtl1 and MTREC activities in strand displacement assays.

### Mtl1 strand displacement activity is more efficient than Mtr4 and suppressed by Red1

To compare Mtr4, Mtl1 and MTREC, strand displacement assays were conducted with the RNA substrate 1 and initiated by adding ATP to samples of Mtr4, Mtl1 or MTREC that were preincubated with RNA substrate 1 (Fig. 1e). Aliquots were removed from each reaction at specified time points and analyzed by native PAGE to measure band intensities to determine the fraction of RNA unwound as a function of time.

Mtr4 catalyzed little to no detectable strand displacement even at concentrations up to 800 nM (Fig. 1e, Fig. S2c, Table S2) in line with prior observations that Mtr4 is a comparatively weak helicase that is enhanced through interactions with cofactors associated with the exosome ^10,13,35^. In contrast, Mtl1 exhibited higher activity with 50 nM Mtl1 displaying a rate of K_obs_ = 0.05 ± 0.01 (nM)(min^−1^) with a calculated half time of t_1/2_ = 15 min (Fig. 1e, Fig. S2d, Table S2). As stated earlier, Mtr4 and Mtl1 are predicted to share similar domain architectures, so it remains unclear what drives the observed differences in activities; however, it is clear from this analysis that they possess different intrinsic activities in this assay.

Strand displacement assays were next conducted with MTREC to determine if Red1 influenced strand displacement activity (Fig. 1f, Fig. S2e, Table S2). At lower concentrations, initial rates for MTREC were similar to Mtl1, as 50 nM MTREC achieved a rate of K_obs_ = 0.08 ± 0.01 (nM)(min^−1^) compared to K_obs_ = 0.05 ± 0.01 (nM)(min^−1^) for Mtl1. While initial rates were similar, 50 nM MTREC has a shorter half time of t_1/2_ = 8.3 min owing to a reduced reaction amplitude. 50 nM MTREC was calculated to achieve approximately 40% strand displacement while the equivalent concentration of Mtl1 reached 75% displacement. While it was anticipated that MTREC would be more efficient at displacement given its higher affinity for this substrate, Red1 appears to attenuate rather than enhance overall displacement, an effect that became more evident at higher concentrations. For instance, at 800 nM Mtl1 achieved a rate of K_obs_ = 1.19 ± 0.03 (nM)(min^−1^) with t_1/2_ = 0.58 min to reach an overall amplitude of 99%, but MTREC only reached a rate of K_obs_ = 0.3 ± 0.02 (nM)(min^−1^) and plateaued at 85% displacement with t_1/2_ = 2.4 min (Fig. 1f). Further, the rate of strand displacement scaled linearly with Mtl1 concentration while higher concentrations of MTREC appeared to inhibit the reaction (Fig. 1g). It is also noteworthy that while Mtl1 achieved complete displacement in just minutes at concentrations well below the measured K_D_, MTREC saturated at around 85% displacement at concentrations above its apparent K_D_ for this substrate, suggesting that it may be stuck on the ends of substrates under the conditions tested. A similar phenomenon was observed with the NEXT complex which appears to inhibit strand displacement at concentrations greater than 1 μM ^13^. These results are consistent with the concept that MTREC and NEXT may possess chaperone like properties that would enable them to stall on substrates long enough to be recognized for delivery to the RNA exosome.

### Cryo-EM reconstructions for MTREC

Experimentally derived structures for Mtl1 are not available, although it is predicted to include the classic domain architecture of Ski2 family helicases with N-terminal RecA domains that bind and hydrolyze ATP, the arch domain that includes the KOW domain which mediates protein-protein interactions, and the helical bundle (HB) domain 4 (Fig. 2a) ^16^. Less is known about Red1 as it contains few conserved domains with predicted functions ^41^. The C-terminus of Red1 encodes the C3H1 zinc finger motif (ZnF) which was shown to be necessary for Mtl1 binding (Fig. 2a), consistent with a recent structure of the *Chaetomium thermophilum* ZnF from Red1 fused to a fragment of *Chaetomium thermophilum* Mtr4 ^59^. Binding interfaces have also been elaborated between Red1, Pla1, Iss10, and Ars2 ^59,60,61^. Red1 was proposed to form a dimer mediated by a coiled-coil domain (residues 472-519) as evidenced by yeast two-hybrid and AlphaFold predictions using the sequence of the putative coiled-coil ^60^. However, to date there has been no biochemical or structural evidence that such a dimer forms in the context of MTREC. While these data highlight several domains relevant for Red1 function, its interaction with fission yeast Mtl1 has not been structurally characterized ^59,60^.

**Figure 2.**
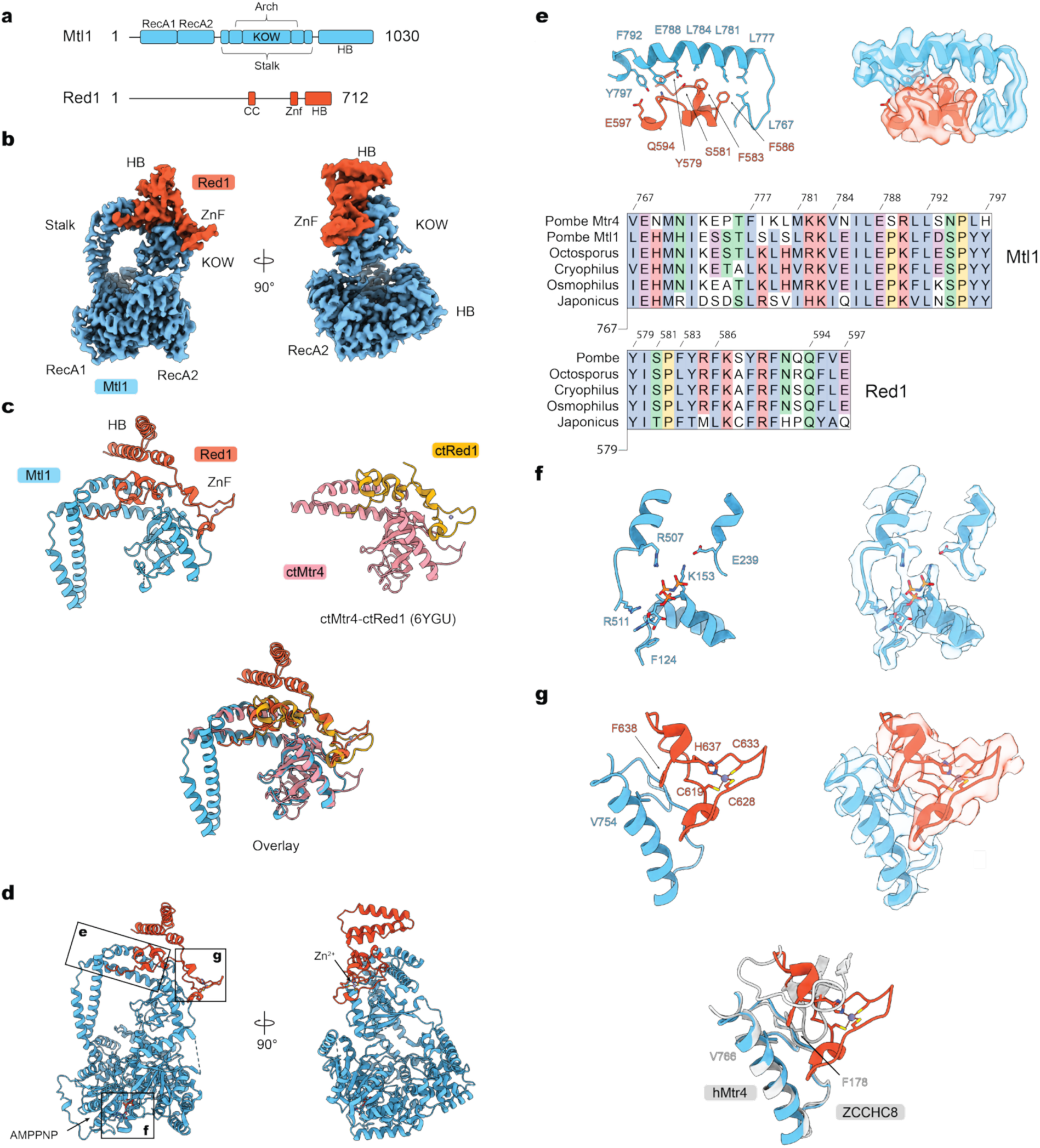
Cryo-EM structure of the MTREC Mtl1-Red1 ZnF-HB monomer. **(a)** Domain schematics of Mtl1 and Red1. Proteins are colored and labeled. **(b)** EM density map of the MTREC monomer structure highlighting conserved structural domains in Mtl1 and MTREC (contour at σ = 6.07). **(c)** Comparison of MTREC structure with the crystal structure of the ctMtr4 KOW domain with the ZnF domain of ctRed1 (PDB:6YGU). **(d)** Ribbon depiction of the atomic model of the MTREC monomer. **(e)** Magnified view of the residues in the hydrophobic channel mediating the interaction between Mtl1 and Red1 (contour at σ = 6.07). The sequence alignment shows the conservation of hydrophobic amino acids across fission yeast. **(f)** Ribbon and EM density map of the ATP binding site between the RecA domains. Density is observed for the *γ*-phosphate of the AMPPNP as well as key catalytic residues Lys153 and Glu239 (contour at σ = 6.07). **(g)** View of the Red1 zinc finger interaction with the KOW domain of Mtl1 (contour at σ = 6.07). The zinc finger positions a phenylalanine (638) proximal to a conserved valine (754) in Mtl1, mimicking a similar interaction observed between human MTR4 and ZCCHC8 in the NEXT complex as indicated in the superposed structures (below; PDB 7S7B).

To determine architectures for MTREC and interactions between Mtl1 and Red1 that could explain biochemical differences observed thus far, three structures were determined by single particle cryo-electron microscopy (Fig. S3, Table S3). In the first, data were collected on grids using reconstituted MTREC complex in a slight excess (+0.5 µM) of the 16 bp RNA duplex with a 10 nt, 3’ poly(U) overhang, magnesium and AMPPNP which resulted in a reconstruction with an overall resolution of 3.28 Å (Fig. 2b). Reconstructions and modeling revealed the architecture for Mtl1 which adopts the hook-like structure characteristic of Ski2 family helicases in addition to densities assigned to the C-terminal domains of Red1 (residues 570-712), including the conserved zinc finger domain that was previously characterized and a three-helix bundle located above the ZnF domain we annotate as the ZnF-HB domain (Fig. 2b). RNA was not observed in reconstructions, nor were the N-terminal residues of Mtl1 (1-85) or other residues of Red1. This sample was prepared in HEPES buffer (20 mM pH 7.5) which differed from the buffer used to purify the complex, so another attempt was made to determine reconstructions of MTREC in the purification conditions with Tris pH 8, using RNA substrate 1 in a equimolar concentrations with MTREC in the presence of AMPPNP and magnesium. While particles containing RNA were not observed, reconstructions revealed two distinct classes that correspond to MTREC complexes bound to the dimeric Red1 coiled-coil, one with a single Mtl1 molecule bound to the Red1 dimeric coiled-coil and ZnF-HB domains (monomer-CC) and the other for a 2:2 MTREC dimer with two molecules of Mtl1 bound to the dimeric coiled-coil and respective ZnF-HB domains (Fig. 3). Consistent with dimerization, gel filtration profiles of the purified MTREC show a retention volume that eluted much earlier (122 ml) relative to Mtl1 (212 ml), suggesting that MTREC forms a higher order oligomer relative to Mtl1 (Fig. S1b, S1c).

**Figure 3.**
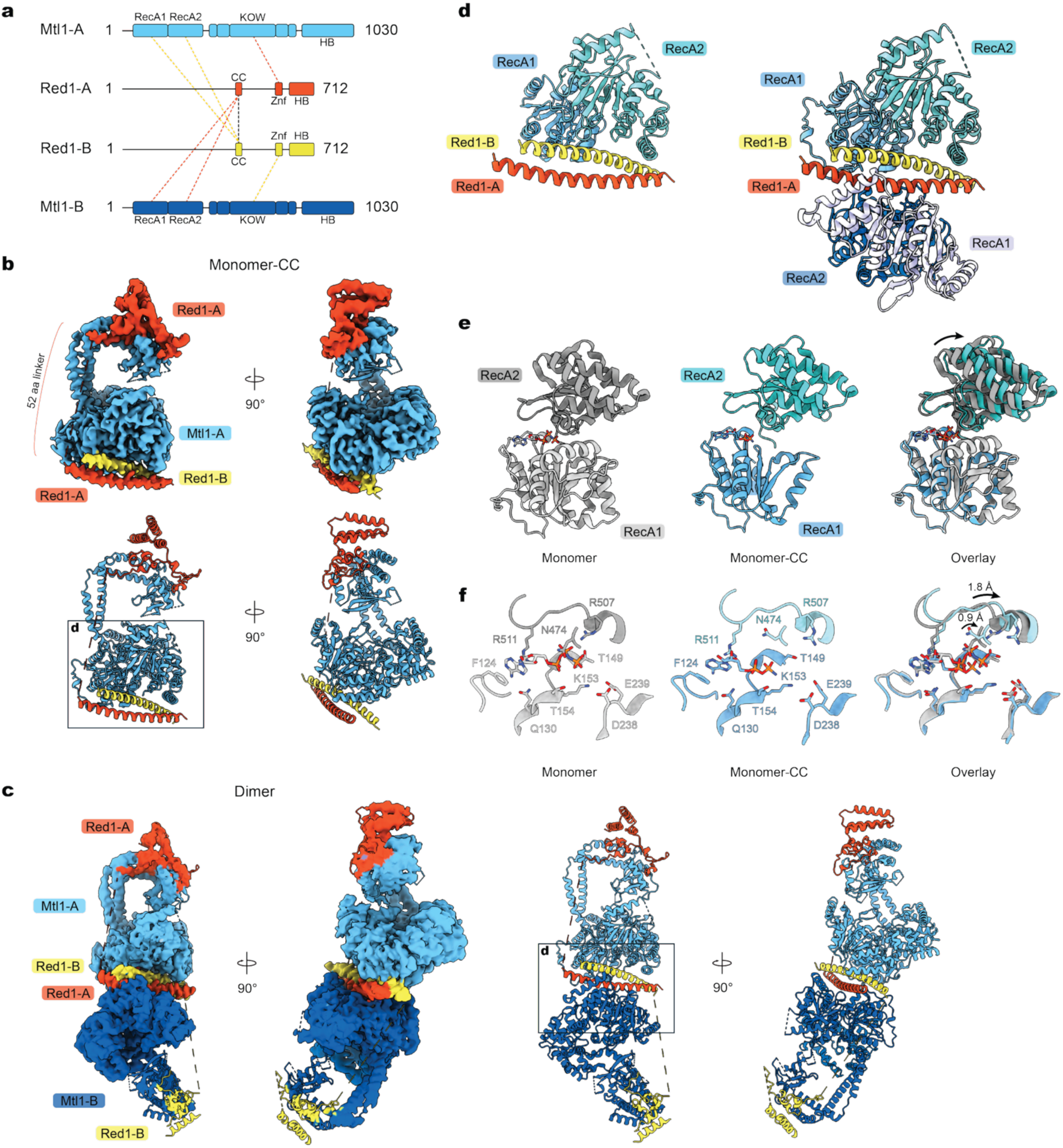
Cryo-EM structures of the MTREC monomer-CC and dimer. **(a)** Domain schematics of the MTREC dimer. Dashed lines indicate the organization of the observed interactions between subunits along with the putative assignment for connectivity. Red lines indicate contacts made by the Red1-A protomer and yellow lines indicate contacts made by Red1-B protomer. The black line indicates dimerization between Red1s mediated by the coiled-coil. **(b)** EM density map and ribbon depiction of the MTREC monomer-CC structure (contour at σ = 7.73). **(c)** EM density map and ribbon depiction of the MTREC dimer structure (contour at σ = 5.48). **(d)** Comparison of the monomer-CC (left) and dimer (right) structures shows that the RecA conformation is symmetric across the axis of the Red1 coiled-coil. **(e)** Aligning RecA1 domains of monomer-CC and monomer structures shows that the RecA2 domain from monomer-CC is shifted away from RecA1. **(f)** Schematic of key residues in the ATP binding site for monomer and monomer-CC and overlay showing that RecA1 and RecA2 move apart by 1-2 Å.

### Reconstruction of a Mtl1-Red1 ZnF-HB monomer complex

The reconstruction for the Mtl1-Red1 ZnF-HB monomer complex was used to build an initial model given its higher quality and resolution. In this reconstruction details for the ZnF-HB domain are clear and show that the C-terminus of Red1 is coordinated by the Mtl1 arch domain through extensive contacts. Interactions between the zinc finger and Mtl1 arch are similar to those reported for a crystal structure of a ctMtr4 fragment (residues 654-865) from *Chaetomium thermophilum* fused to a peptide (residues 1014-1091) from the ctRed1 homolog ^59^ (Fig. 2c). A hydrophobic channel in Mtl1 spanning residues 767-797 is conserved between other species of fission yeast, forming the primary interaction surface for a cluster of aromatic residues in Red1 (Fig. 2d,e) that bury a total surface area of 782 Å^2^ as calculated by UCSF Chimera^62,63^.

Densities are apparent for the C3H1 zinc finger which straddles the face of the KOW domain. Immediately following the zinc finger is a phenylalanine (F638) which extends down towards the KOW to form interactions with V766, an interaction that mimics the canonical arch-interacting motif (AIM) found in several Mtr4-binding proteins including the NEXT subunit ZCCHC8 ^13,35,64–68^, suggesting a convergent strategy for protein interactions with Mtr4 and Mtl1 helicases (Fig. 2d,g). As noted above, Red1 includes a C-terminal 3-helix bundle that stacks on top of the zinc finger that appears conserved in higher eukaryotes including humans as this 3-helix bundle is predicted in Red1 homologs and in AlphaFold models for human ZFC3H1 (accession ID: AF-O60293-F1) ^69–71^.

The Mtl1 ATPase active site appears intact compared to available structures of Mtr4 (PDB 2XGJ) ^72^. Densities are present for the AMPPNP nucleotide in the ATP binding cleft between the RecA1 and RecA2 domains of Mtl1 (Fig. 2f) along with side chains of Phe124 and Arg511 π-stacking above and below the purine ring of the AMPPNP and while densities for the three phosphates are observed surrounded by side chains of Lys153, Glu239 and Arg507 which are implicated in catalysis, magnesium is not visible. We hoped the Mtl1 structure might help to uncover interfaces within Mtl1 that could explain its higher intrinsic strand displacement activity relative to Mtr4, but no structural differences or amino acid substitutions within the Mtl1 RecA domain interfaces, predicted RNA binding sites or ATPase active site could be identified as different as they are all conserved between fission yeast Mtr4 and Mtl1.

### The MTREC dimer is mediated by the coiled-coil domain of Red1

The monomer-CC structure resembles the structure of the monomer Mtl1-Red1 (ZnF-HB) complex with respect to ZnF-HB interactions, but additional densities consistent with the Red1 antiparallel coiled-coil are present at the base of the helicase core (Fig. 3b, Fig. S3, Table S3). The MTREC dimer structure features two Mtl1 proteins arranged “tail-to-tail” with the coiled-coil buried between the RecA domains of each Mtl1 protomer (Fig. 3c). The helicase cores of the Mtl1 protomers are symmetric via a plane running through the coiled-coil (Fig. 3d). While this portion of the dimer is symmetric, the overall dimer structure appears asymmetric due to disorder and differences in the orientations of the respective arch domains (Fig. 3c). The monomer-CC and dimer structures are highly similar within respect to interactions between the coiled-coil and RecA domains so subsequent discussion of this interface will not distinguish between these two reconstructions.

Observed densities for the antiparallel Red1 coiled-coil include residues 476-521 and match the length previously predicted for the Red1 coiled-coil ^59^. Densities were not observed for residues 522-575 that span between the Red1 coiled-coil and ZnF-HB domains. It is not formally possible to assign the Red1 protomer coil that is interacting with the core, but we tentatively assign the coiled-coil protomer 2 as being connected to the ZnF-HB domain of protomer 1 because the terminus of Red1 ZnF-HB domain from protomer 1 resides on the same side of the complex as terminus of the Red1 coil from protomer 2. Furthermore, the distance between termini is 25 Å shorter for this pair (75 Å versus 100 Å). Regardless, each coil within the coiled-coil contributes to interactions with each RecA domain of Mtl1 including contacts within the Mtl1 ATPase active site. The interface between a single Mtl1 protomer and the Red1 coiled-coil dimer buries a total surface area of 1091 Å^2^, calculated by UCSF Chimera ^62,63^.

Densities for the nucleotide within the Mtl1 ATP binding site are observed in both the monomer-CC and dimer structures, however it is less ordered in comparison to the monomer MTREC structure as densities are not apparent for the terminal phosphate of AMPPNP. Consistent with this, superposing RecA domains from monomer and monomer-CC structures shows that the RecA1 rotates away from RecA2 (Fig. 3e) which results in RecA1 residues in the ATP active site shifting away from RecA2 by approximately 1-2 Å (Fig. 3f). Disruption of the ATPase active site via interactions with the Red1 coiled-coil might explain why MTREC is less active in displacement assays relative to Mtl1.

### Red1 coiled-coil contacts to Mtl1 RecA domains modulate Mtl1 activities

As noted above, the Red1 coiled-coil interacts across the RecA domains of Mtl1, forming two distinct binding interfaces and anchoring the coiled-coil just below the ATP binding site (Fig. 4a). While slightly displaced from that observed for the MTREC Mtl1-Red1 ZnF-HB monomer, Mtl1 RecA1 Phe124 and RecA2 Arg511 form π-stacking interactions with the AMPPNP nucleobase. But in complexes with the Red1 coiled-coil, residues flanking Phe124 interact with the coil of Red1-A while residues flanking Arg511 interact with the coil of Red1-B (Fig. 4b). Specifically, RecA1 Pro123 contacts the methyl group of Red1-A Thr503, and the aliphatic portion of the Glu125 sidechain contacts Tyr510 through nonpolar interactions while its carboxylate group is positioned proximal to Lys513 (Fig. 4b). For the Red1-B coil, the aliphatic portion of the Lys494 side chain of Red1-B interacts through nonpolar contacts to Ile513 of RecA2 while two glutamates, Glu486 and Glu489, are positioned to make electrostatic interactions with Arg91 of RecA1. In addition, the Thr493 hydroxyl of Red1-B appears within hydrogen bonding distance of the N9 atom of the adenine.

**Figure 4.**
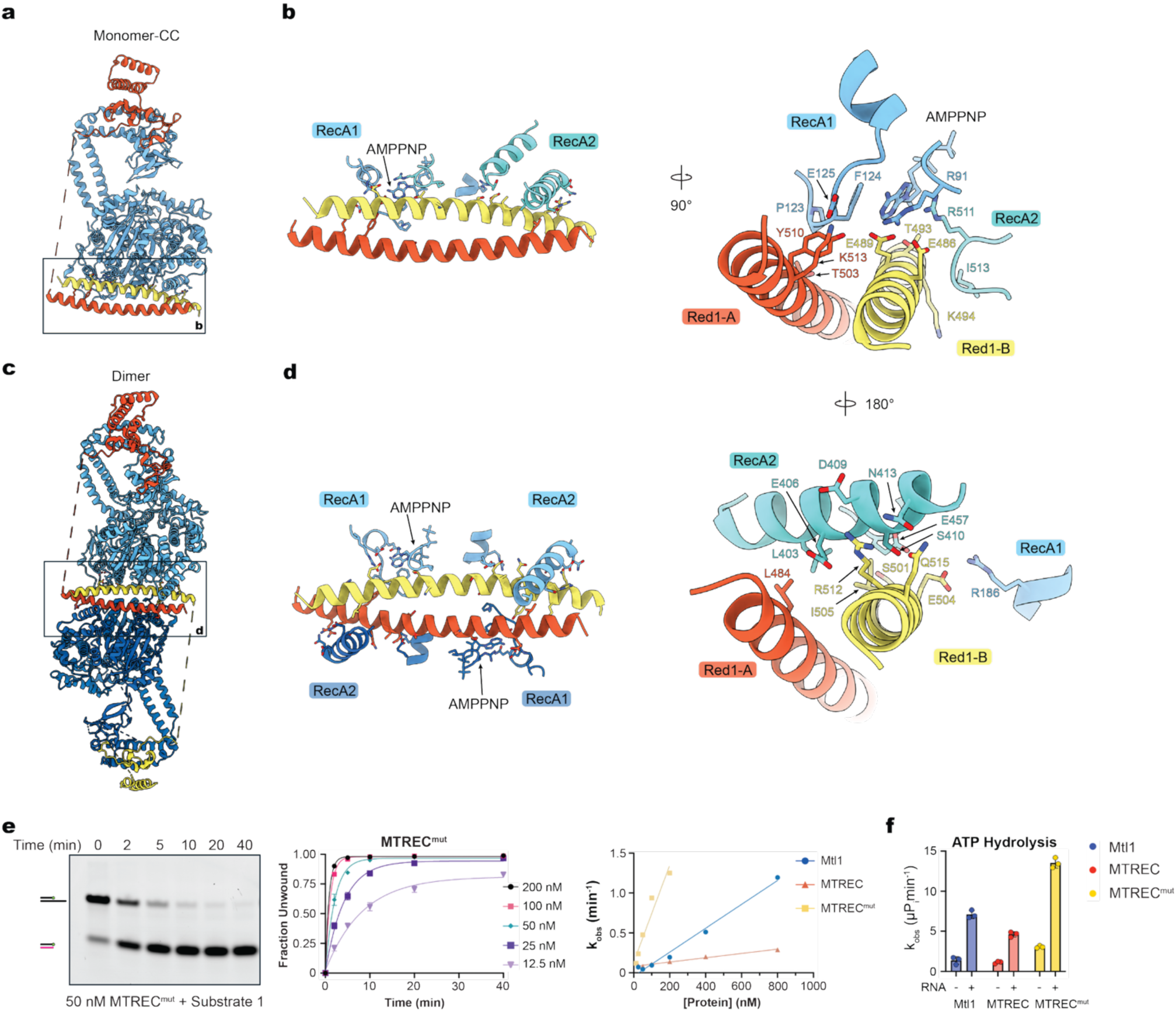
The Red1 coiled-coil contacts with the helicase core modulate Mtl1 activity. **(a)** Cartoon depiction of the MTREC monomer-CC structure. **(b)** Schematic of the Red1 coiled-coil interactions with Mtl1 (left) and detailed view with side chain interactions with the RecA1 and RecA2 domains of Mtl1 (right and bottom). Side chains and AMPPNP shown in stick representation superposed on ribbon cartoon representation of the backbone. **(c)** Cartoon depiction of the MTREC dimer structure with close-up **(d)** of Mtl1 RecA domains from both Mtl1 protomers to illustrate symmetrical interactions with the coiled-coil domain depicted as in (a). **(e)** Representative gel (left) for strand displacement shown next to graph of quantified data (middle) and graph of rates across relevant concentrations of MTREC^mut^, MTREC and Mtl1 for comparison (right). **(f)** Graph depicting rates obtained from ATP hydrolysis assays in the absence and presence of RNA for Mtl1, MTREC and MTREC^mut^. Biochemical data and quantification were obtained from experiments performed in triplicate and error bars represent the standard error of the mean (SEM) as calculated by GraphPad Prism.

A second major interface between the Red1 coiled-coil and Mtl1 primarily involves contacts between RecA2 and both Red1-A and Red1-B coils (Fig. 4b). Specifically, RecA2 Leu403 of Mtl1 contacts Leu484 of Red1-A via nonpolar interactions, Glu406 and Asp409 are positioned to form ionic interactions with the guanidinium of Arg512 while its aliphatic side chain forms nonpolar interactions with Ser410. Additional contacts also include hydrogen bonding contacts between the Mtl1 Asn413 side chain and Red1-B Gln515 side chain and the aliphatic portion of the Glu457 side chain contacting methyl group of Ser501 and side chain of Ile505 (Fig. 4b). One additional contact in this second interface is consistent with an ionic interaction between RecA1 domain Arg186 and Red1-B Glu504 side chains. Collectively, interactions between both Red1 protomers and Mtl1 RecA1 and RecA2 domains appear to contribute to an open configuration of the RecA domains that in turn influences the conformation of the nucleotide and residues within the Mtl1 ATPase active site (Fig. 3f). Interactions described between Red1 and Mtl1 for the monomer-CC structure are identical in the dimer structure (Fig. 4c,d).

Biochemical data presented thus far suggest that Red1 alters Mtl1 activities in the context of the MTREC core complex by enhancing binding affinity to RNA but impeding its ability to fully complete strand displacement relative to Mtl1 even at concentrations above its apparent K_D_. To determine if amino acid side chains in the interface between the Red1 coiled-coil and Mtl1 helicase domains contribute to this effect, mutants were constructed to remove the coiled-coil or generate mutations on its surface. Attempts to express a Red1 coiled-coil truncation mutant failed as this construct was poorly soluble yielding insufficient protein for biochemistry, but a mutant of Red1 that retained coiled-coil dimerization while replacing Red1 Glu486, Glu489, Thr493, Ser501, Gln508, Tyr510, Arg512, Gln515 residues with alanine could be expressed, purified and reconstituted similar to wild type MTREC (Fig. S1e,f). These residues contribute to interfacial contacts to Mtl1 as observed in our structures. Gel filtration profiles of the purified complex demonstrated a retention volume (122 mL) equivalent to wild type MTREC, indicating that oligomerization was preserved (Fig S1e). The resulting mutant MTREC^mut^ was used in strand displacement and ATP hydrolysis assays in comparison to MTREC and Mtl1 (Fig. 4e,f, Fig. S4, Tables S5, S6).

Strand displacement assays with the Mtl1-Red1^mut^ complex showed enhanced activity relative to both MTREC and Mtl1 (Fig. 4e, Fig. S4a, Table S5). Comparisons reveal that at 50 nM Mtl1 catalyzes displacement with an apparent K_obs_ = 0.05 ± 0.01 (nM)(min^−1^) while MTREC^mut^ reaches a rate almost 10-fold higher at K_obs_ = 0.48 ± 0.05 (nM)(min^−1^). Further, 50 nM Mtl1 has a half time of t_1/2_ = 15 min while MTREC^mut^ has a half time of t_1/2_ = 1.4 min (Fig. 4e, Table S5). It is also noteworthy that MTREC^mut^ achieved full strand displacement to levels comparable to Mtl1, suggesting that Mtl1-Red1 coiled coil interactions may also contribute to the reduced displacement capacity observed in assays with MTREC. Collectively, these data suggest that Red1 coiled-coil contacts to Mtl1 suppress strand displacement activity, perhaps through disruption of the RecA domains and ATPase active site. To test this directly, ATPase assays were conducted in the absence and presence of RNA at 1 µM concentration for Mtl1, MTREC and MTREC^mut^. ATPase activities were stimulated by RNA in each instance, but ATPase activities were highest for MTREC^mut^ and lowest for MTREC relative to Mtl1 (Fig. 4f, Fig. S4b, Table S6). These data are consistent with structural observations thus far. We posit that the rate enhancements observed for MTREC^mut^ in both strand displacement and ATPase assays is likely a combination of alleviating the inhibitory properties of the coiled-coil interactions with Mtl1 RecA domains coupled with the ability of Red1 to enhance RNA binding in the context of MTREC.

## DISCUSSION

Biochemical assays and structural characterization of reconstituted *S. pombe* Mtl1 and MTREC core complex and comparisons to Mtr4 reveal that Mtl1 is more active relative to MTREC or Mtr4, that MTREC binds RNA better than Mtl1, and that Red1 includes an autoinhibitory coiled-coil domain that dimerizes MTREC through contacts to the Mtl1 RecA domains to disrupt its ATPase active site as mutations predicted to disrupt interactions between the Red1 coiled-coil and Mtl1 result in enhanced ATPase and strand displacement activity.

Previous studies relegated Red1 to a scaffolding role due to a lack of predicted functional domains coupled with a role in mediating interactions between Mtl1 and MTREC associated subcomplexes ^15,16,31,59,60^.

However, our data reveals that MTREC displays properties unique amongst identified RNA exosome cofactors. While the ancillary subunits within TRAMP and NEXT complexes appear to stimulate translocation by Mtr4, Red1 appears to suppress strand displacement by Mtl1 within MTREC by inducing a shift in the RecA domains that would be predicted to inhibit ATP hydrolysis. These data suggest that Red1 may endow MTREC with the ability to bind RNA while slowing translocation so that it remains bound to RNA long enough to chaperone it to the RNA exosome for processing or degradation.

While activities differ, features are emerging that suggest some protein cofactors of Mtr4 and Mtl1 serve to modulate their activities to prevent or slow RNA translocation, all of which cover interfaces that are predicted to be required for interaction with the MPP6 cofactor for the nuclear RNA exosome, a cofactor that stimulates translocation and anchors the helicase to the RNA exosome core ^10,73,74^.For MTREC, the Red1 coiled-coil covers a similar interface between RecA domains that is observed interacting with ZCCHC8 in NEXT and MPP6 in a complex with the human nuclear exosome (Fig. 5a) ^10,13,35,38^. But unlike interactions observed in RNA-bound NEXT complexes that would preclude interactions with both MPP6 and the EXOSC2 exosome core subunit, the Red1 coiled-coil would not be predicted to interfere with docking onto the exosome core, though it would need to dissociate from the Mtl1 RecA domains if Mpp6 interacts similarly with Mtl1 (Fig. 5b).

**Figure 5.**
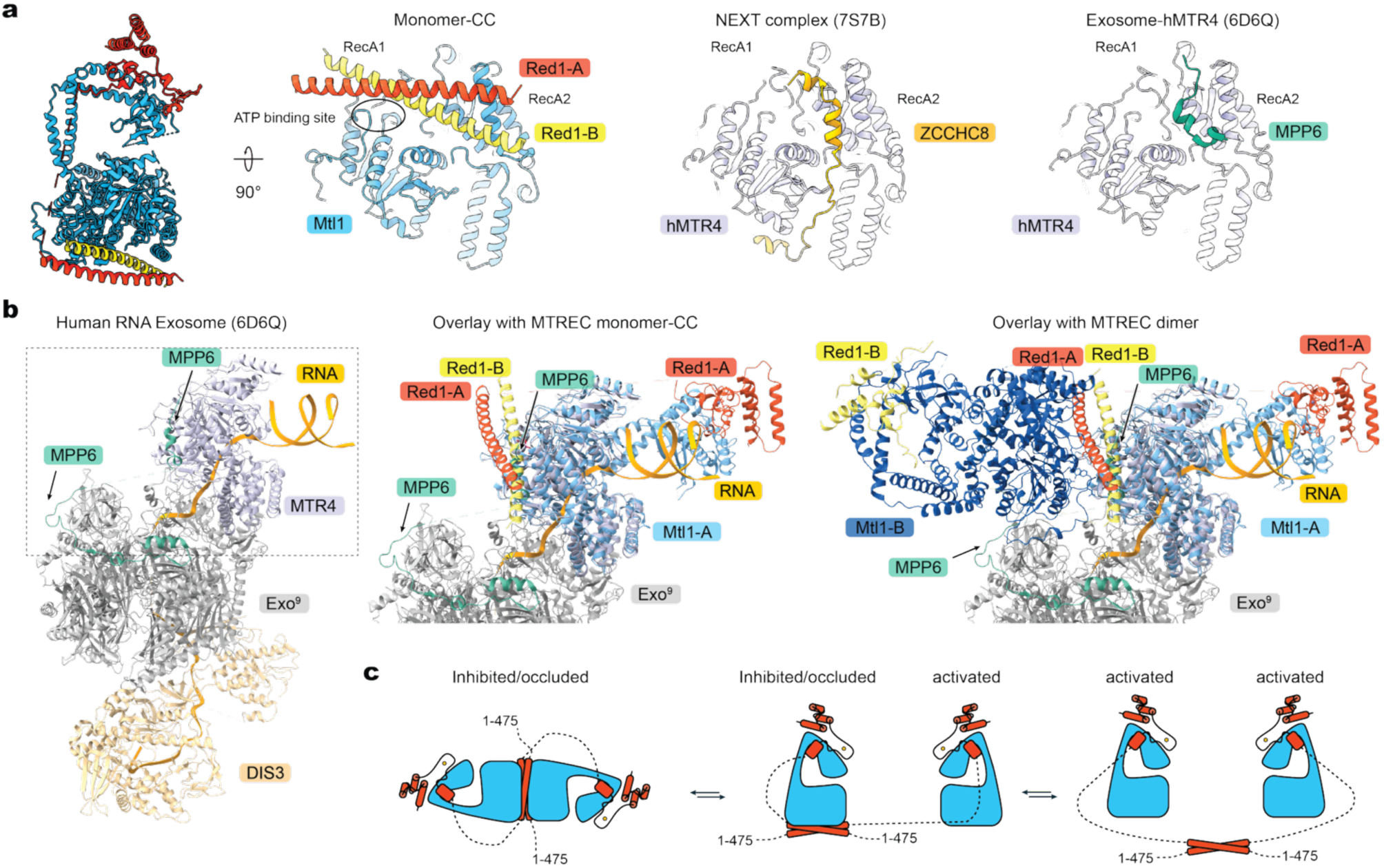
Comparison of MTREC with other Mtr4-containing complexes. **(a)** Comparison of the MTREC monomer-CC structure to the structure of the human NEXT (PDB 7S7B) and of MTR4 bound to MPP6, the cofactor that bridges Mtr4 to the RNA exosome (PDB 6D6Q). Both the C-terminal domain of ZCCHC8 in the NEXT complex and the N-terminal domain of exosome cofactor MPP6 make contacts with the RecA2 domain of MTR4. Red1 contacts an overlapping surface in RecA2 but extends toward RecA1 proximal to the ATP binding site. **(b)** Structure of the human MTR4-MPP6-RNA exosome structure (left). Alignment of the MTREC monomer-CC (middle) and MTREC dimer (right) to MTR4 in the exosome structure to illustrate that the Red1 coiled-coil would sterically occlude MPP6 interaction, but would not sterically occlude interaction with the exosome core. **(c)** Schematic of possible MTREC dimer configurations.

MTREC appears most like the PAXT-like complexes in humans due in part to similarities between Red1 and the C3H1 zinc-finger protein ZFC3H1 along with the ability of these core complexes to interact with nascent RNP cofactors that bind to the RNA cap and poly(A) tails ^15,16,18,59,60,75^. Structures characterizing binding interfaces between Red1 and MTREC associated proteins have shed some light on how Red1 connects Mtl1 to various subcomplexes, but further structural and biochemical data will be required to investigate interactions between intact MTREC, RNA and these RNA binding cofactors, especially as we were unable to observe interactions with RNA in our cryo-EM reconstructions. It is also interesting to note that NEXT and MTREC form dimers (Fig. 5c). While it remains unclear if dimerization is important for orchestrating RNA capture or delivery to the exosome, dimerization interfaces via ZCCHC8 in NEXT and Red1 in MTREC modulate helicase activities while occluding surfaces required for interactions with the nuclear exosome or its cofactors. Details differ between NEXT and MTREC, but both present plausible hierarchical strategies to modulate activities and orchestrate remodeling of protein surfaces to coordinate RNA capture, delivery and translocation.

## METHODS

### Cloning

Codon optimized genes for Mtl1 and Red1 were obtained from Life Tech, and Red1^mut^ was obtained from Genscript. DNA encoding Mtl1 was inserted into a pSmt3 vector between the BamHI and SalI sites, resulting in a his_6_Smt3 tag N-terminal to Mtl1. The coding region of Red1 and Red1^mut^ was inserted into a modified pET-28(b) vector encoding an N-terminal his_6_GST tag between the BamHI and SalI sites.

### Protein Expression and Purification

*S. pombe* Mtr4 was purified as described previously^10^. pSmt3-Mtl1 was transformed into SoluBL21 cells (Genlantis) and cultures grown to an OD_600_ of ∼1.2. Before induction, cultures were cooled in an ice bath for 15 minutes and supplemented with 0.2% ethanol to induce chaperone expression. Gene expression was induced by the addition of 0.4 mM isopropyl *β*-D-1-thiogalactopyranoside (IPTG) and cultures were grown overnight at 18°C for 18 hours. Cells were pelleted by centrifugation and suspended in lysis buffer (50 mM Tris pH 8, 500 mM NaCl, 20% w/v sucrose, 0.1% v/v IGEPAL, 1 mM DTT), flash frozen in liquid nitrogen and stored at −80°C.

Frozen cell pellets were thawed and then lysed by sonication for 20 minutes at 20% amplitude and centrifuged at 16,000 g for 20 min to pellet cell debris. The supernatant was removed and then applied to Ni^2+^-NTA resin (Qiagen) on a gravity column. The resin and associated proteins were washed with wash buffer A (20 mM Tris pH 8, 500 mM NaCl, 25 mM imidazole, 1 mM DTT) and chaperones were removed by an ATP wash (wash buffer plus 2 mM ATP and 2.5 mM MgCl_2_). His_6_Smt3-Mtl1 was eluted in wash buffer that included 250 mM imidazole, concentrated in a 50K MWCO Amicon to a volume <10 mL, and then dialyzed into size exclusion chromatography (SEC) buffer (20 mM Tris pH 8, 350 mM NaCl, 1 mM TCEP) overnight at 4°C in the presence of the Ulp1 protease (20 ng/µL) to remove the His_6_Smt3 solubility tag from Mtl1 ^76^.

The protein mixture was separated by size exclusion chromatography using a HiLoad S200 26/600 size exclusion column (GE) and the peak corresponding to Mtl1 was identified by SDS-PAGE analysis. Mtl1 was then diluted in buffer A (20 mM Tris pH 8, 1 mM TCEP) to a final ionic strength of 87.5 mM NaCl before application to a HiTrap Heparin column to remove contaminating nucleic acids. Mtl1 was eluted with a linear salt gradient from 87.5 mM to 1M NaCl over 15 column volumes (Fig. S1). The resulting peak that included Mtl1 was buffer exchanged into SEC buffer with a 50K MWCO Amicon (Sigma-Aldrich), concentrated, and flash frozen in liquid nitrogen for storage at −80°C.

The pET28b(+)-his_6_GST-Red1 plasmid was transformed into Arctic Express cells (Agilent). Cells were grown, induced for gene expression, isolated, prepared and lysed as described for cells expressing Mtl1. Clarified lysate was applied to glutathione resin (Cytiva) and washed with wash buffer B (20 mM Tris pH 8, 500 mM NaCl, 1 mM DTT). Mtl1-expressing SoluBL21 cells were lysed and clarified by centrifugation as described above and the supernatant was then applied to the glutathione resin and allowed to incubate for 10 minutes. Proteins that remained bound to the resin were washed with wash buffer B and chaperones were removed with an ATP wash (wash buffer supplemented with 2 mM ATP and 2.5 mM MgCl_2_). Proteins were then eluted from the resin using wash buffer B supplemented with 100 mM reduced glutathione over 4 column volumes. Fractions containing the majority of protein as determined by SDS-PAGE were combined and concentrated to <10 mL with a 100K MWCO Amicon and dialyzed overnight into SEC buffer in the presence of 20 ng/µL Ulp1 and 10 ng/µL TEV protease to remove the his_6_Smt3 and his_6_GST tags from Mtl1 and Red1 respectively. After dialysis and tag cleavage, MTREC was purified via gel filtration and a heparin column as described above for Mtl1, concentrated to 24 mg/ml, and snap frozen in liquid nitrogen for storage at −80°C (Fig. S1).

The pET28b(+)-his_6_GST-Red1^mut^ plasmid was transformed into Arctic Express cells (Agilent). Cells were grown, induced for protein expression, isolated, prepared and lysed as described for cells expressing Red1. Purification of MTREC^mut^ proceeded as described for the wild type MTREC complex (Fig. S1).

### Preparation of RNA Substrates for Biochemical Assays

RNA oligos (Table S4) were obtained from Integrated DNA Technologies (IDT) as lyophilized powder after HPLC-purification. Extinction coefficients for single stranded oligos and duplex substrates were estimated using the IDT Oligo Analyzer tool ^77^, and sample concentrations were determined by measuring absorbance at 260 nm with a spectrophotometer (Nanodrop 2000). Duplex substrates were obtained by mixing complementary strands at equimolar concentrations, heating to 95°C for three minutes, and passive cooling to room temperature.

### Electrophoretic Mobility Shift Assays (EMSAs)

Assays were performed in a buffer containing 20 mM HEPES pH 7.5, 140 mM NaCl, 1 mM TCEP, 1 U/µL murine RNase inhibitor (New England Biolabs), 5% glycerol (v/v). For 50 µL assay volumes, 10 nM 6-carboxyfluorescein (FAM) labeled RNA substrates were added with protein to 45 µL. Assays were supplemented with 5 µL of buffer containing 20 mM AMPPNP and 20 mM MgCl_2_. The resulting mixtures were incubated on ice for 20 minutes. Then, aliquots of each assay condition were analyzed by applying samples to a 4-20% Novex TBE gel (ThermoFisher) in pre-cooled 1x TBE at 4°C.

Gel image files were linearized before processing in ImageJ (https://imagej.nih.gov/ij/plugins/linearize-gel-data.html) ^78^. Band intensities were measured by selecting lanes and integrating the area under the plotted curve. The fraction of RNA duplex bound was calculated by dividing the intensity of the free RNA band for each lane by the intensity of the free RNA band in the absence of protein and subtracting from 1 (Fig. S2, Table S1). Values for fraction bound at each protein concentration were plotted in GraphPad Prism and curves were fit to a quadratic binding model

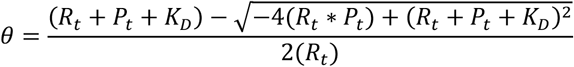

where *θ* is fraction bound, *R_t_* is the concentration of RNA, and *P_t_* is the concentration of protein to account for a potential decrease in the concentration of free protein ^79,80^.

### Strand Displacement Assays

Strand displacement assays were performed as described previously ^13^. Assays were performed in a buffer containing 20 mM Bis-Tris pH 6.5, 40 mM NaCl, 5 mM *β*-mercaptoethanol (BME), 1 U/µL murine RNase inhibitor (New England Biolabs), 5% glycerol (v/v), and 0.5 mM MgCl_2_. For assays at volumes of 50 µL, 10 nM 6-carboxyfluorescein (FAM) labeled RNA substrates were added with protein to 45 µL and equilibrated at 30°C for 5 minutes. Reactions were initiated by adding 5 µL of start buffer (20 mM ATP, 20 mM MgCl_2_, 4 µM DNA trap oligo) pre-warmed to 30°C.

5 µL aliquots were removed from each reaction at specified time points and reactions stopped by addition of 2X quench buffer (10 mM EDTA, 1% SDS (w/v), 10% glycerol, 0.00125% xylene cyanol (w/v), 80 U/mL Proteinase K (New England Biolabs)). Samples were then incubated at room temperature for an hour to digest protein before separation by native gel electrophoresis using 20% Novex TBE gels (ThermoFisher) in 1x TBE buffer pre-cooled to 4°C and at a voltage of 200V for 40 minutes. Gels were imaged for FAM fluorescence using a Typhoon laser scanner (473 nm 350 V with an LBP 510LP filter).

Gel image files were linearized before processing in ImageJ (https://imagej.nih.gov/ij/plugins/linearize-gel-data.html) ^78^. Gels were analyzed and band intensities quantified as described for EMSA assays. The fraction of RNA duplex unwound or *θ*, was calculated by dividing the intensity of the free RNA band at time t by the intensity of the free RNA band at t = 0. Values obtained for the fraction unwound at each time point were plotted in GraphPad Prism and curves were fit to a one phase association model to determine K_obs_ (Fig. S2, Table S2).

### Cryo-EM sample and grid preparation

Purified MTREC protein and RNA substrates were prepared as described in earlier sections. For the MTREC monomer structure for which the Mtl1 helicase and ZnF-HB domains of Red1 were visualized, MTREC and FAM-labeled RNA duplex with a 10 nt poly(U) overhang were mixed in buffer containing 20 mM HEPES pH 7.5, 140 mM NaCl, 1 mM TCEP, 2 mM MgCl_2_, and 2 mM AMPPNP at concentrations of 10 μM and 10.5 μM, respectively. Reactions were incubated on ice for 30 minutes and then supplemented with 0.003% (w/v) cetyltrimethylammonium bromide (CTAB) before vitrification. For the MTREC monomer-CC and dimer structure where the Mtl1 helicase, ZnF-HB and coiled-coil portions of Red1 were visualized, MTREC and FAM-labeled RNA duplex with a 25 nt poly(A) overhang were mixed in a buffer of 20 mM Tris pH 8, 150 mM NaCl, 1 mM TCEP, 2.5 mM MgCl_2_, and 1 mM AMPPNP at concentrations of 2.5 μM and 3 μM, respectively. Reactions were incubated on ice for 30 minutes and then supplemented with 0.05% (CHAPSO) before vitrification. For the monomer and monomer-CC/dimer samples, 4 µL of sample was spotted onto glow-discharged UltraAUFoil 300 mesh R1.2/1.3 grids (Quantifoil). After 8 seconds, grids were blotted for 3.5 seconds at 100% humidity and plunged into liquid ethane using an FEI Vitrobot Mark IV.

### Cryo-EM data collection

Cryo-EM data were collected using a 300 kV Titan Krios transmission electron microscope (FEI-ThermoFisher) at the MSK Richard Rifkind Center for Cryo-EM. Movies (40 frames per movie, 4 s exposure time) were recorded at a dose rate of ∼20 e^−^ pixel^−1^s^−1^ using a K3 Summit direct electron detector (Gatan) in super-resolution mode at a physical pixel size of 1.064 Å and a dose of 66 e^−^/ Å^2^/movie. Automated collection was performed in Serial EM using image shift to record data from 9 holes per stage movement (Table S3).

### Cryo-EM image processing

Image processing was performed in CryoSPARC unless otherwise indicated ^81^. Movies were gain normalized, 2x Fourier cropped, dose-weighted, and corrected for drift using patch motion correction. Estimation of the contrast transfer function (CTF) was done using CryoSPARC’s patch CTF estimation job. Micrographs with an estimated resolution of worse than 4.5 Å were discarded and remaining images used for particle picking. A circular blob picker was used on a subset of images to generate 2D classes for template picking. Particles identified by template picking were extracted with a box size of 384 pixels and subjected to multiple rounds of 2D and 3D classification to remove junk classes.

### MTREC monomer structure data processing

6,754 movies were used to obtain template picked particles which were culled through 2D classification, and a subset of 1,494,063 particles were used to generate volumes for heterogeneous refinement with ten classes which resulted in four classes that appeared to include good particles. 855,836 particles from these four classes were combined and subjected to additional rounds of 2D classification to further remove junk classes which resulted in 172,912 particles.

2D classes were balanced for particle distribution before using them as training data for CryoSPARC’s implementation of Topaz neural network-based particle picking ^82^. Trained models were applied to the full dataset to generate new particle stacks which were extracted with a box size of 384 pixels resulting in 975,644 particles. The Topaz-picked particles were used to generate 12 volumes via *ab initio* reconstruction and then pruned over the course of 11 rounds of heterogeneous refinement to select the highest quality particles which resulted in 377,130 particles.

This particle stack was then used for homogeneous and non-uniform refinement. The metadata for the final stack of 375,158 particles was converted to the RELION3.1 format using UCSF pyem ^83,84^. The original micrographs were gain normalized, dose-weighted, and binned by a factor of 2, and corrected for particle drift using MotionCor2 ^85^. CTF estimation was done using Gctf on dose-weighted, corrected micrographs ^86^.

The reconstructed volume from the final non-uniform refinement CryoSPARC job was used to generate a volume at an absolute intensity greyscale for use in RELION3.1 ^83,87^. A reference structure for Bayesian polishing was generated by applying a mask to the helicase core and performing another round of 3D auto-refinement to generate a focused refinement for the arch domain.

The final stack of 375,158 particles was iterated through 2 rounds of Bayesian polishing ^88^. Particles were further segregated by 3D classification without image alignment to separate different conformations of the helicase core, arch, and KOW domains. Particles from the same class of conformers were combined and used for 3D auto-refinement to generate final reconstructions, which were then combined to generate a composite map (Fig. S3, Table S3). The composite map was used to build an atomic structure using Coot based on prior structures and AlphaFold models (accession IDs: AF-O13799-F1-v6, AF-Q9UTR8-F1-v6), and refined using Phenix ^69,89–91^.

### MTREC monomer-CC and dimer structure data processing

2,309 movies were used to identify 1,170,338 template picked particles that were further segregated by 2D classification to remove junk classes. The resulting 425,869 particles were used as input for *ab initio* reconstructions to generate templates for heterogeneous refinement. The particle stack was iterated through heterogeneous refinement to remove junk particles, yielding a final stack of 176,300 particles. Two distinct classes were observed within this dataset which were designated monomer-CC as it included one molecule of Mtl1 and the ZnF-HB and the coiled-coil dimer from Red1 and dimer as it included two molecules of Mtl1, two copies of the ZnF-HB domain as well as the coiled-coil dimer of Red1. These particles were used as input for ab initio reconstructions to generate models of each of the two classes observed in this dataset.

Monomer-CC and dimer reconstructions were used to train a Topaz autopicker which resulted in 1,618,019 particles that were pruned to 844,548 particles via 2D classification to remove junk particles. Using reconstructions from the previous ab initio job, particles were iterated through rounds of heterogeneous refinement to separate monomer and dimer classes (Fig. S3).

Particles corresponding to the monomer structure were discarded as they resulted in reconstructions that were worse than previously obtained, and 90,218 particles corresponding to the monomer-CC class were selected and re-extracted. These particles were used to generate an ab initio reconstruction, and duplicate particles were removed. The ab initio reconstruction was refined using non-uniform refinement and subsequently used in CryoSPARC’s reference-based motion correction to obtain a reconstruction through homogeneous refinement. A mask was created that encompassed the Red1 coiled-coil and Mtl1 RecA domains and used for analysis with 3D classification without image alignment to segregate particles with and without the coiled-coil domains. Similar classes were combined to obtain a stack of 69,515 particles which were used to generate a new reconstruction using non-uniform refinement. Additional masks were generated encompassing the helicase core and the arch domains for use in local refinement jobs to obtain higher resolution reconstructions for each region. The refined reconstructions were combined to generate a composite monomer-CC map in Phenix which was then used to manually build and refine atomic models based on docking of the monomer MTREC and coiled-coil AlphaFold models using Coot and Phenix, respectively (Table S3) ^89,90^.

103,503 particles corresponding to the dimer class were re-extracted and used to generate an ab initio reconstruction. Duplicate particles were removed, resulting in 73,236 particles. The resulting reconstruction was used in CryoSPARC for reference-based motion correction, and particles output from that job were used to generate an *ab initio* reconstruction followed by non-uniform refinement. Masks were created encompassing the dimer interface which included the Red1 coiled-coil and Mtl1 RecA domains as well as two masks covering each of the two Mtl1 protomers. Masks were used for focused refinement, and resulting maps were combined to generate a composite map which was then used to manually build and refine atomic models based on docking of the monomer-CC MTREC models using Coot and Phenix, respectively (Table S3) ^89,90^.

## DATA AVAILABILITY

EM reconstructions and atomic coordinates are deposited in the Electron Microscopy Data Bank (EMDB) and Protein Data Bank (PDB) with the following accession codes: Model and composite map for *S. pombe* Mtl1-Red1 monomer bound to AMPPNP, EMD-76234, PDB ID 11ZY; Focused reconstruction for Mtl1-Red1 MTREC monomer RecA domains bound to AMPPNP, EMD-76236; Focused reconstruction for Mtl1-Red1 monomer stalk/KOW domains, EMD-76237; Focused reconstruction for Mtl1-Red1 monomer KOW domain, EMD-76238; Model and composite map for *S. pombe* Mtl1-Red1 dimer with Red1 coiled-coil dimer bound to AMPPNP, EMD-76239, PDB ID 11ZY; Overall reconstruction for Mtl1-Red1 monomer-H2 with Red1 coil-coil dimer bound to AMPPNP, EMD-76240; Focused reconstruction for Mtl1-Red1 monomer-H2 RecA domains with Red1 coil-coil dimer bound to AMPPNP, EMD-76241; Focused reconstruction for Mtl1-Red1 monomer-H2 Stalk/KOW domains with Red1 coil-coil dimer bound to AMPPNP, EMD-76242; Model and composite map for *S. pombe* Mtl1-Red1 dimer with Red1 coiled-coil bound to AMPPNP, EMD-76243, PDB ID 11ZY; Overall reconstruction for Mtl1-Red1 dimer with Red1 coiled-coil bound to AMPPNP, EMD-76244; Focused reconstruction for Mtl1-Red1 dimer RecA domains with Red1 coiled-coil bound to AMPPNP, EMD-76245; Focused reconstruction for Mtl1-Red1 dimer protomer A with Red1 coiled-coil bound to AMPPNP, EMD-76246; Focused reconstruction for Mtl1-Red1 dimer protomer B with Red1 coiled-coil bound to AMPPNP, EMD-76247, and will be released upon review by RCSB staff.

## ACKNOWLEDGEMENTS

We thank members of the Lima lab for advice. We thank Jason De La Cruz and Sagnik Sen for assistance during cryo-EM data collection.

## AUTHOR CONTRIBUTIONS

**Lucas D. Repeta:** Conceptualization (co-lead); Investigation (lead); Writing – original draft (lead).

**Christopher D. Lima:** Conceptualization (co-lead); Investigation (advised); Writing – editing.

