## Supporting Information for "Architectures and biochemical activities of Mtl1-Red1 MTREC helicase complexes"

**This PDF file includes:**

Figures S1 to S4
Tables S1 to S6

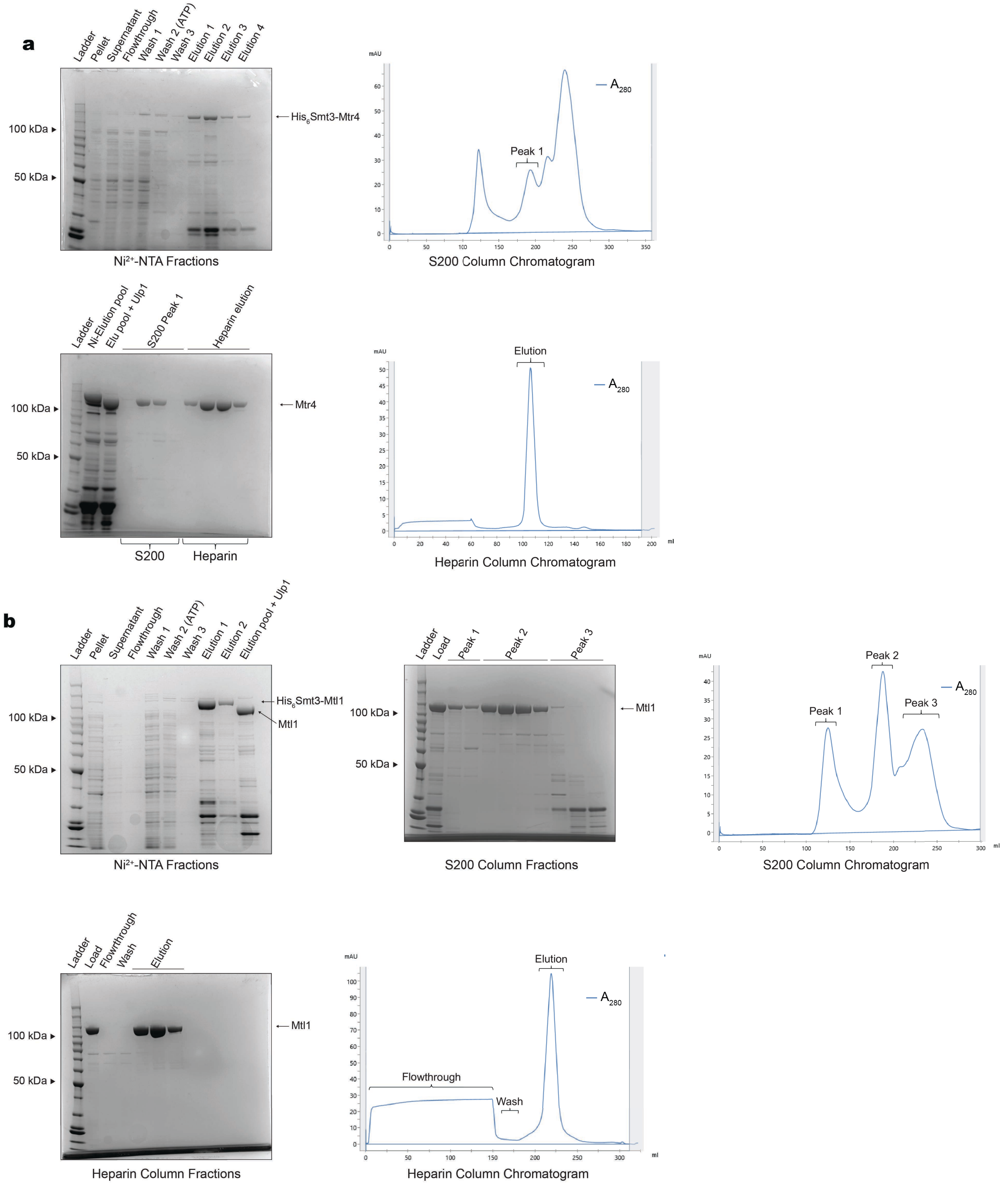
a
Ladder
Pellet
Supernatant
Flowthrough
Wash 1
Wash 2 (ATP)
Wash 3
Elution 1
Elution 2
Elution 3
Elution 4
100 kDa
50 kDa
His6Smt3-Mtr4
Ni2+-NTA Fractions
mAU
A280
Peak 1
S200 Column Chromatogram
Ladder
Ni-Elution pool
Elu pool + Ulp1
S200 Peak 1
Heparin elution
Mtr4
S200
Heparin
Elution
Heparin Column Chromatogram
b
Ladder
Pellet
Supernatant
Flowthrough
Wash 1
Wash 2 (ATP)
Wash 3
Elution 1
Elution 2
Elution pool + Ulp1
His6Smt3-Mtl1
Mtl1
Ni2+-NTA Fractions
Ladder
Load
Peak 1
Peak 2
Peak 3
S200 Column Fractions
S200 Column Chromatogram
Ladder
Load
Flowthrough
Wash
Elution
Heparin Column Fractions
Flowthrough
Wash
Elution
ml
Heparin Column Chromatogram

**c**

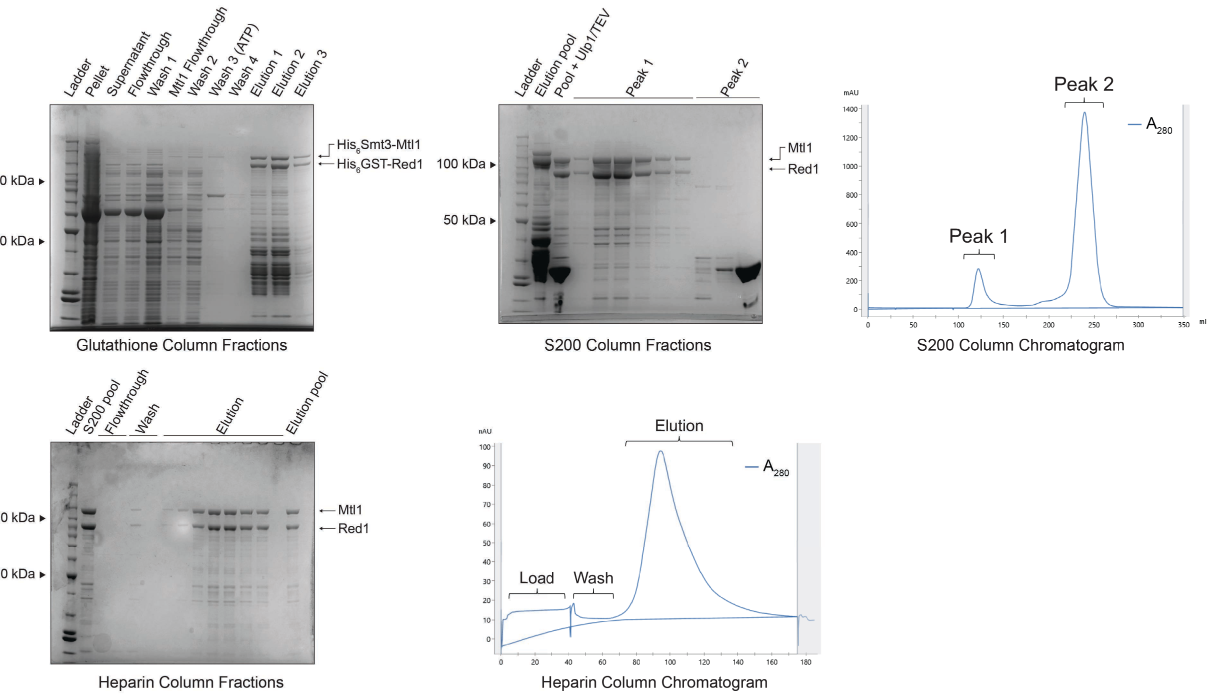
Ladder
Pellet
Supernatant
Flowthrough
Wash 1
Mtl1 Flowthrough
Wash 2
Wash 3 (ATP)
Wash 4
Elution 1
Elution 2
Elution 3
100 kDa
50 kDa
His6Smt3-Mtl1
His6GST-Red1
Glutathione Column Fractions
Ladder
Elution pool
Pool + Ulp1/TEV
Peak 1
Peak 2
100 kDa
50 kDa
Mtl1
Red1
S200 Column Fractions
mAU
Peak 2
Peak 1
A280
ml
S200 Column Chromatogram
Ladder
S200 pool
Flowthrough
Wash
Elution
Elution pool
100 kDa
50 kDa
Mtl1
Red1
Heparin Column Fractions
mAU
Elution
A280
Load
Wash
Heparin Column Chromatogram

**d**

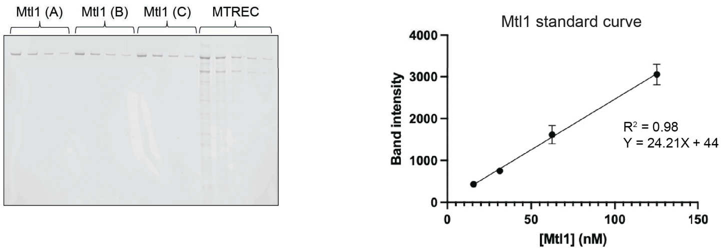
Mtl1 (A)
Mtl1 (B)
Mtl1 (C)
MTREC
Mtl1 standard curve
Band intensity
R² = 0.98
Y = 24.21X + 44
[Mtl1] (nM)

**e**

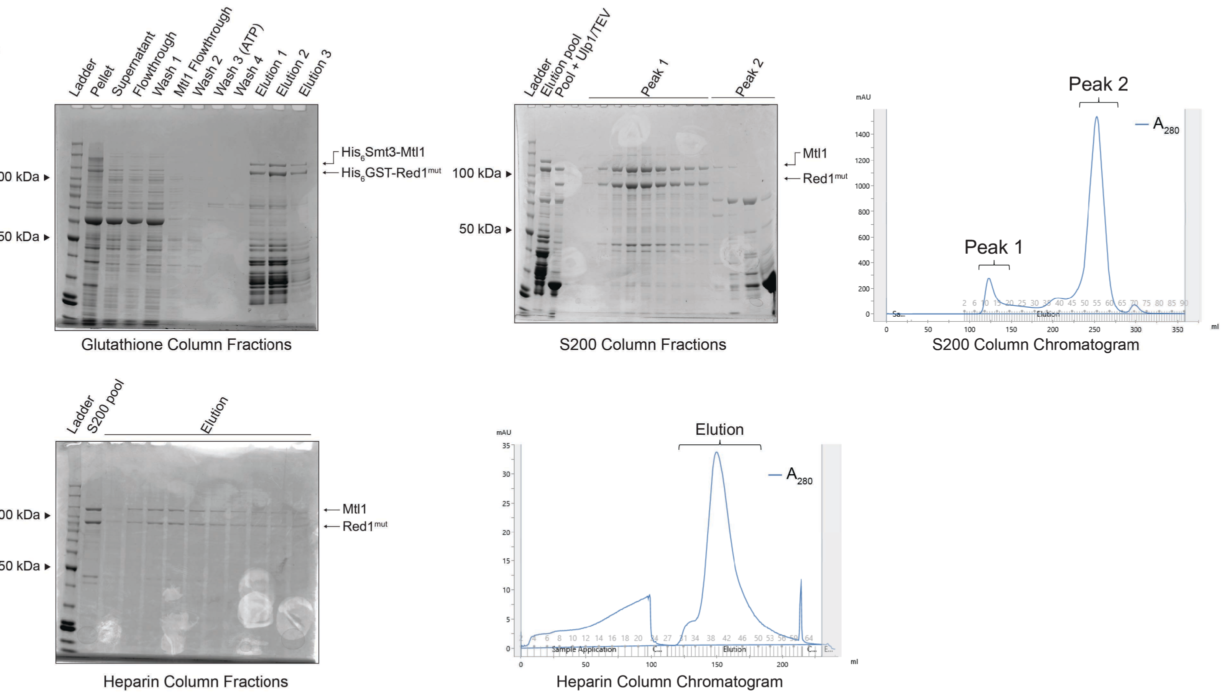


**f**

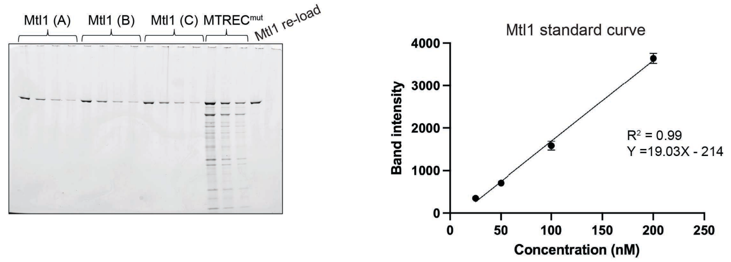


**Figure S1. | Purifications of Mtr4, Mtl1, and MTREC. (a)** SDS-PAGE gels from each step of the purification of Mtr4 from BL21 STAR cells with accompanying chromatograms. **(b)** SDS-PAGE gels from each step of the purification of Mtl1 from SoluBL21 cells with accompanying chromatograms. **(c)** SDS-PAGE gels from each step of the purification of MTREC from Arctic Express and SoluBL21 cells with accompanying chromatograms. **(d)** Sample of gel demonstrating how MTREC concentrations were determined using serial dilutions of Mtl1 to calculate a standard curve based on the fluorescent intensity of SYPRO Ruby (Thermo Fisher) staining. Serial dilutions of MTREC were run on the same gel and band intensities within the linear range of the standard curve were used to calculate the undiluted MTREC concentration. **(e)** SDS-PAGE gels from each step of the purification of $MTREC^{mut}$ from Arctic Express and SoluBL21 cells with accompanying chromatograms. **(f)** Sample of gel demonstrating how $MTREC^{mut}$ concentrations were determined using serial dilutions of Mtl1.

**a**

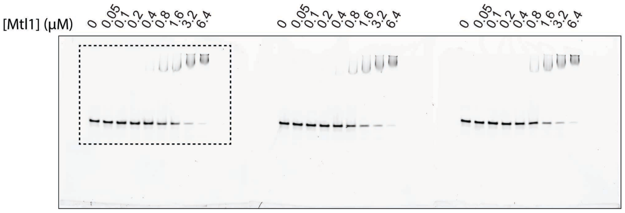


Mtl1 + Substrate 1 (Replicates A-C)

**b**

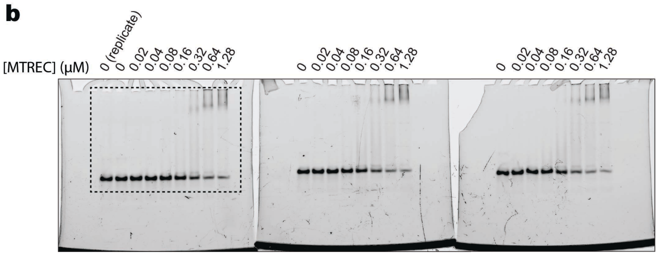


MTREC + Substrate 1 (Replicates A-C)

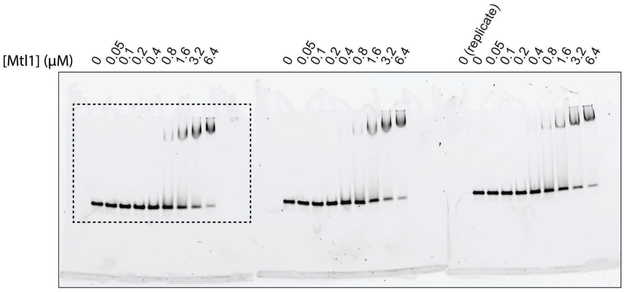


Mtl1 + Substrate 2 (Replicates A-C)

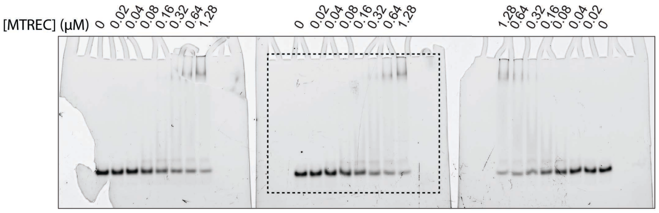


MTREC + Substrate 2 (Replicates A-C)

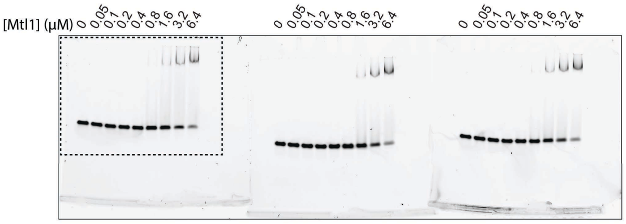


Mtl1 + Substrate 3 (Replicates A-C)

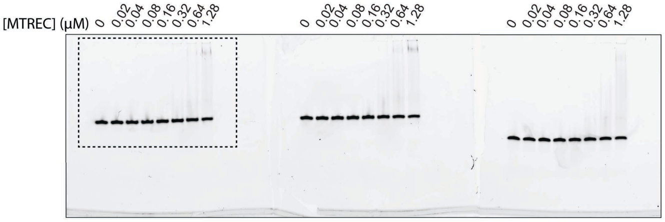


MTREC + Substrate 3 (Replicates A-C)

c

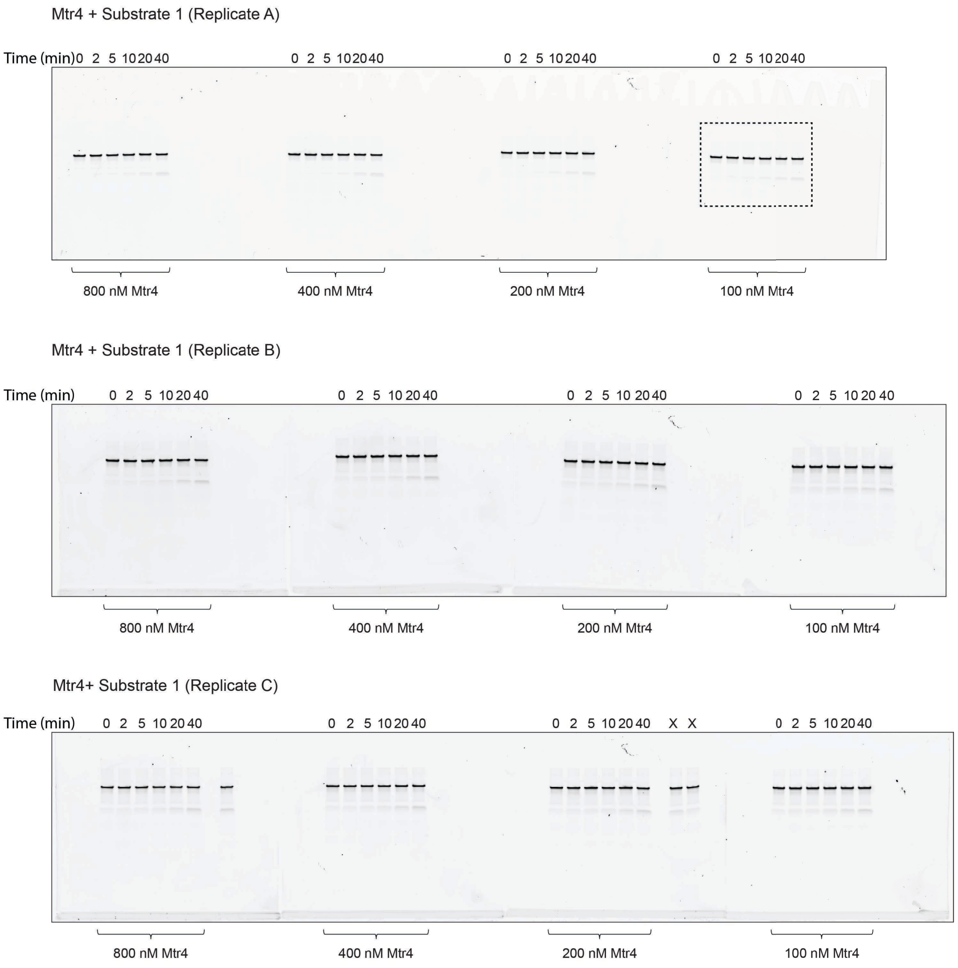
Mtr4 + Substrate 1 (Replicate A)
Time (min) 0 2 5 10 20 40
0 2 5 10 20 40
0 2 5 10 20 40
0 2 5 10 20 40
800 nM Mtr4
400 nM Mtr4
200 nM Mtr4
100 nM Mtr4
Mtr4 + Substrate 1 (Replicate B)
Time (min) 0 2 5 10 20 40
0 2 5 10 20 40
0 2 5 10 20 40
0 2 5 10 20 40
800 nM Mtr4
400 nM Mtr4
200 nM Mtr4
100 nM Mtr4
Mtr4+ Substrate 1 (Replicate C)
Time (min) 0 2 5 10 20 40
0 2 5 10 20 40
0 2 5 10 20 40 X X
0 2 5 10 20 40
800 nM Mtr4
400 nM Mtr4
200 nM Mtr4
100 nM Mtr4

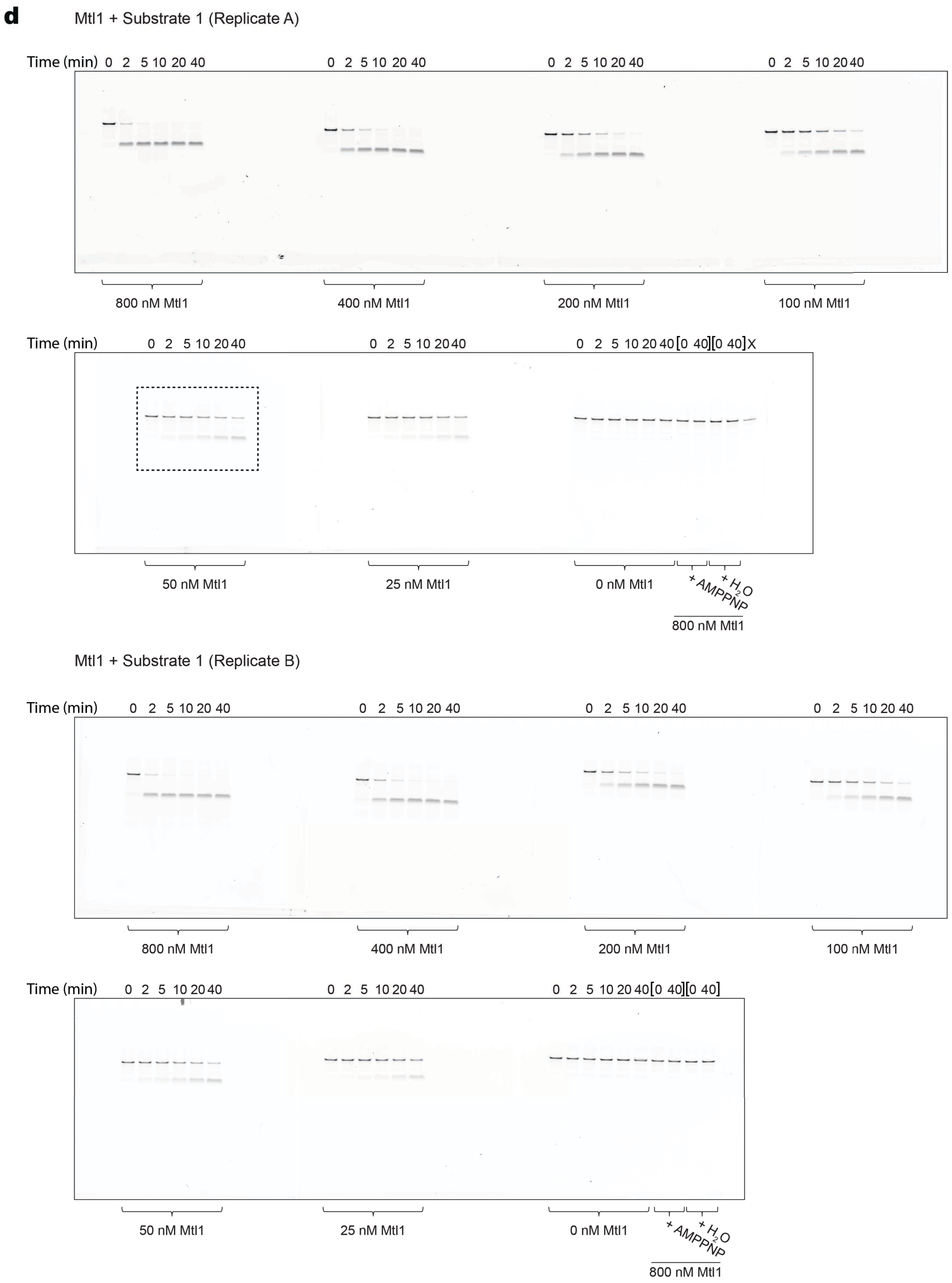
d
Mtl1 + Substrate 1 (Replicate A)
Time (min) 0 2 5 10 20 40 0 2 5 10 20 40 0 2 5 10 20 40 0 2 5 10 20 40
800 nM Mtl1
400 nM Mtl1
200 nM Mtl1
100 nM Mtl1
Time (min) 0 2 5 10 20 40 0 2 5 10 20 40 0 2 5 10 20 40 [0 40][0 40] X
50 nM Mtl1
25 nM Mtl1
0 nM Mtl1
+ AMPPNP
+ H2O
800 nM Mtl1
Mtl1 + Substrate 1 (Replicate B)
Time (min) 0 2 5 10 20 40 0 2 5 10 20 40 0 2 5 10 20 40 0 2 5 10 20 40
800 nM Mtl1
400 nM Mtl1
200 nM Mtl1
100 nM Mtl1
Time (min) 0 2 5 10 20 40 0 2 5 10 20 40 0 2 5 10 20 40 [0 40][0 40]
50 nM Mtl1
25 nM Mtl1
0 nM Mtl1
+ AMPPNP
+ H2O
800 nM Mtl1

Mtl1 + Substrate 1 (Replicate C)

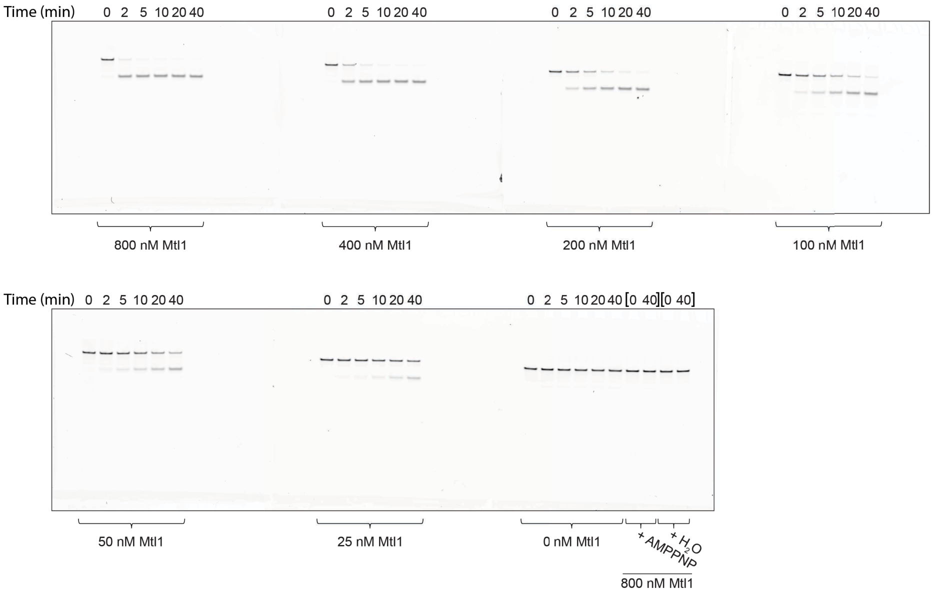


**e**

MTREC + Substrate 1 (Replicate A)

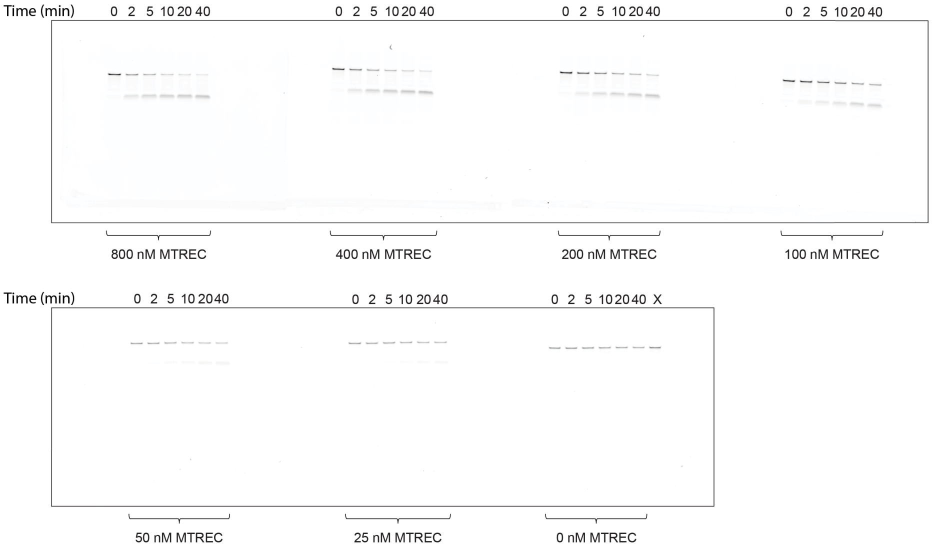

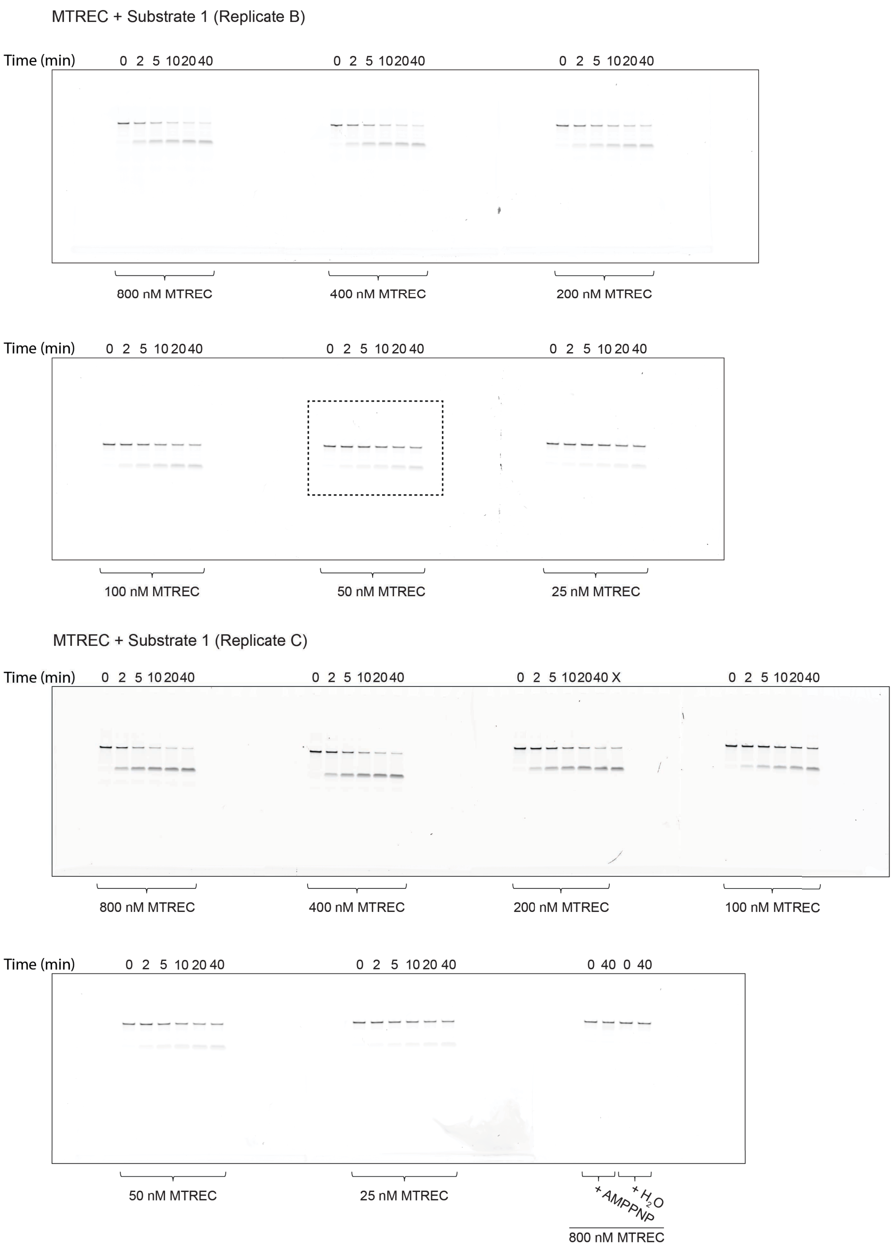


**Figure S2. | EMSA and strand displacement assay data. (a)** Native PAGE gels of EMSA experiments performed with Mtl1 and the three RNA substrates. **(b)** Native PAGE gels of EMSA experiments performed with MTREC and the three RNA substrates. **(c)** Native PAGE gels of strand displacement experiments performed with Mtr4 and the 25 nt poly(A) tailed substrate 1. **(d)** Native PAGE gels of strand displacement experiments performed with Mtl1 and the 25 nt poly(A) tailed substrate 1. **(e)** Native PAGE gels of strand displacement experiments performed with MTREC and the 25 nt poly(A) tailed substrate 1. Lanes labeled with "X" are the result of incorrectly loaded wells, or replicates of other lanes. Dashed box insets in gels indicated cropped gel images in main Figure 1.

**a**

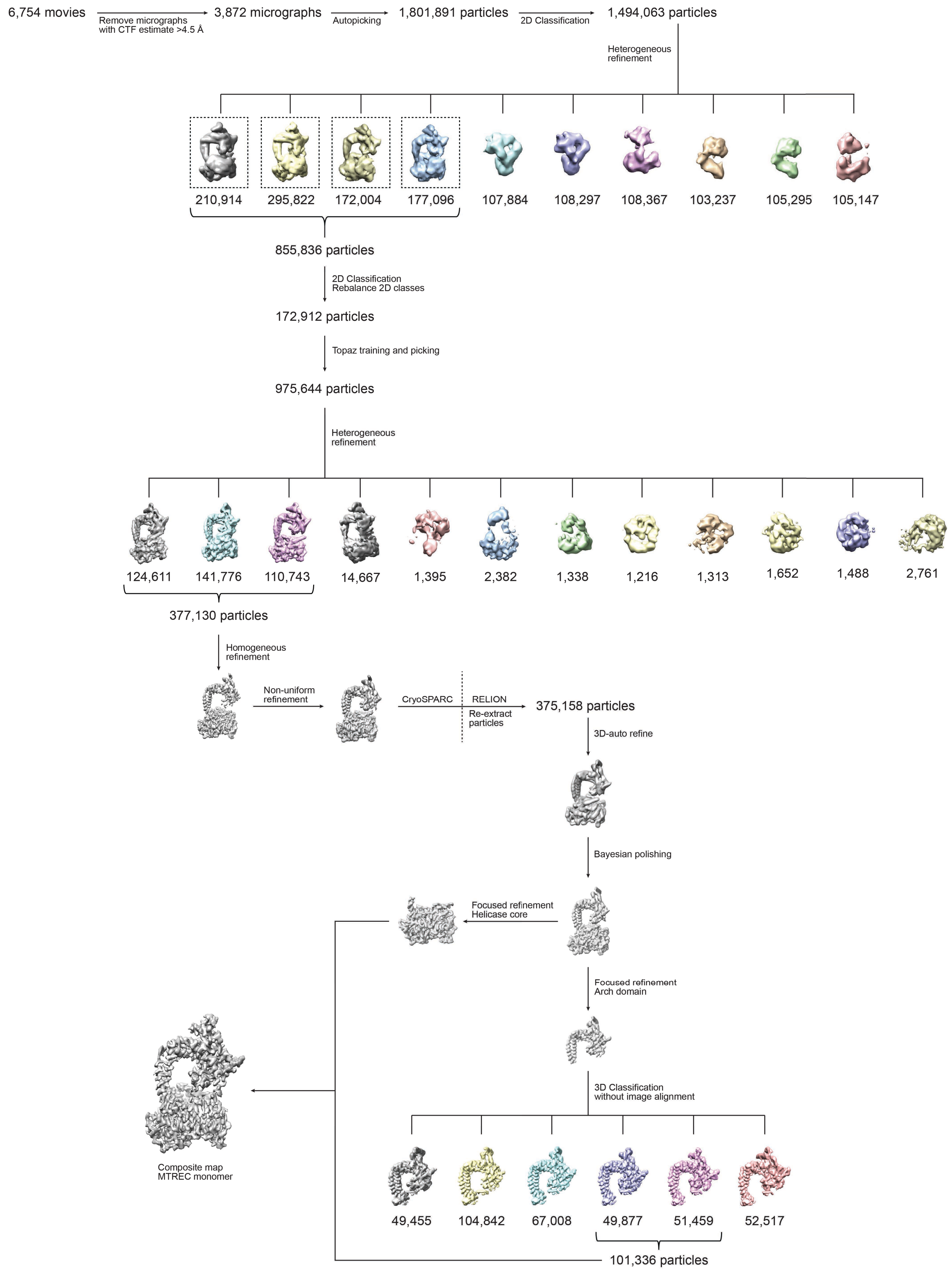

6,754 movies
Remove micrographs with CTF estimate >4.5 Å
3,872 micrographs
Autopicking
1,801,891 particles
2D Classification
1,494,063 particles
Heterogeneous refinement
210,914
295,822
172,004
177,096
107,884
108,297
108,367
103,237
105,295
105,147
855,836 particles
2D Classification Rebalance 2D classes
172,912 particles
Topaz training and picking
975,644 particles
Heterogeneous refinement
124,611
141,776
110,743
14,667
1,395
2,382
1,338
1,216
1,313
1,652
1,488
2,761
377,130 particles
Homogeneous refinement
Non-uniform refinement
CryoSPARC
RELION
Re-extract particles
375,158 particles
3D-auto refine
Bayesian polishing
Focused refinement Helicase core
Focused refinement Arch domain
3D Classification without image alignment
49,455
104,842
67,008
49,877
51,459
52,517
101,336 particles
Composite map MTREC monomer

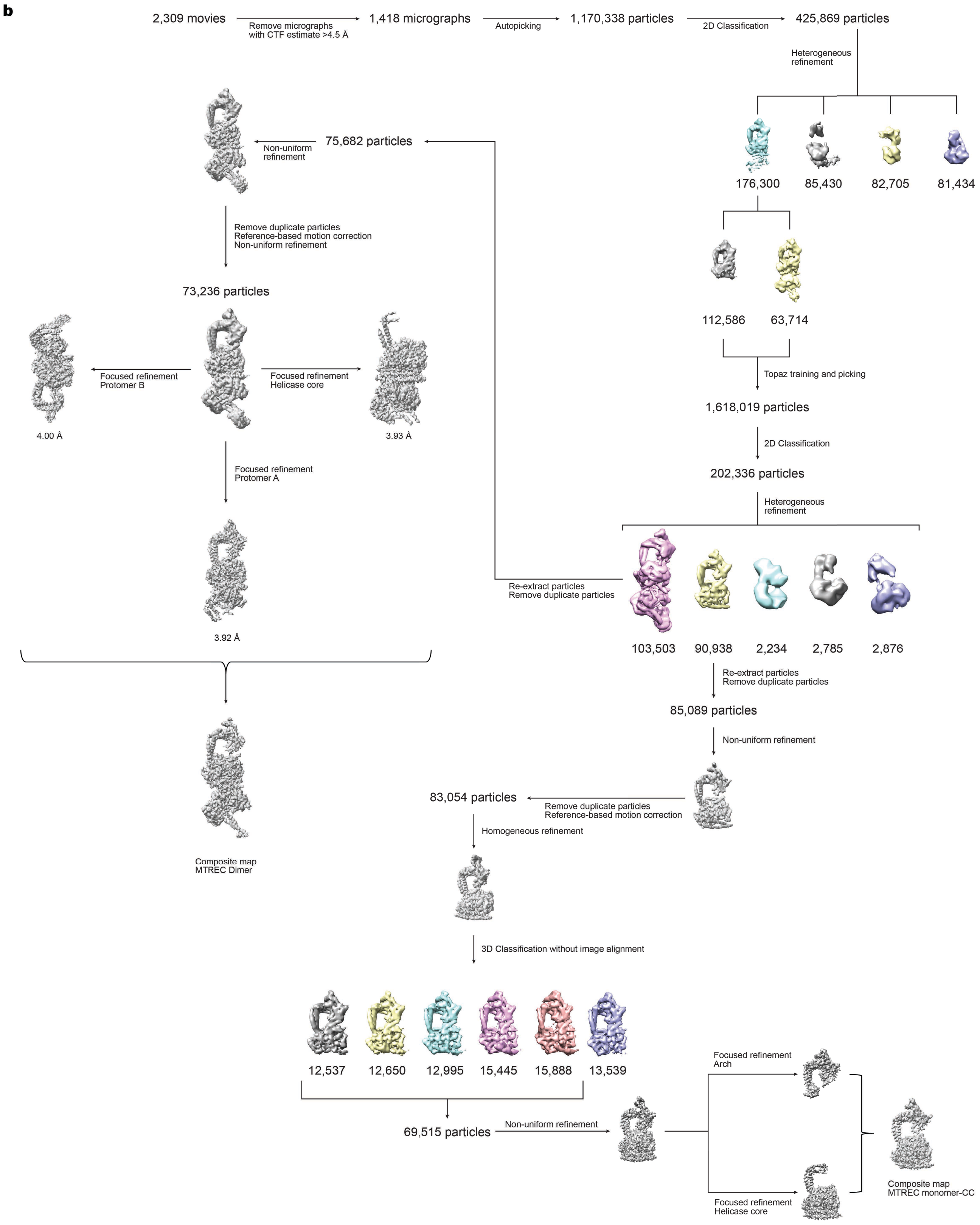


**Figure S3. | Cryo-EM data processing flowcharts. (a)** Data processing flowchart for the MTREC monomer reconstruction. **(b)** Data processing flowchart for the MTREC monomer-CC and MTREC dimer reconstruction.

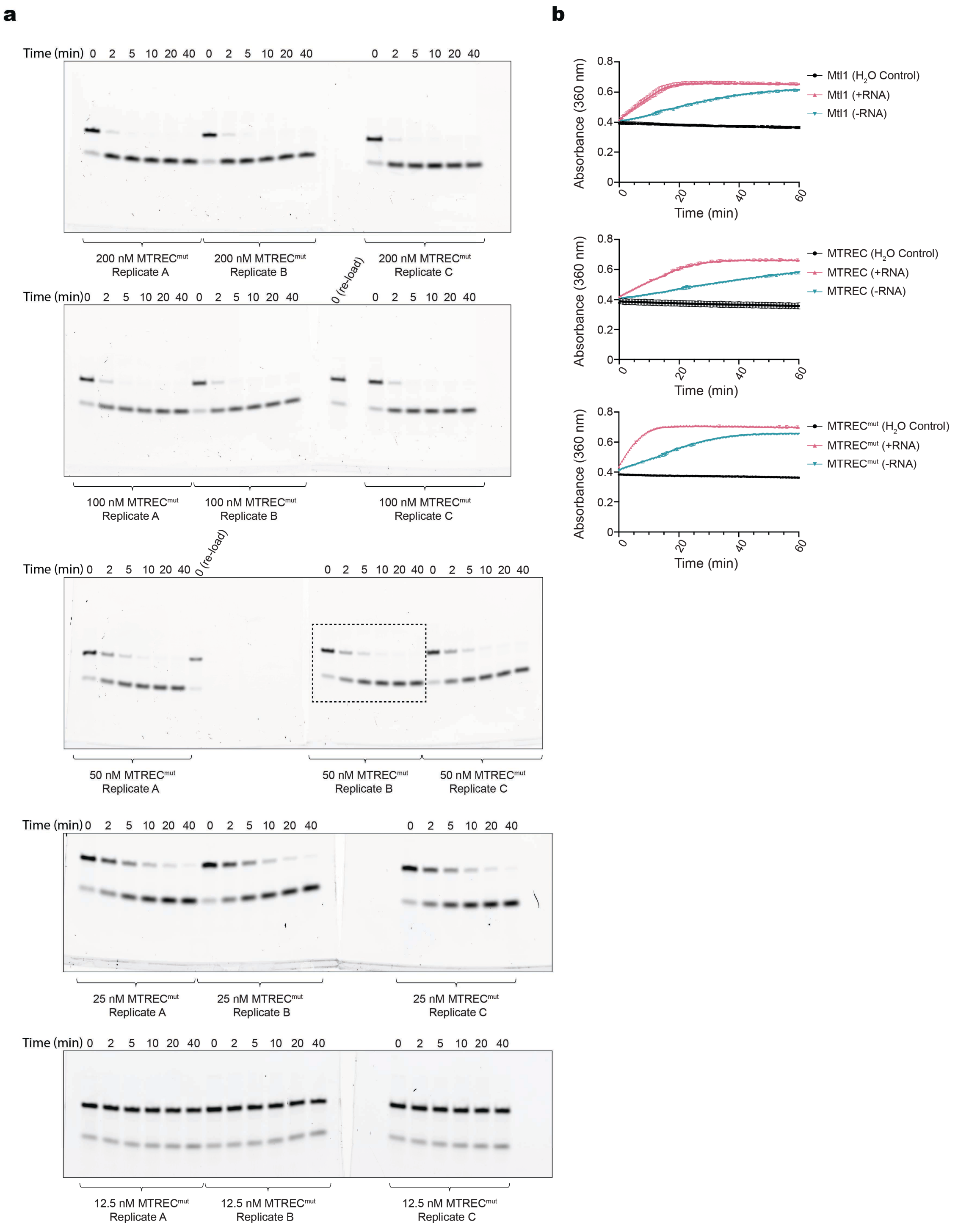


**Figure S4. | MTREC$^{mut}$ strand displacement and ATP hydrolysis data. (a)** Uncropped gels from strand displacement assays performed with MTREC$^{mut}$. **(b)** Raw absorbance data from the ATP hydrolysis assays. Dashed box insets in gels indicated cropped gel images in main Figure 4.

**Mtl1 + Substrate 1 Replicate A**

| [Mtl1] (nM) | Band intensity (I) | $I/I_0$ | $1-I/I_0$ |
|---|---|---|---|
| 0 | 9154.518 | 1 | 0 |
| 50 | 8965.104 | 0.979309233 | 0.020691 |
| 100 | 8919.154 | 0.974289853 | 0.02571 |
| 200 | 9137.64 | 0.99815632 | 0.001844 |
| 400 | 9043.882 | 0.987914601 | 0.012085 |
| 800 | 5692.447 | 0.62181832 | 0.378182 |
| 1600 | 4562.083 | 0.498342239 | 0.501658 |
| 3200 | 1147.163 | 0.125311131 | 0.874689 |
| 6400 | 315.506 | 0.034464512 | 0.965535 |

**Mtl1 + Substrate 1 Replicate B**

| [Mtl1] (nM) | Band intensity (I) | $I/I_0$ | $1-I/I_0$ |
|---|---|---|---|
| 0 | 10762.468 | 1 | 0 |
| 50 | 11318.761 | 1.05168824 | -0.0516882 |
| 100 | 10564.882 | 0.9816412 | 0.0183588 |
| 200 | 10178.882 | 0.94577582 | 0.05422418 |
| 400 | 9110.933 | 0.84654681 | 0.15345319 |
| 800 | 6358.74 | 0.59082545 | 0.40917455 |
| 1600 | 3230.305 | 0.30014538 | 0.69985462 |
| 3200 | 1578.234 | 0.14664239 | 0.85335761 |
| 6400 | 347.213 | 0.03226147 | 0.96773853 |

**Mtl1 + Substrate 1 Replicate C**

| [Mtl1] (nM) | Band intensity (I) | $I/I_0$ | $1-I/I_0$ |
|---|---|---|---|
| 0 | 11676.468 | 1 | 0 |
| 50 | 10554.104 | 0.90387812 | 0.09612188 |
| 100 | 11490.397 | 0.98406444 | 0.01593556 |
| 200 | 10400.861 | 0.89075404 | 0.10924596 |
| 400 | 8846.69 | 0.7576512 | 0.2423488 |
| 800 | 6809.861 | 0.58321241 | 0.41678759 |
| 1600 | 3258.77 | 0.27908868 | 0.72091132 |
| 3200 | 1273.87 | 0.1090972 | 0.8909028 |
| 6400 | 362.213 | 0.03102077 | 0.96897923 |

**Mtl1 + Substrate 2 Replicate A**

| [Mtl1] (nM) | Band intensity (I) | $I/I_0$ | $1-I/I_0$ |
|---|---|---|---|
| 0 | 6263.69 | 1 | 0 |
| 50 | 6202.276 | 0.990195 | 0.009805 |
| 100 | 6645.983 | 1.061033 | -0.061033 |
| 200 | 5958.518 | 0.951279 | 0.048721 |
| 400 | 5539.933 | 0.884452 | 0.115548 |
| 800 | 3766.347 | 0.601298 | 0.398702 |
| 1600 | 2233.276 | 0.356543 | 0.643457 |
| 3200 | 1390.376 | 0.221974 | 0.778026 |
| 6400 | 513.627 | 0.082001 | 0.917999 |

**Mtl1 + Substrate 2 Replicate B**

| [Mtl1] (nM) | Band intensity (I) | $I/I_0$ | $1-I/I_0$ |
|---|---|---|---|
| 0 | 5642.154 | 1 | 0 |
| 50 | 5877.912 | 1.041785 | -0.041785 |
| 100 | 5947.154 | 1.054057 | -0.054057 |
| 200 | 6272.69 | 1.111754 | -0.111754 |
| 400 | 5060.518 | 0.896912 | 0.103088 |
| 800 | 4042.69 | 0.716515 | 0.283485 |
| 1600 | 2174.619 | 0.385424 | 0.614576 |
| 3200 | 1254.891 | 0.222413 | 0.777587 |
| 6400 | 549.456 | 0.097384 | 0.902616 |

**Mtl1 + Substrate 2 Replicate C**

| [Mtl1] (nM) | Band intensity (I) | $I/I_0$ | $1-I/I_0$ |
|---|---|---|---|
| 0 | 7261.497 | 1 | 0 |
| 50 | 6730.497 | 0.926875 | 0.073125 |
| 100 | 6878.447 | 0.947249 | 0.052751 |
| 200 | 6399.447 | 0.881285 | 0.118715 |
| 400 | 5561.811 | 0.765932 | 0.234068 |
| 800 | 3627.69 | 0.499579 | 0.500421 |
| 1600 | 2400.912 | 0.330636 | 0.669364 |
| 3200 | 1131.012 | 0.155755 | 0.844245 |
| 6400 | 557.284 | 0.076745 | 0.923255 |

**Mtl1 + Substrate 3 Replicate A**

| [Mtl1] (nM) | Band intensity (I) | $I/I_0$ | $1-I/I_0$ |
|---|---|---|---|
| 0 | 10106.86 | 1 | 0 |
| 50 | 9055.74 | 0.895999 | 0.104001 |
| 100 | 9389.69 | 0.929041 | 0.070959 |
| 200 | 8825.447 | 0.873213 | 0.126787 |
| 400 | 8624.518 | 0.853333 | 0.146667 |
| 800 | 8828.104 | 0.873476 | 0.126524 |
| 1600 | 6624.983 | 0.655494 | 0.344506 |
| 3200 | 4213.154 | 0.416861 | 0.583139 |
| 6400 | 2251.497 | 0.222769 | 0.777231 |

**Mtl1 + Substrate 3 Replicate B**

| [Mtl1] (nM) | Band intensity (I) | $I/I_0$ | $1-I/I_0$ |
|---|---|---|---|
| 0 | 10649.98 | 1 | 0 |
| 50 | 9903.983 | 0.929953 | 0.070047 |
| 100 | 10115.28 | 0.949793 | 0.050207 |
| 200 | 10224.23 | 0.960023 | 0.039977 |
| 400 | 10171.35 | 0.955058 | 0.044942 |
| 800 | 8645.518 | 0.811787 | 0.188213 |
| 1600 | 7275.589 | 0.683155 | 0.316845 |
| 3200 | 4238.397 | 0.397972 | 0.602028 |
| 6400 | 2172.79 | 0.204018 | 0.795982 |

**Mtl1 + Substrate 3 Replicate C**

| [Mtl1] (nM) | Band intensity (I) | $I/I_0$ | $1-I/I_0$ |
|---|---|---|---|
| 0 | 8857.861 | 1 | 0 |
| 50 | 8693.983 | 0.981499 | 0.018501 |
| 100 | 8646.033 | 0.976086 | 0.023914 |
| 200 | 8247.619 | 0.931107 | 0.068893 |
| 400 | 8534.447 | 0.963488 | 0.036512 |
| 800 | 7744.397 | 0.874297 | 0.125703 |
| 1600 | 5866.64 | 0.662309 | 0.337691 |
| 3200 | 3832.326 | 0.432647 | 0.567353 |
| 6400 | 2047.255 | 0.231123 | 0.768877 |

**MTREC + Substrate 1 Replicate A**

| [MTREC] (nM) | Band intensity (I) | $I/I_0$ | $1-I/I_0$ |
|---|---|---|---|
| 0 | 12786.167 | 1 | 0 |
| 20 | 11328.418 | 0.885990149 | 0.11400985 |
| 40 | 11400.782 | 0.891649702 | 0.1083503 |
| 80 | 10219.024 | 0.799224975 | 0.20077503 |
| 160 | 7556.468 | 0.590987745 | 0.40901226 |
| 320 | 3762.205 | 0.294240252 | 0.70575975 |
| 640 | 2469.841 | 0.193165082 | 0.80683492 |
| 1280 | 1841.77 | 0.14404395 | 0.85595605 |

**MTREC + Substrate 1 Replicate B**

| [MTREC] (nM) | Band intensity (I) | $I/I_0$ | $1-I/I_0$ |
|---|---|---|---|
| 0 | 12725.388 | 1 | 0 |
| 20 | 11857.146 | 0.931770882 | 0.06822912 |
| 40 | 11203.489 | 0.880404511 | 0.11959549 |
| 80 | 11047.853 | 0.868174157 | 0.13182584 |
| 160 | 8339.418 | 0.655337032 | 0.34466297 |
| 320 | 4394.933 | 0.345367308 | 0.65463269 |
| 640 | 2995.033 | 0.235358875 | 0.76464113 |
| 1280 | 2515.962 | 0.197712007 | 0.80228799 |

**MTREC + Substrate 1 Replicate C**

| [MTREC] (nM) | Band intensity (I) | $I/I_0$ | $1-I/I_0$ |
|---|---|---|---|
| 0 | 11454.832 | 1 | 0 |
| 20 | 10355.246 | 0.904006798 | 0.0959932 |
| 40 | 10543.832 | 0.920470243 | 0.07952976 |
| 80 | 9451.418 | 0.825103153 | 0.17489685 |
| 160 | 7122.761 | 0.621812786 | 0.37818721 |
| 320 | 4675.518 | 0.408169932 | 0.59183007 |
| 640 | 2399.426 | 0.209468458 | 0.79053154 |
| 1280 | 1759.912 | 0.153639268 | 0.84636073 |

**MTREC + Substrate 2 Replicate A**

| [MTREC] (nM) | Band intensity (I) | $I/I_0$ | $1-I/I_0$ |
|---|---|---|---|
| 0 | 19495.38 | 1 | 0 |
| 20 | 16146.43 | 0.828218275 | 0.17178172 |
| 40 | 15665.48 | 0.803548328 | 0.19645167 |
| 80 | 12140.066 | 0.622715023 | 0.37728498 |
| 160 | 8560.459 | 0.439101931 | 0.56089807 |
| 320 | 6673.731 | 0.34232372 | 0.65767628 |
| 640 | 5777.903 | 0.296372936 | 0.70362706 |
| 1280 | 3795.882 | 0.194706746 | 0.80529325 |

**MTREC + Substrate 2 Replicate B**

| [MTREC] (nM) | Band intensity (I) | $I/I_0$ | $1-I/I_0$ |
|---|---|---|---|
| 0 | 19031.844 | 1 | 0 |
| 20 | 17677.894 | 0.928858706 | 0.07114129 |
| 40 | 17078.673 | 0.897373528 | 0.10262647 |
| 80 | 14099.622 | 0.740843714 | 0.25915629 |
| 160 | 10904.602 | 0.57296613 | 0.42703387 |
| 320 | 7195.874 | 0.378096521 | 0.62190348 |
| 640 | 5295.024 | 0.278219178 | 0.72178082 |
| 1280 | 4714.489 | 0.247715828 | 0.75228417 |

**MTREC + Substrate 2 Replicate C**

| [MTREC] (nM) | Band intensity (I) | $I/I_0$ | $1-I/I_0$ |
|---|---|---|---|
| 0 | 19856.551 | 1 | 0 |
| 20 | 18005.894 | 0.906798668 | 0.09320133 |
| 40 | 17190.844 | 0.865751761 | 0.13424824 |
| 80 | 13754.894 | 0.69271315 | 0.30728685 |
| 160 | 10495.652 | 0.528573769 | 0.47142623 |
| 320 | 6900.51 | 0.347518056 | 0.65248194 |
| 640 | 5357.368 | 0.269803552 | 0.73019645 |
| 1280 | 4084.882 | 0.205719614 | 0.79428039 |

### MTREC + Substrate 3 Replicate A

| [MTREC] (nM) | Band intensity (I) | $I/I_0$ | $1-I/I_0$ |
|---|---|---|---|
| 0 | 9985.882 | 1 | 0 |
| 20 | 9571.397 | 0.9584929 | 0.0415071 |
| 40 | 8961.518 | 0.897418776 | 0.10258122 |
| 80 | 8861.347 | 0.887387514 | 0.11261249 |
| 160 | 8830.225 | 0.884270914 | 0.11572909 |
| 320 | 8269.376 | 0.828106721 | 0.17189328 |
| 640 | 7563.447 | 0.757414017 | 0.24258598 |
| 1280 | 6226.74 | 0.623554334 | 0.37644567 |

### MTREC + Substrate 3 Replicate B

| [MTREC] (nM) | Band intensity (I) | $I/I_0$ | $1-I/I_0$ |
|---|---|---|---|
| 0 | 9312.761 | 1 | 0 |
| 20 | 9021.397 | 0.968713467 | 0.03128653 |
| 40 | 8929.569 | 0.958853019 | 0.04114698 |
| 80 | 9016.69 | 0.968208032 | 0.03179197 |
| 160 | 8492.083 | 0.911875973 | 0.08812403 |
| 320 | 8025.326 | 0.861755821 | 0.13824418 |
| 640 | 7111.497 | 0.763629282 | 0.23637072 |
| 1280 | 5647.083 | 0.60638118 | 0.39361882 |

### MTREC + Substrate 3 Replicate C

| [MTREC] (nM) | Band intensity (I) | $I/I_0$ | $1-I/I_0$ |
|---|---|---|---|
| 0 | 8714.518 | 1 | 0 |
| 20 | 8423.811 | 0.96664107 | 0.03335893 |
| 40 | 8478.79 | 0.972949967 | 0.02705003 |
| 80 | 8197.397 | 0.940659828 | 0.05934017 |
| 160 | 7635.983 | 0.876236988 | 0.12376301 |
| 320 | 7298.933 | 0.83756015 | 0.16243985 |
| 640 | 6689.276 | 0.767601375 | 0.23239862 |
| 1280 | 5241.861 | 0.601508999 | 0.398491 |

**Table S1.** | **EMSA assay raw data and calculations.**

**800 nM Mtr4 + Substrate 1 Replicate A**

| Time (min) | Band intensity (I) | $I/I_0$ | $1-I/I_0$ |
|---|---|---|---|
| 0 | 3515.941 | 1 | 0 |
| 2 | 3356.991 | 0.95479162 | 0.04520838 |
| 5 | 3079.577 | 0.87588984 | 0.12411016 |
| 10 | 3058.87 | 0.87000038 | 0.12999962 |
| 20 | 3140.991 | 0.89335714 | 0.10664286 |
| 40 | 2897.284 | 0.82404227 | 0.17595773 |

**800 nM Mtr4 + Substrate 1 Replicate B**

| Time (min) | Band intensity (I) | $I/I_0$ | $1-I/I_0$ |
|---|---|---|---|
| 0 | 2953.406 | 1 | 0 |
| 2 | 2651.87 | 0.89790229 | 0.10209771 |
| 5 | 2741.577 | 0.92827637 | 0.07172363 |
| 10 | 2646.749 | 0.89616836 | 0.10383164 |
| 20 | 2568.577 | 0.86969993 | 0.13030007 |
| 40 | 2134.163 | 0.72261078 | 0.27738922 |

**800 nM Mtr4 + Substrate 1 Replicate C**

| Time (min) | Band intensity (I) | $I/I_0$ | $1-I/I_0$ |
|---|---|---|---|
| 0 | 2861.82 | 1 | 0 |
| 2 | 2748.577 | 0.96042973 | 0.03957027 |
| 5 | 2815.284 | 0.98373902 | 0.01626098 |
| 10 | 2665.577 | 0.9314272 | 0.0685728 |
| 20 | 2837.406 | 0.99146907 | 0.00853093 |
| 40 | 2740.82 | 0.95771921 | 0.04228079 |

**400 nM Mtr4 + Substrate 1 Replicate A**

| Time (min) | Band intensity (I) | $I/I_0$ | $1-I/I_0$ |
|---|---|---|---|
| 0 | 3502.941 | 1 | 0 |
| 2 | 3290.406 | 0.9393267 | 0.0606733 |
| 5 | 3515.698 | 1.0036418 | -0.0036418 |
| 10 | 3239.87 | 0.92489996 | 0.07510004 |
| 20 | 3207.991 | 0.91579932 | 0.08420068 |
| 40 | 3012.749 | 0.86006273 | 0.13993727 |

**400 nM Mtr4 + Substrate 1 Replicate B**

| Time (min) | Band intensity (I) | $I/I_0$ | $1-I/I_0$ |
|---|---|---|---|
| 0 | 3082.406 | 1 | 0 |
| 2 | 2991.991 | 0.97066739 | 0.02933261 |
| 5 | 3069.698 | 0.99587725 | 0.00412275 |
| 10 | 2797.991 | 0.90772955 | 0.09227045 |
| 20 | 2674.577 | 0.86769134 | 0.13230866 |
| 40 | 2745.991 | 0.89085961 | 0.10914039 |

### 400 nM Mtr4 + Substrate 1 Replicate C

| Time (min) | Band intensity (I) | $I/I_0$ | $1-I/I_0$ |
|---|---|---|---|
| 0 | 3070.82 | 1 | 0 |
| 2 | 3113.941 | 1.01404218 | -0.0140422 |
| 5 | 3039.406 | 0.98977016 | 0.01022984 |
| 10 | 2763.698 | 0.89998697 | 0.10001303 |
| 20 | 3059.82 | 0.99641789 | 0.00358211 |
| 40 | 2935.234 | 0.95584697 | 0.04415303 |

### 200 nM Mtr4 + Substrate 1 Replicate A

| Time (min) | Band intensity (I) | $I/I_0$ | $1-I/I_0$ |
|---|---|---|---|
| 0 | 3227.991 | 1 | 0 |
| 2 | 3160.284 | 0.97902503 | 0.02097497 |
| 5 | 3252.406 | 1.00756353 | -0.0075635 |
| 10 | 3344.991 | 1.03624545 | -0.0362455 |
| 20 | 3150.284 | 0.97592713 | 0.02407287 |
| 40 | 3063.113 | 0.94892241 | 0.05107759 |

### 200 nM Mtr4 + Substrate 1 Replicate B

| Time (min) | Band intensity (I) | $I/I_0$ | $1-I/I_0$ |
|---|---|---|---|
| 0 | 3031.698 | 1 | 0 |
| 2 | 2849.284 | 0.93983108 | 0.06016892 |
| 5 | 2754.284 | 0.9084955 | 0.0915045 |
| 10 | 2691.456 | 0.8877718 | 0.1122282 |
| 20 | 2741.577 | 0.90430412 | 0.09569588 |
| 40 | 2748.991 | 0.90674962 | 0.09325038 |

### 200 nM Mtr4 + Substrate 1 Replicate C

| Time (min) | Band intensity (I) | $I/I_0$ | $1-I/I_0$ |
|---|---|---|---|
| 0 | 3065.82 | 1 | 0 |
| 2 | 3097.234 | 1.01024652 | -0.0102465 |
| 5 | 3103.062 | 1.01214748 | -0.0121475 |
| 10 | 3042.527 | 0.99240236 | 0.00759764 |
| 20 | 2897.891 | 0.94522542 | 0.05477458 |
| 40 | 2779.77 | 0.90669707 | 0.09330293 |

### 100 nM Mtr4 + Substrate 1 Replicate A

| Time (min) | Band intensity (I) | $I/I_0$ | $1-I/I_0$ |
|---|---|---|---|
| 0 | 3413.698 | 1 | 0 |
| 2 | 3427.991 | 1.00418696 | -0.004187 |
| 5 | 3437.284 | 1.00690922 | -0.0069092 |
| 10 | 3323.284 | 0.97351435 | 0.02648565 |
| 20 | 3332.698 | 0.97627207 | 0.02372793 |
| 40 | 3090.577 | 0.90534576 | 0.09465424 |

**100 nM Mtr4 + Substrate 1 Replicate B**

| Time (min) | Band intensity (I) | $I/I_0$ | $1-I/I_0$ |
|---|---|---|---|
| 0 | 3263.406 | 1 | 0 |
| 2 | 3024.991 | 0.92694289 | 0.07305711 |
| 5 | 3039.87 | 0.93150224 | 0.06849776 |
| 10 | 3100.991 | 0.95023145 | 0.04976855 |
| 20 | 2912.698 | 0.89253314 | 0.10746686 |
| 40 | 2801.406 | 0.85843012 | 0.14156988 |

**100 nM Mtr4 + Substrate 1 Replicate C**

| Time (min) | Band intensity (I) | $I/I_0$ | $1-I/I_0$ |
|---|---|---|---|
| 0 | 3089.698 | 1 | 0 |
| 2 | 3277.527 | 1.06079203 | -0.060792 |
| 5 | 3015.698 | 0.97604944 | 0.02395056 |
| 10 | 3062.698 | 0.99126128 | 0.00873872 |
| 20 | 3135.527 | 1.01483284 | -0.0148328 |
| 40 | 3180.941 | 1.02953137 | -0.0295314 |

**800 nM Mtl1 + Substrate 1 Replicate A**

| Time (min) | Band intensity (I) | $I/I_0$ | $1-I/I_0$ |
|---|---|---|---|
| 0 | 5138.648 | 1 | 0 |
| 2 | 1003.163 | 0.19521925 | 0.80478075 |
| 5 | 278.678 | 0.05423177 | 0.94576823 |
| 10 | 149.435 | 0.02908061 | 0.97091939 |
| 20 | 65.192 | 0.01268661 | 0.98731339 |
| 40 | 62.778 | 0.01221683 | 0.98778317 |

**800 nM Mtl1 + Substrate 1 Replicate B**

| Time (min) | Band intensity (I) | $I/I_0$ | $1-I/I_0$ |
|---|---|---|---|
| 0 | 4328.527 | 1 | 0 |
| 2 | 711.92 | 0.16447166 | 0.83552834 |
| 5 | 184.849 | 0.04270483 | 0.95729517 |
| 10 | 109.314 | 0.02525432 | 0.97474568 |
| 20 | 47.778 | 0.01103794 | 0.98896206 |
| 40 | 46.364 | 0.01071127 | 0.98928873 |

**800 nM Mtl1 + Substrate 1 Replicate C**

| Time (min) | Band intensity (I) | $I/I_0$ | $1-I/I_0$ |
|---|---|---|---|
| 0 | 3670.477 | 1 | 0 |
| 2 | 376.799 | 0.10265668 | 0.89734332 |
| 5 | 69.778 | 0.01901061 | 0.98098939 |
| 10 | 54.95 | 0.01497081 | 0.98502919 |
| 20 | 29.536 | 0.00804691 | 0.99195309 |
| 40 | 24.828 | 0.00676424 | 0.99323576 |

**400 nM Mtl1 + Substrate 1 Replicate A**

| Time (min) | Band intensity (I) | $I/I_0$ | $1-I/I_0$ |
|---|---|---|---|
| 0 | 5839.648 | 1 | 0 |
| 2 | 2330.991 | 0.39916635 | 0.60083365 |
| 5 | 898.749 | 0.15390465 | 0.84609535 |
| 10 | 322.385 | 0.05520624 | 0.94479376 |
| 20 | 188.556 | 0.03228893 | 0.96771107 |
| 40 | 115.728 | 0.01981763 | 0.98018237 |

**400 nM Mtl1 + Substrate 1 Replicate B**

| Time (min) | Band intensity (I) | $I/I_0$ | $1-I/I_0$ |
|---|---|---|---|
| 0 | 4689.941 | 1 | 0 |
| 2 | 1719.456 | 0.36662636 | 0.63337364 |
| 5 | 512.092 | 0.10918943 | 0.89081057 |
| 10 | 216.971 | 0.04626306 | 0.95373694 |
| 20 | 120.728 | 0.0257419 | 0.9742581 |
| 40 | 87.899 | 0.01874203 | 0.98125797 |

**400 nM Mtl1 + Substrate 1 Replicate C**

| Time (min) | Band intensity (I) | $I/I_0$ | $1-I/I_0$ |
|---|---|---|---|
| 0 | 3450.527 | 1 | 0 |
| 2 | 1335.749 | 0.38711449 | 0.61288551 |
| 5 | 373.799 | 0.10833099 | 0.89166901 |
| 10 | 125.435 | 0.03635242 | 0.96364758 |
| 20 | 71.778 | 0.02080204 | 0.97919796 |
| 40 | 65.364 | 0.01894319 | 0.98105681 |

**200 nM Mtl1 + Substrate 1 Replicate A**

| Time (min) | Band intensity (I) | $I/I_0$ | $1-I/I_0$ |
|---|---|---|---|
| 0 | 5643.062 | 1 | 0 |
| 2 | 3774.355 | 0.66884876 | 0.33115124 |
| 5 | 2210.991 | 0.39180697 | 0.60819303 |
| 10 | 1099.749 | 0.19488515 | 0.80511485 |
| 20 | 422.092 | 0.0747984 | 0.9252016 |
| 40 | 188.849 | 0.0334657 | 0.9665343 |

**200 nM Mtl1 + Substrate 1 Replicate B**

| Time (min) | Band intensity (I) | $I/I_0$ | $1-I/I_0$ |
|---|---|---|---|
| 0 | 5143.477 | 1 | 0 |
| 2 | 2954.577 | 0.57443185 | 0.42556815 |
| 5 | 1495.335 | 0.29072454 | 0.70927546 |
| 10 | 810.506 | 0.1575794 | 0.8424206 |
| 20 | 338.678 | 0.06584612 | 0.93415388 |
| 40 | 174.849 | 0.03399432 | 0.96600568 |

**200 nM Mtl1 + Substrate 1 Replicate C**

| Time (min) | Band intensity (I) | $I/I_0$ | $1-I/I_0$ |
|---|---|---|---|
| 0 | 3458.234 | 1 | 0 |
| 2 | 2384.406 | 0.6894866 | 0.3105134 |
| 5 | 1479.87 | 0.42792651 | 0.57207349 |
| 10 | 644.92 | 0.18648825 | 0.81351175 |
| 20 | 304.385 | 0.08801747 | 0.91198253 |
| 40 | 117.021 | 0.03383837 | 0.96616163 |

**100 nM Mtl1 + Substrate 1 Replicate A**

| Time (min) | Band intensity (I) | $I/I_0$ | $1-I/I_0$ |
|---|---|---|---|
| 0 | 6041.648 | 1 | 0 |
| 2 | 4884.234 | 0.80842744 | 0.19157256 |
| 5 | 3993.234 | 0.66095112 | 0.33904888 |
| 10 | 2587.113 | 0.42821313 | 0.57178687 |
| 20 | 1443.87 | 0.23898612 | 0.76101388 |
| 40 | 666.627 | 0.1103386 | 0.8896614 |

**100 nM Mtl1 + Substrate 1 Replicate B**

| Time (min) | Band intensity (I) | $I/I_0$ | $1-I/I_0$ |
|---|---|---|---|
| 0 | 4663.113 | 1 | 0 |
| 2 | 3796.284 | 0.81410937 | 0.18589063 |
| 5 | 2939.991 | 0.63047818 | 0.36952182 |
| 10 | 2080.577 | 0.44617769 | 0.55382231 |
| 20 | 1088.042 | 0.23332954 | 0.76667046 |
| 40 | 480.799 | 0.10310687 | 0.89689313 |

**100 nM Mtl1 + Substrate 1 Replicate C**

| Time (min) | Band intensity (I) | $I/I_0$ | $1-I/I_0$ |
|---|---|---|---|
| 0 | 4529.891 | 1 | 0 |
| 2 | 3262.355 | 0.720184 | 0.279816 |
| 5 | 2552.234 | 0.56342062 | 0.43657938 |
| 10 | 1654.284 | 0.36519289 | 0.63480711 |
| 20 | 912.042 | 0.20133862 | 0.79866138 |
| 40 | 422.506 | 0.09327068 | 0.90672932 |

**50 nM Mtl1 + Substrate 1 Replicate A**

| Time (min) | Band intensity (I) | $I/I_0$ | $1-I/I_0$ |
|---|---|---|---|
| 0 | 4349.113 | 1 | 0 |
| 2 | 3590.698 | 0.82561617 | 0.17438383 |
| 5 | 3566.406 | 0.82003066 | 0.17996934 |
| 10 | 2734.698 | 0.62879442 | 0.37120558 |
| 20 | 2096.577 | 0.48207002 | 0.51792998 |
| 40 | 1351.456 | 0.3107429 | 0.6892571 |

**50 nM Mtl1 + Substrate 1 Replicate B**

| Time (min) | Band intensity (I) | $I/I_0$ | $1-I/I_0$ |
|---|---|---|---|
| 0 | 4149.284 | 1 | 0 |
| 2 | 3958.991 | 0.95413835 | 0.04586165 |
| 5 | 3490.284 | 0.84117742 | 0.15882258 |
| 10 | 2774.284 | 0.66861753 | 0.33138247 |
| 20 | 2076.749 | 0.5005078 | 0.4994922 |
| 40 | 1198.335 | 0.28880525 | 0.71119475 |

**50 nM Mtl1 + Substrate 1 Replicate C**

| Time (min) | Band intensity (I) | $I/I_0$ | $1-I/I_0$ |
|---|---|---|---|
| 0 | 4646.719 | 1 | 0 |
| 2 | 5085.134 | 1.09434937 | -0.0943494 |
| 5 | 4181.891 | 0.89996641 | 0.10003359 |
| 10 | 3729.598 | 0.80263042 | 0.19736958 |
| 20 | 2579.941 | 0.55521778 | 0.44478222 |
| 40 | 1695.406 | 0.36486088 | 0.63513912 |

**25 nM Mtl1 + Substrate 1 Replicate A**

| Time (min) | Band intensity (I) | $I/I_0$ | $1-I/I_0$ |
|---|---|---|---|
| 0 | 4592.113 | 1 | 0 |
| 2 | 4191.113 | 0.91267636 | 0.08732364 |
| 5 | 4083.113 | 0.88915778 | 0.11084222 |
| 10 | 3847.406 | 0.83782912 | 0.16217088 |
| 20 | 3226.113 | 0.70253345 | 0.29746655 |
| 40 | 2435.284 | 0.53031883 | 0.46968117 |

**25 nM Mtl1 + Substrate 1 Replicate B**

| Time (min) | Band intensity (I) | $I/I_0$ | $1-I/I_0$ |
|---|---|---|---|
| 0 | 4218.527 | 1 | 0 |
| 2 | 4163.991 | 0.98707226 | 0.01292774 |
| 5 | 3980.698 | 0.94362274 | 0.05637726 |
| 10 | 3548.991 | 0.84128678 | 0.15871322 |
| 20 | 3166.87 | 0.75070516 | 0.24929484 |
| 40 | 2331.456 | 0.55267064 | 0.44732936 |

**25 nM Mtl1 + Substrate 1 Replicate C**

| Time (min) | Band intensity (I) | $I/I_0$ | $1-I/I_0$ |
|---|---|---|---|
| 0 | 5081.598 | 1 | 0 |
| 2 | 4820.184 | 0.94855673 | 0.05144327 |
| 5 | 4571.477 | 0.89961406 | 0.10038594 |
| 10 | 3983.477 | 0.78390243 | 0.21609757 |
| 20 | 3685.062 | 0.72517779 | 0.27482221 |
| 40 | 3236.062 | 0.63681976 | 0.36318024 |

**800 nM MTREC + Substrate 1 Replicate A**

| Time (min) | Band intensity (I) | $I/I_0$ | $1-I/I_0$ |
|---|---|---|---|
| 0 | 3501.113 | 1 | 0 |
| 2 | 1930.87 | 0.55150177 | 0.44849823 |
| 5 | 1258.456 | 0.35944455 | 0.64055545 |
| 10 | 786.042 | 0.22451203 | 0.77548797 |
| 20 | 601.92 | 0.17192247 | 0.82807753 |
| 40 | 394.971 | 0.11281298 | 0.88718702 |

**800 nM MTREC + Substrate 1 Replicate B**

| Time (min) | Band intensity (I) | $I/I_0$ | $1-I/I_0$ |
|---|---|---|---|
| 0 | 3708.82 | 1 | 0 |
| 2 | 2349.284 | 0.63343166 | 0.36656834 |
| 5 | 1539.456 | 0.41507973 | 0.58492027 |
| 10 | 851.627 | 0.22962209 | 0.77037791 |
| 20 | 569.506 | 0.1535545 | 0.8464455 |
| 40 | 466.799 | 0.12586186 | 0.87413814 |

**800 nM MTREC + Substrate 1 Replicate C**

| Time (min) | Band intensity (I) | $I/I_0$ | $1-I/I_0$ |
|---|---|---|---|
| 0 | 2956.113 | 1 | 0 |
| 2 | 1751.87 | 0.5926262 | 0.4073738 |
| 5 | 966.456 | 0.32693473 | 0.67306527 |
| 10 | 596.92 | 0.20192733 | 0.79807267 |
| 20 | 573.213 | 0.19390768 | 0.80609232 |
| 40 | 330.678 | 0.11186244 | 0.88813756 |

**400 nM MTREC + Substrate 1 Replicate A**

| Time (min) | Band intensity (I) | $I/I_0$ | $1-I/I_0$ |
|---|---|---|---|
| 0 | 3340.113 | 1 | 0 |
| 2 | 2391.991 | 0.71614074 | 0.28385926 |
| 5 | 1730.284 | 0.51803158 | 0.48196842 |
| 10 | 1101.456 | 0.32976609 | 0.67023391 |
| 20 | 745.627 | 0.22323406 | 0.77676594 |
| 40 | 501.385 | 0.15011019 | 0.84988981 |

**400 nM MTREC + Substrate 1 Replicate B**

| Time (min) | Band intensity (I) | $I/I_0$ | $1-I/I_0$ |
|---|---|---|---|
| 0 | 3874.941 | 1 | 0 |
| 2 | 2684.991 | 0.69291145 | 0.30708855 |
| 5 | 1843.456 | 0.47573782 | 0.52426218 |
| 10 | 1197.456 | 0.30902561 | 0.69097439 |
| 20 | 797.456 | 0.20579823 | 0.79420177 |
| 40 | 585.506 | 0.15110062 | 0.84889938 |

**400 nM MTREC + Substrate 1 Replicate C**

| Time (min) | Band intensity (I) | $I/I_0$ | $1-I/I_0$ |
|---|---|---|---|
| 0 | 3289.355 | 1 | 0 |
| 2 | 2167.82 | 0.65904106 | 0.34095894 |
| 5 | 1365.698 | 0.41518717 | 0.58481283 |
| 10 | 788.92 | 0.23984033 | 0.76015967 |
| 20 | 550.506 | 0.16735986 | 0.83264014 |
| 40 | 418.092 | 0.12710455 | 0.87289545 |

**200 nM MTREC + Substrate 1 Replicate A**

| Time (min) | Band intensity (I) | $I/I_0$ | $1-I/I_0$ |
|---|---|---|---|
| 0 | 3563.698 | 1 | 0 |
| 2 | 2917.698 | 0.81872763 | 0.18127237 |
| 5 | 2281.577 | 0.64022737 | 0.35977263 |
| 10 | 1531.456 | 0.42973787 | 0.57026213 |
| 20 | 1123.627 | 0.31529804 | 0.68470196 |
| 40 | 858.335 | 0.24085515 | 0.75914485 |

**200 nM MTREC + Substrate 1 Replicate B**

| Time (min) | Band intensity (I) | $I/I_0$ | $1-I/I_0$ |
|---|---|---|---|
| 0 | 3978.82 | 1 | 0 |
| 2 | 3110.698 | 0.78181421 | 0.21818579 |
| 5 | 2314.577 | 0.58172448 | 0.41827552 |
| 10 | 1668.163 | 0.41926074 | 0.58073926 |
| 20 | 1210.163 | 0.30415123 | 0.69584877 |
| 40 | 1039.163 | 0.26117366 | 0.73882634 |

**200 nM MTREC + Substrate 1 Replicate C**

| Time (min) | Band intensity (I) | $I/I_0$ | $1-I/I_0$ |
|---|---|---|---|
| 0 | 3623.941 | 1 | 0 |
| 2 | 2961.991 | 0.81733974 | 0.18266026 |
| 5 | 2265.406 | 0.62512221 | 0.37487779 |
| 10 | 1378.456 | 0.38037485 | 0.61962515 |
| 20 | 1134.163 | 0.31296398 | 0.68703602 |
| 40 | 614.627 | 0.16960182 | 0.83039818 |

**100 nM MTREC + Substrate 1 Replicate A**

| Time (min) | Band intensity (I) | $I/I_0$ | $1-I/I_0$ |
|---|---|---|---|
| 0 | 3618.82 | 1 | 0 |
| 2 | 3347.991 | 0.92516096 | 0.07483904 |
| 5 | 2847.406 | 0.78683272 | 0.21316728 |
| 10 | 2125.577 | 0.58736743 | 0.41263257 |
| 20 | 1858.577 | 0.51358647 | 0.48641353 |
| 40 | 1350.577 | 0.37320922 | 0.62679078 |

**100 nM MTREC + Substrate 1 Replicate B**

| Time (min) | Band intensity (I) | $I/I_0$ | $1-I/I_0$ |
|---|---|---|---|
| 0 | 3755.648 | 1 | 0 |
| 2 | 3101.82 | 0.82590807 | 0.17409193 |
| 5 | 2744.113 | 0.73066299 | 0.26933701 |
| 10 | 2133.698 | 0.56813045 | 0.43186955 |
| 20 | 1749.284 | 0.46577422 | 0.53422578 |
| 40 | 1529.577 | 0.40727379 | 0.59272621 |

**100 nM MTREC + Substrate 1 Replicate C**

| Time (min) | Band intensity (I) | $I/I_0$ | $1-I/I_0$ |
|---|---|---|---|
| 0 | 3293.941 | 1 | 0 |
| 2 | 2890.941 | 0.87765415 | 0.12234585 |
| 5 | 2554.406 | 0.77548626 | 0.22451374 |
| 10 | 1943.698 | 0.59008282 | 0.40991718 |
| 20 | 1611.577 | 0.48925497 | 0.51074503 |
| 40 | 1415.456 | 0.42971504 | 0.57028496 |

**50 nM MTREC + Substrate 1 Replicate A**

| Time (min) | Band intensity (I) | $I/I_0$ | $1-I/I_0$ |
|---|---|---|---|
| 0 | 2916.991 | 1 | 0 |
| 2 | 2757.577 | 0.94534985 | 0.05465015 |
| 5 | 2628.163 | 0.90098427 | 0.09901573 |
| 10 | 2250.163 | 0.77139868 | 0.22860132 |
| 20 | 1948.163 | 0.66786733 | 0.33213267 |
| 40 | 1687.042 | 0.57835009 | 0.42164991 |

**50 nM MTREC + Substrate 1 Replicate B**

| Time (min) | Band intensity (I) | $I/I_0$ | $1-I/I_0$ |
|---|---|---|---|
| 0 | 3497.82 | 1 | 0 |
| 2 | 3191.113 | 0.91231481 | 0.08768519 |
| 5 | 3187.941 | 0.91140796 | 0.08859204 |
| 10 | 2596.284 | 0.74225775 | 0.25774225 |
| 20 | 2402.698 | 0.68691299 | 0.31308701 |
| 40 | 2231.284 | 0.63790704 | 0.36209296 |

**50 nM MTREC + Substrate 1 Replicate C**

| Time (min) | Band intensity (I) | $I/I_0$ | $1-I/I_0$ |
|---|---|---|---|
| 0 | 3414.355 | 1 | 0 |
| 2 | 3297.527 | 0.96578329 | 0.03421671 |
| 5 | 2996.82 | 0.8777119 | 0.1222881 |
| 10 | 2684.406 | 0.78621174 | 0.21378826 |
| 20 | 2260.406 | 0.66203016 | 0.33796984 |
| 40 | 2241.698 | 0.65655094 | 0.34344906 |

**25 nM MTREC + Substrate 1 Replicate A**

| Time (min) | Band intensity (I) | $I/I_0$ | $1-I/I_0$ |
|---|---|---|---|
| 0 | 3134.991 | 1 | 0 |
| 2 | 2689.284 | 0.8578283 | 0.1421717 |
| 5 | 2973.456 | 0.94847354 | 0.05152646 |
| 10 | 2802.577 | 0.89396652 | 0.10603348 |
| 20 | 2471.991 | 0.78851614 | 0.21148386 |
| 40 | 2385.87 | 0.76104525 | 0.23895475 |

**25 nM MTREC + Substrate 1 Replicate B**

| Time (min) | Band intensity (I) | $I/I_0$ | $1-I/I_0$ |
|---|---|---|---|
| 0 | 3907.234 | 1 | 0 |
| 2 | 3498.941 | 0.89550332 | 0.10449668 |
| 5 | 3332.648 | 0.85294303 | 0.14705697 |
| 10 | 2973.82 | 0.76110619 | 0.23889381 |
| 20 | 2765.113 | 0.70769066 | 0.29230934 |
| 40 | 2631.406 | 0.67347029 | 0.32652971 |

**25 nM MTREC + Substrate 1 Replicate C**

| Time (min) | Band intensity (I) | $I/I_0$ | $1-I/I_0$ |
|---|---|---|---|
| 0 | 3522.355 | 1 | 0 |
| 2 | 3487.648 | 0.99014665 | 0.00985335 |
| 5 | 3117.234 | 0.88498576 | 0.11501424 |
| 10 | 3130.941 | 0.88887719 | 0.11112281 |
| 20 | 2797.113 | 0.79410309 | 0.20589691 |
| 40 | 2725.527 | 0.77377976 | 0.22622024 |

**Table S2. | Strand Displacement assay raw data and calculations for Mtr4, Mtl1, and MTREC.**

| | MTREC reconstructions | | |
|---|---|---|---|
| **Data collection** | Monomer - Dataset | Dimer and Monomer Coil-Coil - Dataset | |
| Microscope | Titan Krios G2 | Titan Krios G2 | |
| Detector/Mode | Gatan K3 Summit | Gatan K3 Summit | |
| Mode | Counting - Super Resolution | Counting - Super Resolution | |
| Data collection software | Serial EM | Serial EM | |
| Energy Filter | n/a | n/a | |
| Magnification | 22,500x | 22,500x | |
| Voltage (kV) | 300 | 300 | |
| Electron exposure ($e^-/Å^2$) | 66 | 66 | |
| Frames (collected/used) | 40/40 | 40/40 | |
| Exposure Time (s) | 3.995 | 3.995 | |
| Defocus range (µm) | -0.8 to -2.3 | -0.8 to -2.3 | |
| Super-resolution pixel size (Å) | 0.532 | 0.532 | |
| Fourier cropped pixel size (Å) | 1.064 | 1.064 | |
| Movies (collected/used) | 6754/3872 | 2309/1418 | |
| Initial particle projections (#) | 1,801,891 | 1,170,338 | |
| **Reconstructions** | Overall Monomer – 1 Mtl1 – 1 Red1 | Overall Monomer-CC – 1 Mtl1 – 2 Red1 CC | Overall Dimer - 2 Mtl1 – 2 Red1 |
| Final particle projections (#) | 375,158 | 69,515 | 73,236 |
| Symmetry | C1 | C1 | C1 |
| Map resolution (Å) FSC threshold = 0.143 | 3.49 | 3.87 | 4.04 |
| Map resolution range (Å) Box (contoured) | 2.73-11.55 (2.93-6.31; 0.013 sig) | 2.87-12.67 (3.31-4.96; 0.13 sig) | 3.18-13.04 (3.45-6.28; 0.06 sig) |
| Map sharpening B factor ($Å^2$) | -94.1 | -143.4 | -127.5 |
| Sphericity (FSC threshold = 0.5) | 0.926 | 0.988 | 0.988 |
| EMDB | EMD-76235 | EMD-76240 | EMD-76244 |
| **Focused Refinement Reconstructions** | RecA domains job221 | RecA-CC domains J245_004 | Protomer A/B RecA-CC domains J267_003 |
| Final particle projections (#) | 375,158 | 69,515 | 73,236 |
| Symmetry | C1 | C1 | C1 |
| Map resolution (Å) FSC threshold = 0.143 | 3.28 | 3.63 | 3.93 |
| Map resolution range (Å) Box (contoured) | 2.59-11.12 (2.88-7.29; 0.018 sig) | 2.61-12.20 (3.09-4.78; 0.13 sig) | 2.81-12.97 (3.42-8.73; 0.06 sig) |
| Map sharpening B factor ($Å^2$) | -96.7 | -135.0 | -132.1 |
| Sphericity (FSC threshold = 0.5) | 0.946 | 0.984 | 0.984 |
| EMDB | EMD-76236 | EMD-76241 | EMD-76245 |
| **Focused Refinement Reconstructions** | Stalk-KOW job214 | Stalk-KOW J246_006 | Protomer A-CC J292_004 |
| Final particle projections (#) | 375,158 | 69,515 | 73,236 |
| Symmetry | C1 | C1 | C1 |
| Map resolution (Å) FSC threshold = 0.143 | 3.54 | 4.14 | 3.92 |
| Map resolution range (Å) Box (contoured) | 2.61-11.87 (2.97-4.59; 0.012 sig) | 3.19-12.96 (3.64-6.14; 0.10 sig) | 2.78-12.73 (3.35-7.04; 0.08 sig) |
| Map sharpening B factor ($Å^2$) | -94.1 | -187.5 | -138.5 |
| Sphericity (FSC threshold = 0.5) | 0.937 | 0.623 | 0.981 |
| EMDB | EMD-76237 | EMD-76242 | EMD-76246 |
| **Focused Refinement Reconstructions** | Just KOW job215 | | Protomer B-CC J293_005 |
| Final particle projections (#) | 375,158 | | 73,236 |
| Symmetry | C1 | | C1 |
| Map resolution (Å) FSC threshold = 0.143 | 3.49 | | 4.01 |
| Map resolution range (Å) Box (contoured) | 2.65-11.61 (3.00-5.93; 0.011 sig) | | 3.04-12.98 (3.42-6.84; 0.06 sig) |
| Map sharpening B factor ($Å^2$) | -104.8 | | -126.0 |
| Sphericity (FSC threshold = 0.5) | 0.931 | | 0.839 |
| EMDB | EMD-76238 | | EMD-76247 |
| **Refinement** | | | |
| Initial models used (PDB code) | AF-O13799-F1-v6 | AF-O13799-F1-v6/AF-Q9UTR8-F1-v6 | AF-O13799-F1-v6/AF-Q9UTR8-F1-v6 |
| Model resolution (Å) FSC threshold = 0.5 | 3.34 | 3.95 | 4.11 |
| EMDB (Composite map) | EMD-76234 | EMD-76239 | EMD-76243 |
| Final model (PDB code) | 11ZX<br>pdb_000011ZX | 11ZY<br>pdb_000011ZY | 11ZZ<br>pdb_000011ZZ |
| **Model composition** | | | |
| Non-hydrogen atoms | 8,640 | 9,133 | 17,087 |
| Protein residues | 1,057 | 1,117 | 2090 |
| Nucleic acid residues (ANPPNP) | 1 | 1 | 2 |
| Ligand ($Zn^{2+}$) | 1 | 1 | 1 |
| **Mean *B* factors** | | | |
| Protein | 100.9 | 213.7 | 284.2 |
| Nucleic acid | 102.1 | 184.7 | 225.1 |
| Ligand ($Zn^{2+}$) | 154.2 | 322.3 | 432.3 |
| **RMS deviations** | | | |
| Bond lengths (Å) | 0.002 | 0.002 | 0.004 |
| Bond angles (°) | 0.440 | 0.481 | 0.527 |
| **Validation** | | | |
| Molprobity score | 1.37 | 1.62 | 1.90 |
| Clashscore | 4.85 | 7.73 | 8.61 |
| CC volume/mask | 0.79/0.79 | 0.60/0.59 | 0.62/0.61 |
| EMRinger score | 3.05 | 1.51 | 1.39 |
| Rotamer Outliers (%) | 0.62 | 0 | 0 |
| C-beta deviations (%) | 0 | 0 | 0 |
| CaBLAM outliers (%) | 1.54 | 1.47 | 2.32 |
| **Ramachandran plot** | | | |
| % favored | 97.43 | 96.82 | 93.34 |
| % allowed | 2.57 | 3.18 | 6.66 |
| % outliers | 0 | 0 | 0 |

**Table S3. | Cryo-EM data and refinement table.**

| Oligo ID | Sequence (5'-3') | Description |
|---|---|---|
| DNA Trap | GCGTCTTTACGGTGCT | DNA trap oligo complementary to LD1001 |
| LDR1000 | 6-FAM-AGCACCGUAAAGACGC | 5' FAM labeled "top" strand |
| LDR1001 | GCGUCUUUACGGUGCUAAAAAAAAAAAAAAAAAAAAAAAAA | 25 nt poly(A) overhang "bottom" strand (substrate 1) |
| LDR1002 | GCGUCUUUACGGUGCUAAAAAAAAAA | 10 nt poly(A) overhang "bottom" strand (substrate 2) |
| RP250 | GCGUCUUUACGGUGCUAAAAA | 5 nt poly(A) overhang "bottom" strand (substrate 3) |
| LDR1004 | GCGUCUUUACGGUGCUUUUUUUUUUU | 10 nt poly(U) overhang strand |

**Table S4. | Oligo table.** Sequence and descriptions of the DNA and RNA oligos used in this study.

**200 nM MTREC$^{mut}$ + Substrate 1 Replicate A**

| Time (min) | Band intensity (I) | $I/I_0$ | $1-I/I_0$ |
|---|---|---|---|
| 0 | 13388.175 | 1 | 0 |
| 2 | 1392.841 | 0.10403517 | 0.89596483 |
| 5 | 424.263 | 0.03168938 | 0.96831062 |
| 10 | 351.263 | 0.02623681 | 0.97376319 |
| 20 | 207.607 | 0.01550674 | 0.98449326 |
| 40 | 110.364 | 0.00824339 | 0.99175661 |

**200 nM MTREC$^{mut}$ + Substrate 1 Replicate B**

| Time (min) | Band intensity (I) | $I/I_0$ | $1-I/I_0$ |
|---|---|---|---|
| 0 | 12470.933 | 1 | 0 |
| 2 | 1369.426 | 0.10980943 | 0.89019057 |
| 5 | 345.678 | 0.0277187 | 0.9722813 |
| 10 | 199.021 | 0.01595879 | 0.98404121 |
| 20 | 273.435 | 0.02192579 | 0.97807421 |
| 40 | 123.192 | 0.00987833 | 0.99012167 |

**200 nM MTREC$^{mut}$ + Substrate 1 Replicate C**

| Time (min) | Band intensity (I) | $I/I_0$ | $1-I/I_0$ |
|---|---|---|---|
| 0 | 13461.832 | 1 | 0 |
| 2 | 1099.305 | 0.08166088 | 0.91833912 |
| 5 | 333.263 | 0.02475614 | 0.97524386 |
| 10 | 277.435 | 0.02060901 | 0.97939099 |
| 20 | 206.021 | 0.01530408 | 0.98469592 |
| 40 | 165.607 | 0.01230197 | 0.98769803 |

**100 nM MTREC$^{mut}$ + Substrate 1 Replicate A**

| Time (min) | Band intensity (I) | $I/I_0$ | $1-I/I_0$ |
|---|---|---|---|
| 0 | 12931.125 | 1 | 0 |
| 2 | 2052.548 | 0.15872927 | 0.84127073 |
| 5 | 502.92 | 0.03889221 | 0.96110779 |
| 10 | 280.435 | 0.02168682 | 0.97831318 |
| 20 | 166.728 | 0.01289354 | 0.98710646 |
| 40 | 157.778 | 0.01220141 | 0.98779859 |

**100 nM MTREC$^{mut}$ + Substrate 1 Replicate B**

| Time (min) | Band intensity (I) | $I/I_0$ | $1-I/I_0$ |
|---|---|---|---|
| 0 | 11842.246 | 1 | 0 |
| 2 | 1967.255 | 0.16612178 | 0.83387822 |
| 5 | 429.335 | 0.03625452 | 0.96374548 |
| 10 | 231.435 | 0.01954317 | 0.98045683 |
| 20 | 205.607 | 0.01736216 | 0.98263784 |
| 40 | 152.778 | 0.0129011 | 0.9870989 |

**100 nM MTREC$^{mut}$ + Substrate 1 Replicate C**

| Time (min) | Band intensity (I) | $I/I_0$ | $1-I/I_0$ |
|---|---|---|---|
| 0 | 12452.004 | 1 | 0 |
| 2 | 2218.376 | 0.17815413 | 0.82184587 |
| 5 | 569.577 | 0.04574179 | 0.95425821 |
| 10 | 248.678 | 0.01997092 | 0.98002908 |
| 20 | 149.607 | 0.01201469 | 0.98798531 |
| 40 | 184.192 | 0.01479216 | 0.98520784 |

**50 nM MTREC$^{mut}$ + Substrate 1 Replicate A**

| Time (min) | Band intensity (I) | $I/I_0$ | $1-I/I_0$ |
|---|---|---|---|
| 0 | 11186.518 | 1 | 0 |
| 2 | 4702.74 | 0.42039355 | 0.57960645 |
| 5 | 1771.134 | 0.15832755 | 0.84167245 |
| 10 | 525.87 | 0.04700927 | 0.95299073 |
| 20 | 243.435 | 0.02176146 | 0.97823854 |
| 40 | 189.021 | 0.01689722 | 0.98310278 |

**50 nM MTREC$^{mut}$ + Substrate 1 Replicate B**

| Time (min) | Band intensity (I) | $I/I_0$ | $1-I/I_0$ |
|---|---|---|---|
| 0 | 11259.69 | 1 | 0 |
| 2 | 3707.205 | 0.32924574 | 0.67075426 |
| 5 | 1674.548 | 0.14872061 | 0.85127939 |
| 10 | 539.284 | 0.0478951 | 0.9521049 |
| 20 | 317.263 | 0.02817689 | 0.97182311 |
| 40 | 156.192 | 0.01387179 | 0.98612821 |

**50 nM MTREC$^{mut}$ + Substrate 1 Replicate C**

| Time (min) | Band intensity (I) | $I/I_0$ | $1-I/I_0$ |
|---|---|---|---|
| 0 | 11271.347 | 1 | 0 |
| 2 | 4490.447 | 0.39839489 | 0.60160511 |
| 5 | 1752.134 | 0.15545028 | 0.84454972 |
| 10 | 573.991 | 0.05092479 | 0.94907521 |
| 20 | 269.849 | 0.02394115 | 0.97605885 |
| 40 | 187.021 | 0.0165926 | 0.9834074 |

**25 nM MTREC$^{mut}$ + Substrate 1 Replicate A**

| Time (min) | Band intensity (I) | $I/I_0$ | $1-I/I_0$ |
|---|---|---|---|
| 0 | 11380.539 | 1 | 0 |
| 2 | 7089.175 | 0.62292085 | 0.37707915 |
| 5 | 4425.104 | 0.38883079 | 0.61116921 |
| 10 | 2063.912 | 0.1813545 | 0.8186455 |
| 20 | 999.477 | 0.08782334 | 0.91217666 |
| 40 | 444.092 | 0.03902205 | 0.96097795 |

### 25 nM MTREC$^{mut}$ + Substrate 1 Replicate B

| Time (min) | Band intensity (I) | $I/I_0$ | $1-I/I_0$ |
|---|---|---|---|
| 0 | 11293.418 | 1 | 0 |
| 2 | 6906.054 | 0.61151141 | 0.38848859 |
| 5 | 3901.983 | 0.34550948 | 0.65449052 |
| 10 | 1675.669 | 0.14837572 | 0.85162428 |
| 20 | 753.527 | 0.06672267 | 0.93327733 |
| 40 | 337.263 | 0.02986368 | 0.97013632 |

### 25 nM MTREC$^{mut}$ + Substrate 1 Replicate C

| Time (min) | Band intensity (I) | $I/I_0$ | $1-I/I_0$ |
|---|---|---|---|
| 0 | 11829.953 | 1 | 0 |
| 2 | 6778.64 | 0.5730065 | 0.4269935 |
| 5 | 3850.276 | 0.32546841 | 0.67453159 |
| 10 | 1796.497 | 0.15186003 | 0.84813997 |
| 20 | 817.527 | 0.06910653 | 0.93089347 |
| 40 | 325.849 | 0.0275444 | 0.9724556 |

### 12.5 nM MTREC$^{mut}$ + Substrate 1 Replicate A

| Time (min) | Band intensity (I) | $I/I_0$ | $1-I/I_0$ |
|---|---|---|---|
| 0 | 9689.64 | 1 | 0 |
| 2 | 7824.447 | 0.80750647 | 0.19249353 |
| 5 | 5909.912 | 0.6099207 | 0.3900793 |
| 10 | 4621.619 | 0.47696499 | 0.52303501 |
| 20 | 3079.083 | 0.31777063 | 0.68222937 |
| 40 | 1738.719 | 0.17944103 | 0.82055897 |

### 12.5 nM MTREC$^{mut}$ + Substrate 1 Replicate B

| Time (min) | Band intensity (I) | $I/I_0$ | $1-I/I_0$ |
|---|---|---|---|
| 0 | 9927.225 | 1 | 0 |
| 2 | 7934.397 | 0.79925629 | 0.20074371 |
| 5 | 6044.569 | 0.60888808 | 0.39111192 |
| 10 | 4377.619 | 0.44097107 | 0.55902893 |
| 20 | 2956.79 | 0.29784658 | 0.70215342 |
| 40 | 1774.719 | 0.17877292 | 0.82122708 |

### 12.5 nM MTREC$^{mut}$ + Substrate 1 Replicate C

| Time (min) | Band intensity (I) | $I/I_0$ | $1-I/I_0$ |
|---|---|---|---|
| 0 | 10744.296 | 1 | 0 |
| 2 | 8164.104 | 0.75985472 | 0.24014528 |
| 5 | 6282.983 | 0.58477382 | 0.41522618 |
| 10 | 4386.74 | 0.40828548 | 0.59171452 |
| 20 | 2815.083 | 0.26200721 | 0.73799279 |
| 40 | 1629.841 | 0.15169361 | 0.84830639 |

**Table S5. | Strand Displacement assay raw data and calculations MTREC$^{mut}$.**

| $[P_i]$ (μM) | Rep A (Abs 360 nm) | Rep B (Abs 360 nm) | Rep C (Abs 360 nm) | Average (Abs 360 nm) |
|---|---|---|---|---|
| 120 | 0.6587 | 0.6571 | 0.6591 | 0.6583 |
| 80 | 0.5792 | 0.5779 | 0.5701 | 0.57573333 |
| 60 | 0.5397 | 0.5306 | 0.5314 | 0.5339 |
| 40 | 0.4861 | 0.4928 | 0.4611 | 0.48 |
| 30 | 0.4652 | 0.4611 | 0.4821 | 0.46946667 |
| 20 | 0.4422 | 0.4372 | 0.4194 | 0.43293333 |
| 15 | 0.4316 | 0.4194 | 0.4318 | 0.4276 |
| 10 | 0.3863 | 0.4172 | 0.3995 | 0.401 |
| 7.5 | 0.4391 | 0.3995 | 0.4123 | 0.41696667 |
| 5 | 0.4061 | 0.4053 | 0.402 | 0.40446667 |
| 3.75 | 0.3991 | 0.402 | 0.4014 | 0.40083333 |
| 2.5 | 0.4018 | 0.3924 | 0.3903 | 0.39483333 |
| 0 | 0.3902 | 0.3903 | 0.3862 | 0.3889 |

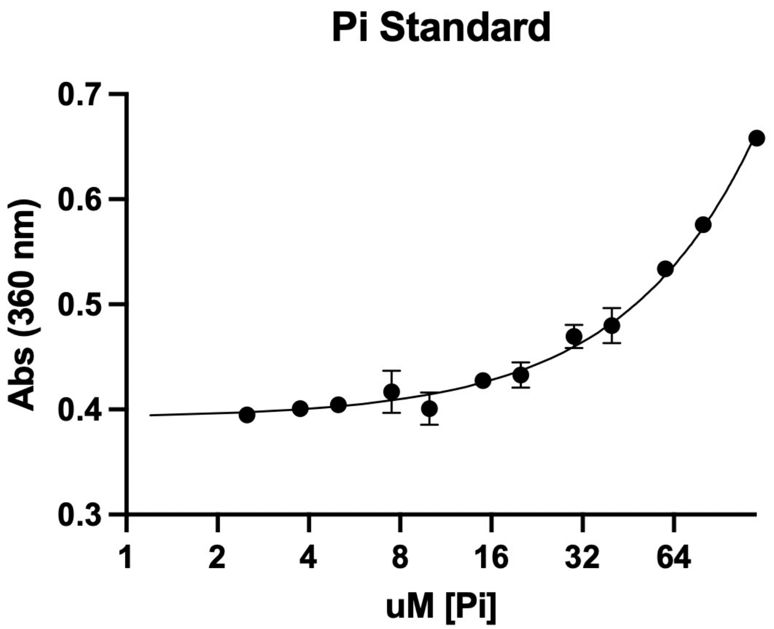


y = 0.0023x + 0.3917 (y is absorbance and x is μM $P_i$)
$R^2 = 0.9944$

**Mtl1 (Abs 360 nm)**

| Time (hr:min:sec) | $H_2O$ Ctrl (Rep A) | $H_2O$ Ctrl (Rep B) | $H_2O$ Ctrl (Rep C) | +RNA (RepA) | +RNA (RepB) | +RNA (RepC) | -RNA (Rep A) | -RNA (Rep B) | -RNA (Rep C) |
|---|---|---|---|---|---|---|---|---|---|
| 0:00:00 | 0.3923 | 0.3932 | 0.3878 | 0.4251 | 0.4147 | 0.4107 | 0.4038 | 0.408 | 0.4069 |
| 0:00:30 | 0.4076 | 0.3938 | 0.3868 | 0.4357 | 0.4242 | 0.4196 | 0.4069 | 0.4074 | 0.4092 |
| 0:01:00 | 0.4088 | 0.3927 | 0.3865 | 0.4455 | 0.4308 | 0.4267 | 0.4097 | 0.4116 | 0.4117 |
| 0:01:30 | 0.4066 | 0.3921 | 0.3865 | 0.4542 | 0.4418 | 0.4367 | 0.4101 | 0.4024 | 0.4135 |
| 0:02:00 | 0.4056 | 0.3914 | 0.3864 | 0.4632 | 0.4501 | 0.4438 | 0.412 | 0.407 | 0.4143 |
| 0:02:30 | 0.4053 | 0.3915 | 0.3875 | 0.4715 | 0.4561 | 0.4513 | 0.4155 | 0.4075 | 0.4155 |
| 0:03:00 | 0.4054 | 0.3897 | 0.3851 | 0.4788 | 0.4618 | 0.4578 | 0.4171 | 0.409 | 0.4184 |
| 0:03:30 | 0.4051 | 0.3903 | 0.3868 | 0.489 | 0.469 | 0.4668 | 0.417 | 0.4116 | 0.4201 |
| 0:04:00 | 0.4051 | 0.3911 | 0.3859 | 0.4964 | 0.4806 | 0.4722 | 0.4194 | 0.414 | 0.4228 |
| 0:04:30 | 0.4043 | 0.3914 | 0.3848 | 0.5041 | 0.4869 | 0.4804 | 0.4224 | 0.4155 | 0.4228 |
| 0:05:00 | 0.4047 | 0.3905 | 0.3842 | 0.5137 | 0.4941 | 0.4872 | 0.4247 | 0.4171 | 0.4242 |
| 0:05:30 | 0.4044 | 0.3884 | 0.3843 | 0.5216 | 0.4998 | 0.4936 | 0.4283 | 0.4187 | 0.4271 |
| 0:06:00 | 0.4036 | 0.3882 | 0.3823 | 0.5295 | 0.5082 | 0.5012 | 0.4273 | 0.4213 | 0.4319 |
| 0:06:30 | 0.403 | 0.3908 | 0.3823 | 0.5367 | 0.5177 | 0.5085 | 0.4309 | 0.42 | 0.4306 |
| 0:07:00 | 0.4034 | 0.3898 | 0.3822 | 0.5467 | 0.5251 | 0.5142 | 0.4344 | 0.4247 | 0.4338 |
| 0:07:30 | 0.4026 | 0.3887 | 0.3816 | 0.5533 | 0.5286 | 0.5212 | 0.4361 | 0.4276 | 0.4369 |
| 0:08:00 | 0.4025 | 0.3852 | 0.3823 | 0.559 | 0.5358 | 0.5284 | 0.4369 | 0.429 | 0.4399 |
| 0:08:30 | 0.4025 | 0.387 | 0.3828 | 0.5687 | 0.5443 | 0.5345 | 0.4401 | 0.4309 | 0.4426 |
| 0:09:00 | 0.4004 | 0.3885 | 0.3805 | 0.5769 | 0.5509 | 0.542 | 0.4425 | 0.4342 | 0.4421 |
| 0:09:30 | 0.4003 | 0.3867 | 0.3797 | 0.5825 | 0.5555 | 0.548 | 0.4451 | 0.4359 | 0.446 |
| 0:10:00 | 0.3996 | 0.3853 | 0.3802 | 0.5901 | 0.5606 | 0.5544 | 0.4475 | 0.4378 | 0.4508 |
| 0:10:30 | 0.3983 | 0.3851 | 0.38 | 0.5969 | 0.5692 | 0.5579 | 0.4496 | 0.4416 | 0.4517 |
| 0:11:00 | 0.3991 | 0.3865 | 0.38 | 0.6044 | 0.5766 | 0.5664 | 0.4535 | 0.443 | 0.4571 |
| 0:11:30 | 0.3958 | 0.3858 | 0.3798 | 0.6141 | 0.5818 | 0.5728 | 0.457 | 0.4458 | 0.4618 |
| 0:12:00 | 0.3981 | 0.3839 | 0.3794 | 0.6184 | 0.5851 | 0.5775 | 0.4595 | 0.4477 | 0.468 |
| 0:12:30 | 0.3957 | 0.3849 | 0.3795 | 0.6243 | 0.5973 | 0.5825 | 0.4638 | 0.4501 | 0.4703 |
| 0:13:00 | 0.3969 | 0.3852 | 0.3785 | 0.6324 | 0.6015 | 0.5923 | 0.4675 | 0.4536 | 0.4746 |
| 0:13:30 | 0.3953 | 0.3856 | 0.3779 | 0.6374 | 0.6109 | 0.5965 | 0.4732 | 0.4588 | 0.476 |
| 0:14:00 | 0.3948 | 0.3825 | 0.3778 | 0.6411 | 0.6132 | 0.6028 | 0.4786 | 0.4655 | 0.4806 |
| 0:14:30 | 0.3938 | 0.3816 | 0.3779 | 0.645 | 0.6202 | 0.6089 | 0.4796 | 0.4705 | 0.4817 |
| 0:15:00 | 0.3925 | 0.3824 | 0.3777 | 0.6484 | 0.6261 | 0.6133 | 0.4772 | 0.4746 | 0.4859 |
| 0:15:30 | 0.3938 | 0.3836 | 0.3759 | 0.6493 | 0.6296 | 0.6197 | 0.4803 | 0.4782 | 0.484 |
| 0:16:00 | 0.3926 | 0.3826 | 0.3763 | 0.6529 | 0.6337 | 0.6245 | 0.4809 | 0.4795 | 0.4872 |
| 0:16:30 | 0.3923 | 0.3813 | 0.3755 | 0.6546 | 0.635 | 0.6291 | 0.4828 | 0.4813 | 0.4899 |
| 0:17:00 | 0.3925 | 0.3795 | 0.3759 | 0.6569 | 0.6373 | 0.6326 | 0.4827 | 0.483 | 0.4947 |
| 0:17:30 | 0.3925 | 0.3811 | 0.3747 | 0.6561 | 0.6437 | 0.633 | 0.4835 | 0.4828 | 0.4949 |
| 0:18:00 | 0.3921 | 0.3818 | 0.3754 | 0.6619 | 0.6468 | 0.6342 | 0.4789 | 0.4842 | 0.4967 |
| 0:18:30 | 0.3911 | 0.3816 | 0.3733 | 0.6631 | 0.6473 | 0.6346 | 0.4876 | 0.4862 | 0.4984 |
| 0:19:00 | 0.3907 | 0.3799 | 0.3736 | 0.6649 | 0.6467 | 0.6385 | 0.4934 | 0.4897 | 0.5016 |
| 0:19:30 | 0.3907 | 0.3798 | 0.3746 | 0.6631 | 0.6478 | 0.6407 | 0.4957 | 0.4915 | 0.508 |
| 0:20:00 | 0.3896 | 0.3786 | 0.3741 | 0.6669 | 0.6526 | 0.6454 | 0.4996 | 0.4935 | 0.5082 |
| 0:20:30 | 0.3899 | 0.3797 | 0.3735 | 0.6671 | 0.6554 | 0.6449 | 0.5032 | 0.4964 | 0.5099 |
| 0:21:00 | 0.3892 | 0.3797 | 0.3732 | 0.6698 | 0.6552 | 0.6478 | 0.5051 | 0.498 | 0.5128 |
| 0:21:30 | 0.3897 | 0.3793 | 0.372 | 0.67 | 0.6577 | 0.6523 | 0.5073 | 0.5013 | 0.5143 |
| 0:22:00 | 0.389 | 0.3779 | 0.3722 | 0.6722 | 0.6539 | 0.6536 | 0.5131 | 0.5034 | 0.5179 |
| 0:22:30 | 0.3892 | 0.3776 | 0.3721 | 0.6702 | 0.6547 | 0.6493 | 0.5158 | 0.5057 | 0.5217 |
| 0:23:00 | 0.389 | 0.3763 | 0.3724 | 0.6699 | 0.6546 | 0.6501 | 0.5192 | 0.5067 | 0.5215 |
| 0:23:30 | 0.3892 | 0.3772 | 0.3709 | 0.6682 | 0.6583 | 0.6517 | 0.5198 | 0.5101 | 0.5242 |
| 0:24:00 | 0.3895 | 0.3781 | 0.3716 | 0.6699 | 0.6617 | 0.6525 | 0.5229 | 0.5098 | 0.524 |
| 0:24:30 | 0.3887 | 0.3783 | 0.3705 | 0.6711 | 0.6563 | 0.6503 | 0.5236 | 0.5139 | 0.525 |
| 0:25:00 | 0.3891 | 0.3786 | 0.3717 | 0.6724 | 0.6598 | 0.6538 | 0.5287 | 0.517 | 0.5296 |
| 0:25:30 | 0.3885 | 0.3766 | 0.3712 | 0.6736 | 0.657 | 0.6546 | 0.5318 | 0.5188 | 0.5313 |
| 0:26:00 | 0.3875 | 0.3752 | 0.3712 | 0.6709 | 0.6555 | 0.655 | 0.5323 | 0.52 | 0.5344 |
| 0:26:30 | 0.3875 | 0.3769 | 0.3693 | 0.6706 | 0.6584 | 0.6565 | 0.5332 | 0.5213 | 0.5355 |
| 0:27:00 | 0.3888 | 0.3762 | 0.3706 | 0.6714 | 0.6559 | 0.6551 | 0.5354 | 0.5262 | 0.5368 |
| 0:27:30 | 0.3887 | 0.3769 | 0.3692 | 0.6739 | 0.6573 | 0.6537 | 0.5387 | 0.5269 | 0.5359 |
| 0:28:00 | 0.3874 | 0.3768 | 0.369 | 0.6724 | 0.6587 | 0.655 | 0.5403 | 0.5272 | 0.5409 |
| 0:28:30 | 0.3873 | 0.375 | 0.3703 | 0.6741 | 0.6585 | 0.6567 | 0.5446 | 0.5313 | 0.545 |

| | | | | | | | | | |
|---|---|---|---|---|---|---|---|---|---|
| 0:29:00 | 0.3857 | 0.3752 | 0.3691 | 0.6743 | 0.6598 | 0.6592 | 0.5453 | 0.5328 | 0.5439 |
| 0:29:30 | 0.3864 | 0.3732 | 0.3685 | 0.673 | 0.6587 | 0.6557 | 0.5478 | 0.5363 | 0.5502 |
| 0:30:00 | 0.3846 | 0.3739 | 0.3691 | 0.6702 | 0.6606 | 0.6543 | 0.55 | 0.537 | 0.5512 |
| 0:30:30 | 0.3836 | 0.3733 | 0.3667 | 0.6741 | 0.6644 | 0.6554 | 0.5484 | 0.5386 | 0.5547 |
| 0:31:00 | 0.3833 | 0.3749 | 0.3669 | 0.669 | 0.6621 | 0.6532 | 0.5506 | 0.5405 | 0.5543 |
| 0:31:30 | 0.3823 | 0.3739 | 0.3674 | 0.6701 | 0.6601 | 0.6527 | 0.5547 | 0.5418 | 0.5547 |
| 0:32:00 | 0.3826 | 0.373 | 0.3663 | 0.6725 | 0.6571 | 0.6529 | 0.5566 | 0.5437 | 0.5571 |
| 0:32:30 | 0.3823 | 0.3716 | 0.3661 | 0.6718 | 0.6575 | 0.6535 | 0.5591 | 0.5475 | 0.5633 |
| 0:33:00 | 0.3826 | 0.3721 | 0.3664 | 0.6735 | 0.6584 | 0.6544 | 0.5582 | 0.5475 | 0.5654 |
| 0:33:30 | 0.3832 | 0.3725 | 0.3666 | 0.6728 | 0.6589 | 0.6536 | 0.5612 | 0.5518 | 0.5639 |
| 0:34:00 | 0.3831 | 0.3744 | 0.3655 | 0.6727 | 0.6613 | 0.654 | 0.5611 | 0.5509 | 0.5631 |
| 0:34:30 | 0.3818 | 0.3746 | 0.3671 | 0.6713 | 0.662 | 0.6538 | 0.5648 | 0.5528 | 0.5644 |
| 0:35:00 | 0.3825 | 0.3738 | 0.3665 | 0.672 | 0.6649 | 0.6531 | 0.567 | 0.5556 | 0.5669 |
| 0:35:30 | 0.3826 | 0.3708 | 0.3674 | 0.6727 | 0.656 | 0.6527 | 0.5705 | 0.5564 | 0.5684 |
| 0:36:00 | 0.3819 | 0.3704 | 0.3656 | 0.673 | 0.6547 | 0.6531 | 0.5709 | 0.5566 | 0.5715 |
| 0:36:30 | 0.3826 | 0.3709 | 0.3645 | 0.6728 | 0.6608 | 0.6516 | 0.5709 | 0.5594 | 0.5745 |
| 0:37:00 | 0.3805 | 0.3711 | 0.3644 | 0.671 | 0.66 | 0.6536 | 0.5717 | 0.5615 | 0.5752 |
| 0:37:30 | 0.3814 | 0.3713 | 0.3652 | 0.6712 | 0.6623 | 0.6537 | 0.5754 | 0.5648 | 0.5753 |
| 0:38:00 | 0.3811 | 0.3711 | 0.3652 | 0.668 | 0.6586 | 0.6533 | 0.5779 | 0.5658 | 0.5746 |
| 0:38:30 | 0.3793 | 0.3707 | 0.3646 | 0.673 | 0.6561 | 0.6548 | 0.5806 | 0.567 | 0.5812 |
| 0:39:00 | 0.3803 | 0.3706 | 0.3638 | 0.6677 | 0.6533 | 0.6545 | 0.5789 | 0.5679 | 0.5824 |
| 0:39:30 | 0.3797 | 0.3689 | 0.3633 | 0.6686 | 0.658 | 0.6541 | 0.5788 | 0.5701 | 0.5826 |
| 0:40:00 | 0.379 | 0.3716 | 0.3637 | 0.6674 | 0.6594 | 0.6532 | 0.5821 | 0.5707 | 0.5842 |
| 0:40:30 | 0.3791 | 0.3719 | 0.363 | 0.6698 | 0.6579 | 0.6545 | 0.5853 | 0.5728 | 0.5826 |
| 0:41:00 | 0.379 | 0.3718 | 0.3631 | 0.6707 | 0.657 | 0.654 | 0.5853 | 0.5736 | 0.5866 |
| 0:41:30 | 0.3786 | 0.3673 | 0.3633 | 0.6695 | 0.6551 | 0.6534 | 0.5883 | 0.576 | 0.5895 |
| 0:42:00 | 0.3789 | 0.368 | 0.3632 | 0.6683 | 0.6567 | 0.6533 | 0.5857 | 0.5757 | 0.5901 |
| 0:42:30 | 0.3787 | 0.3712 | 0.3626 | 0.6678 | 0.6562 | 0.6521 | 0.5894 | 0.5782 | 0.5888 |
| 0:43:00 | 0.378 | 0.3705 | 0.363 | 0.6697 | 0.6568 | 0.6542 | 0.5921 | 0.5798 | 0.5888 |
| 0:43:30 | 0.3782 | 0.3692 | 0.3615 | 0.6677 | 0.658 | 0.6498 | 0.5925 | 0.5792 | 0.5894 |
| 0:44:00 | 0.3773 | 0.3684 | 0.3619 | 0.6686 | 0.6554 | 0.6518 | 0.5941 | 0.5805 | 0.5943 |
| 0:44:30 | 0.3774 | 0.3693 | 0.3616 | 0.6665 | 0.657 | 0.6515 | 0.5926 | 0.5818 | 0.5954 |
| 0:45:00 | 0.3774 | 0.3699 | 0.3603 | 0.6675 | 0.6595 | 0.6513 | 0.5936 | 0.5831 | 0.5952 |
| 0:45:30 | 0.3763 | 0.3686 | 0.3608 | 0.6676 | 0.6559 | 0.6529 | 0.5967 | 0.5861 | 0.5974 |
| 0:46:00 | 0.376 | 0.3669 | 0.3637 | 0.6683 | 0.6543 | 0.6532 | 0.598 | 0.5865 | 0.5979 |
| 0:46:30 | 0.3767 | 0.3665 | 0.3624 | 0.6663 | 0.6555 | 0.6507 | 0.5979 | 0.5863 | 0.6011 |
| 0:47:00 | 0.3755 | 0.3684 | 0.361 | 0.6666 | 0.6581 | 0.6503 | 0.5961 | 0.5872 | 0.597 |
| 0:47:30 | 0.3768 | 0.3691 | 0.3609 | 0.6653 | 0.6589 | 0.6509 | 0.6006 | 0.5884 | 0.604 |
| 0:48:00 | 0.376 | 0.3668 | 0.3599 | 0.6688 | 0.6549 | 0.6518 | 0.6008 | 0.588 | 0.5988 |
| 0:48:30 | 0.3754 | 0.3649 | 0.3597 | 0.6655 | 0.6534 | 0.6513 | 0.6012 | 0.5884 | 0.6014 |
| 0:49:00 | 0.3753 | 0.3668 | 0.3597 | 0.6649 | 0.6559 | 0.6491 | 0.6034 | 0.5915 | 0.6012 |
| 0:49:30 | 0.3754 | 0.3673 | 0.3594 | 0.6648 | 0.6543 | 0.65 | 0.6068 | 0.5936 | 0.6021 |
| 0:50:00 | 0.3743 | 0.3641 | 0.3593 | 0.6654 | 0.6519 | 0.6504 | 0.6045 | 0.5923 | 0.6079 |
| 0:50:30 | 0.3741 | 0.3649 | 0.358 | 0.6669 | 0.654 | 0.6508 | 0.605 | 0.5968 | 0.6089 |
| 0:51:00 | 0.3751 | 0.3685 | 0.359 | 0.6628 | 0.6546 | 0.6503 | 0.6037 | 0.5938 | 0.605 |
| 0:51:30 | 0.374 | 0.3664 | 0.3575 | 0.6633 | 0.6539 | 0.6503 | 0.6067 | 0.5946 | 0.6081 |
| 0:52:00 | 0.3743 | 0.3644 | 0.3569 | 0.6648 | 0.6545 | 0.6512 | 0.61 | 0.597 | 0.6072 |
| 0:52:30 | 0.3739 | 0.3645 | 0.358 | 0.6679 | 0.6548 | 0.6482 | 0.6107 | 0.5975 | 0.6126 |
| 0:53:00 | 0.3739 | 0.3635 | 0.3577 | 0.6647 | 0.652 | 0.6485 | 0.6098 | 0.5962 | 0.6137 |
| 0:53:30 | 0.3741 | 0.3655 | 0.3567 | 0.6668 | 0.6553 | 0.6512 | 0.6074 | 0.5985 | 0.6101 |
| 0:54:00 | 0.3735 | 0.3642 | 0.3569 | 0.6618 | 0.656 | 0.6501 | 0.6083 | 0.6007 | 0.614 |
| 0:54:30 | 0.3732 | 0.3655 | 0.3565 | 0.6637 | 0.655 | 0.6505 | 0.612 | 0.6009 | 0.6117 |
| 0:55:00 | 0.3724 | 0.3632 | 0.3572 | 0.6637 | 0.6531 | 0.6486 | 0.6153 | 0.6007 | 0.6104 |
| 0:55:30 | 0.3726 | 0.3628 | 0.3559 | 0.6638 | 0.651 | 0.6494 | 0.6148 | 0.6032 | 0.6146 |
| 0:56:00 | 0.3726 | 0.3626 | 0.3561 | 0.6635 | 0.6544 | 0.6517 | 0.6125 | 0.6034 | 0.616 |
| 0:56:30 | 0.3722 | 0.3637 | 0.357 | 0.6608 | 0.6537 | 0.6489 | 0.6087 | 0.6047 | 0.6148 |
| 0:57:00 | 0.3723 | 0.3643 | 0.356 | 0.6645 | 0.6517 | 0.6487 | 0.6157 | 0.603 | 0.614 |
| 0:57:30 | 0.3715 | 0.3648 | 0.3564 | 0.6622 | 0.6534 | 0.6501 | 0.6158 | 0.6022 | 0.6148 |
| 0:58:00 | 0.3723 | 0.3633 | 0.3566 | 0.6656 | 0.6527 | 0.6487 | 0.6161 | 0.6029 | 0.6158 |
| 0:58:30 | 0.3716 | 0.3622 | 0.3568 | 0.6633 | 0.652 | 0.6491 | 0.619 | 0.6034 | 0.6166 |

| | | | | | | | | | |
|---|---|---|---|---|---|---|---|---|---|
| 0:59:00 | 0.3718 | 0.3618 | 0.356 | 0.6636 | 0.6525 | 0.6459 | 0.6131 | 0.6025 | 0.6186 |
| 0:59:30 | 0.3713 | 0.3618 | 0.3554 | 0.6617 | 0.6521 | 0.6471 | 0.6149 | 0.6042 | 0.6161 |
| 1:00:00 | 0.371 | 0.3637 | 0.3547 | 0.6599 | 0.6563 | 0.6492 | 0.6178 | 0.6066 | 0.614 |

**Mtl1 (µM $P_i$)**

| Time (hr:min:sec) | H2O Ctrl (Rep A) | H2O Ctrl (Rep B) | H2O Ctrl (Rep C) | +RNA (RepA) | +RNA (RepB) | +RNA (RepC) | -RNA (Rep A) | -RNA (Rep B) | -RNA (Rep C) |
|---|---|---|---|---|---|---|---|---|---|
| 0:00:00 | 0.265 | 0.662 | -1.720 | 14.733 | 10.146 | 8.381 | 5.337 | 7.190 | 6.705 |
| 0:00:30 | 7.014 | 0.926 | -2.161 | 19.409 | 14.336 | 12.307 | 6.705 | 6.925 | 7.719 |
| 0:01:00 | 7.543 | 0.441 | -2.294 | 23.732 | 17.247 | 15.439 | 7.940 | 8.778 | 8.822 |
| 0:01:30 | 6.573 | 0.176 | -2.294 | 27.569 | 22.100 | 19.850 | 8.116 | 4.720 | 9.616 |
| 0:02:00 | 6.131 | -0.132 | -2.338 | 31.539 | 25.761 | 22.982 | 8.955 | 6.749 | 9.969 |
| 0:02:30 | 5.999 | -0.088 | -1.853 | 35.201 | 28.408 | 26.290 | 10.498 | 6.970 | 10.498 |
| 0:03:00 | 6.043 | -0.882 | -2.911 | 38.421 | 30.922 | 29.157 | 11.204 | 7.631 | 11.778 |
| 0:03:30 | 5.911 | -0.618 | -2.161 | 42.920 | 34.098 | 33.127 | 11.160 | 8.778 | 12.528 |
| 0:04:00 | 5.911 | -0.265 | -2.558 | 46.184 | 39.215 | 35.509 | 12.219 | 9.837 | 13.719 |
| 0:04:30 | 5.558 | -0.132 | -3.044 | 49.581 | 41.994 | 39.127 | 13.542 | 10.498 | 13.719 |
| 0:05:00 | 5.734 | -0.529 | -3.308 | 53.816 | 45.170 | 42.126 | 14.557 | 11.204 | 14.336 |
| 0:05:30 | 5.602 | -1.456 | -3.264 | 57.300 | 47.684 | 44.949 | 16.145 | 11.910 | 15.615 |
| 0:06:00 | 5.249 | -1.544 | -4.146 | 60.785 | 51.390 | 48.302 | 15.704 | 13.057 | 17.733 |
| 0:06:30 | 4.985 | -0.397 | -4.146 | 63.961 | 55.580 | 51.522 | 17.292 | 12.483 | 17.159 |
| 0:07:00 | 5.161 | -0.838 | -4.191 | 68.372 | 58.844 | 54.036 | 18.835 | 14.557 | 18.571 |
| 0:07:30 | 4.808 | -1.323 | -4.455 | 71.284 | 60.388 | 57.124 | 19.585 | 15.836 | 19.938 |
| 0:08:00 | 4.764 | -2.867 | -4.146 | 73.798 | 63.564 | 60.300 | 19.938 | 16.453 | 21.262 |
| 0:08:30 | 4.764 | -2.073 | -3.926 | 78.077 | 67.314 | 62.991 | 21.350 | 17.292 | 22.453 |
| 0:09:00 | 3.838 | -1.412 | -4.940 | 81.694 | 70.225 | 66.299 | 22.408 | 18.747 | 22.232 |
| 0:09:30 | 3.794 | -2.206 | -5.293 | 84.164 | 72.254 | 68.946 | 23.555 | 19.497 | 23.952 |
| 0:10:00 | 3.485 | -2.823 | -5.073 | 87.517 | 74.504 | 71.769 | 24.614 | 20.335 | 26.070 |
| 0:10:30 | 2.911 | -2.911 | -5.161 | 90.516 | 78.297 | 73.313 | 25.540 | 22.011 | 26.467 |
| 0:11:00 | 3.264 | -2.294 | -5.161 | 93.824 | 81.562 | 77.062 | 27.261 | 22.629 | 28.849 |
| 0:11:30 | 1.809 | -2.603 | -5.249 | 98.103 | 83.855 | 79.885 | 28.805 | 23.864 | 30.922 |
| 0:12:00 | 2.823 | -3.441 | -5.426 | 100.000 | 85.311 | 81.959 | 29.907 | 24.702 | 33.657 |
| 0:12:30 | 1.764 | -3.000 | -5.382 | 102.603 | 90.693 | 84.164 | 31.804 | 25.761 | 34.671 |
| 0:13:00 | 2.294 | -2.867 | -5.823 | 106.176 | 92.545 | 88.487 | 33.436 | 27.305 | 36.568 |
| 0:13:30 | 1.588 | -2.691 | -6.087 | 108.381 | 96.692 | 90.340 | 35.951 | 29.599 | 37.186 |
| 0:14:00 | 1.367 | -4.058 | -6.131 | 110.013 | 97.706 | 93.119 | 38.333 | 32.554 | 39.215 |
| 0:14:30 | 0.926 | -4.455 | -6.087 | 111.734 | 100.794 | 95.809 | 38.774 | 34.760 | 39.700 |
| 0:15:00 | 0.353 | -4.102 | -6.176 | 113.233 | 103.397 | 97.750 | 37.715 | 36.568 | 41.553 |
| 0:15:30 | 0.926 | -3.573 | -6.970 | 113.630 | 104.940 | 100.573 | 39.082 | 38.156 | 40.715 |
| 0:16:00 | 0.397 | -4.014 | -6.793 | 115.218 | 106.749 | 102.691 | 39.347 | 38.730 | 42.126 |
| 0:16:30 | 0.265 | -4.588 | -7.146 | 115.968 | 107.322 | 104.720 | 40.185 | 39.524 | 43.317 |
| 0:17:00 | 0.353 | -5.382 | -6.970 | 116.983 | 108.337 | 106.264 | 40.141 | 40.273 | 45.434 |
| 0:17:30 | 0.353 | -4.676 | -7.499 | 116.630 | 111.160 | 106.440 | 40.494 | 40.185 | 45.523 |
| 0:18:00 | 0.176 | -4.367 | -7.190 | 119.188 | 112.528 | 106.970 | 38.465 | 40.803 | 46.317 |
| 0:18:30 | -0.265 | -4.455 | -8.116 | 119.718 | 112.748 | 107.146 | 42.303 | 41.685 | 47.067 |
| 0:19:00 | -0.441 | -5.205 | -7.984 | 120.512 | 112.483 | 108.866 | 44.861 | 43.229 | 48.478 |
| 0:19:30 | -0.441 | -5.249 | -7.543 | 119.718 | 112.969 | 109.837 | 45.876 | 44.023 | 51.301 |
| 0:20:00 | -0.926 | -5.779 | -7.764 | 121.394 | 115.086 | 111.910 | 47.596 | 44.905 | 51.390 |
| 0:20:30 | -0.794 | -5.293 | -8.028 | 121.482 | 116.321 | 111.689 | 49.184 | 46.184 | 52.139 |
| 0:21:00 | -1.103 | -5.293 | -8.161 | 122.673 | 116.233 | 112.969 | 50.022 | 46.890 | 53.419 |
| 0:21:30 | -0.882 | -5.470 | -8.690 | 122.761 | 117.336 | 114.954 | 50.993 | 48.346 | 54.080 |
| 0:22:00 | -1.191 | -6.087 | -8.602 | 123.732 | 115.659 | 115.527 | 53.551 | 49.272 | 55.668 |
| 0:22:30 | -1.103 | -6.220 | -8.646 | 122.850 | 116.012 | 113.630 | 54.742 | 50.287 | 57.345 |
| 0:23:00 | -1.191 | -6.793 | -8.513 | 122.717 | 115.968 | 113.983 | 56.242 | 50.728 | 57.256 |
| 0:23:30 | -1.103 | -6.396 | -9.175 | 121.967 | 117.600 | 114.689 | 56.506 | 52.228 | 58.447 |
| 0:24:00 | -0.970 | -5.999 | -8.866 | 122.717 | 119.100 | 115.042 | 57.874 | 52.095 | 58.359 |
| 0:24:30 | -1.323 | -5.911 | -9.352 | 123.247 | 116.718 | 114.071 | 58.183 | 53.904 | 58.800 |
| 0:25:00 | -1.147 | -5.779 | -8.822 | 123.820 | 118.262 | 115.615 | 60.432 | 55.271 | 60.829 |
| 0:25:30 | -1.412 | -6.661 | -9.043 | 124.349 | 117.027 | 115.968 | 61.800 | 56.065 | 61.579 |
| 0:26:00 | -1.853 | -7.278 | -9.043 | 123.158 | 116.365 | 116.145 | 62.020 | 56.595 | 62.947 |
| 0:26:30 | -1.853 | -6.528 | -9.881 | 123.026 | 117.644 | 116.806 | 62.417 | 57.168 | 63.432 |
| 0:27:00 | -1.279 | -6.837 | -9.307 | 123.379 | 116.542 | 116.189 | 63.388 | 59.330 | 64.005 |
| 0:27:30 | -1.323 | -6.528 | -9.925 | 124.482 | 117.159 | 115.571 | 64.843 | 59.638 | 63.608 |
| 0:28:00 | -1.897 | -6.573 | -10.013 | 123.820 | 117.777 | 116.145 | 65.549 | 59.771 | 65.814 |

| | | | | | | | | | |
|---|---|---|---|---|---|---|---|---|---|
| 0:28:30 | -1.941 | -7.367 | -9.440 | 124.570 | 117.689 | 116.895 | 67.446 | 61.579 | 67.622 |
| 0:29:00 | -2.647 | -7.278 | -9.969 | 124.658 | 118.262 | 117.997 | 67.755 | 62.241 | 67.137 |
| 0:29:30 | -2.338 | -8.161 | -10.234 | 124.085 | 117.777 | 116.453 | 68.858 | 63.785 | 69.916 |
| 0:30:00 | -3.132 | -7.852 | -9.969 | 122.850 | 118.615 | 115.836 | 69.828 | 64.094 | 70.357 |
| 0:30:30 | -3.573 | -8.116 | -11.028 | 124.570 | 120.291 | 116.321 | 69.122 | 64.799 | 71.901 |
| 0:31:00 | -3.705 | -7.411 | -10.940 | 122.320 | 119.277 | 115.351 | 70.093 | 65.637 | 71.725 |
| 0:31:30 | -4.146 | -7.852 | -10.719 | 122.805 | 118.394 | 115.130 | 71.901 | 66.211 | 71.901 |
| 0:32:00 | -4.014 | -8.249 | -11.204 | 123.864 | 117.071 | 115.218 | 72.739 | 67.049 | 72.960 |
| 0:32:30 | -4.146 | -8.866 | -11.292 | 123.555 | 117.247 | 115.483 | 73.842 | 68.725 | 75.695 |
| 0:33:00 | -4.014 | -8.646 | -11.160 | 124.305 | 117.644 | 115.880 | 73.445 | 68.725 | 76.621 |
| 0:33:30 | -3.749 | -8.469 | -11.072 | 123.996 | 117.865 | 115.527 | 74.768 | 70.622 | 75.959 |
| 0:34:00 | -3.794 | -7.631 | -11.557 | 123.952 | 118.924 | 115.704 | 74.724 | 70.225 | 75.607 |
| 0:34:30 | -4.367 | -7.543 | -10.851 | 123.335 | 119.232 | 115.615 | 76.356 | 71.063 | 76.180 |
| 0:35:00 | -4.058 | -7.896 | -11.116 | 123.644 | 120.512 | 115.307 | 77.327 | 72.298 | 77.283 |
| 0:35:30 | -4.014 | -9.219 | -10.719 | 123.952 | 116.586 | 115.130 | 78.871 | 72.651 | 77.944 |
| 0:36:00 | -4.323 | -9.396 | -11.513 | 124.085 | 116.012 | 115.307 | 79.047 | 72.739 | 79.312 |
| 0:36:30 | -4.014 | -9.175 | -11.998 | 123.996 | 118.703 | 114.645 | 79.047 | 73.974 | 80.635 |
| 0:37:00 | -4.940 | -9.087 | -12.042 | 123.202 | 118.350 | 115.527 | 79.400 | 74.901 | 80.944 |
| 0:37:30 | -4.543 | -8.999 | -11.689 | 123.291 | 119.365 | 115.571 | 81.032 | 76.356 | 80.988 |
| 0:38:00 | -4.676 | -9.087 | -11.689 | 121.879 | 117.733 | 115.395 | 82.135 | 76.798 | 80.679 |
| 0:38:30 | -5.470 | -9.263 | -11.954 | 124.085 | 116.630 | 116.056 | 83.326 | 77.327 | 83.591 |
| 0:39:00 | -5.029 | -9.307 | -12.307 | 121.747 | 115.395 | 115.924 | 82.576 | 77.724 | 84.120 |
| 0:39:30 | -5.293 | -10.057 | -12.528 | 122.144 | 117.468 | 115.748 | 82.532 | 78.694 | 84.208 |
| 0:40:00 | -5.602 | -8.866 | -12.351 | 121.614 | 118.086 | 115.351 | 83.988 | 78.959 | 84.914 |
| 0:40:30 | -5.558 | -8.734 | -12.660 | 122.673 | 117.424 | 115.924 | 85.399 | 79.885 | 84.208 |
| 0:41:00 | -5.602 | -8.778 | -12.616 | 123.070 | 117.027 | 115.704 | 85.399 | 80.238 | 85.973 |
| 0:41:30 | -5.779 | -10.763 | -12.528 | 122.541 | 116.189 | 115.439 | 86.723 | 81.297 | 87.252 |
| 0:42:00 | -5.646 | -10.454 | -12.572 | 122.011 | 116.895 | 115.395 | 85.576 | 81.165 | 87.517 |
| 0:42:30 | -5.734 | -9.043 | -12.836 | 121.791 | 116.674 | 114.865 | 87.208 | 82.267 | 86.943 |
| 0:43:00 | -6.043 | -9.352 | -12.660 | 122.629 | 116.939 | 115.792 | 88.399 | 82.973 | 86.943 |
| 0:43:30 | -5.955 | -9.925 | -13.322 | 121.747 | 117.468 | 113.851 | 88.575 | 82.708 | 87.208 |
| 0:44:00 | -6.352 | -10.278 | -13.145 | 122.144 | 116.321 | 114.733 | 89.281 | 83.282 | 89.369 |
| 0:44:30 | -6.308 | -9.881 | -13.277 | 121.217 | 117.027 | 114.601 | 88.619 | 83.855 | 89.854 |
| 0:45:00 | -6.308 | -9.616 | -13.851 | 121.659 | 118.130 | 114.513 | 89.060 | 84.429 | 89.766 |
| 0:45:30 | -6.793 | -10.190 | -13.630 | 121.703 | 116.542 | 115.218 | 90.428 | 85.752 | 90.737 |
| 0:46:00 | -6.925 | -10.940 | -12.351 | 122.011 | 115.836 | 115.351 | 91.001 | 85.929 | 90.957 |
| 0:46:30 | -6.617 | -11.116 | -12.925 | 121.129 | 116.365 | 114.248 | 90.957 | 85.840 | 92.369 |
| 0:47:00 | -7.146 | -10.278 | -13.542 | 121.262 | 117.512 | 114.071 | 90.163 | 86.237 | 90.560 |
| 0:47:30 | -6.573 | -9.969 | -13.586 | 120.688 | 117.865 | 114.336 | 92.148 | 86.767 | 93.648 |
| 0:48:00 | -6.925 | -10.984 | -14.027 | 122.232 | 116.101 | 114.733 | 92.236 | 86.590 | 91.354 |
| 0:48:30 | -7.190 | -11.822 | -14.116 | 120.776 | 115.439 | 114.513 | 92.413 | 86.767 | 92.501 |
| 0:49:00 | -7.234 | -10.984 | -14.116 | 120.512 | 116.542 | 113.542 | 93.383 | 88.134 | 92.413 |
| 0:49:30 | -7.190 | -10.763 | -14.248 | 120.468 | 115.836 | 113.939 | 94.883 | 89.060 | 92.810 |
| 0:50:00 | -7.675 | -12.175 | -14.292 | 120.732 | 114.777 | 114.116 | 93.869 | 88.487 | 95.368 |
| 0:50:30 | -7.764 | -11.822 | -14.865 | 121.394 | 115.704 | 114.292 | 94.089 | 90.472 | 95.809 |
| 0:51:00 | -7.322 | -10.234 | -14.424 | 119.585 | 115.968 | 114.071 | 93.516 | 89.149 | 94.089 |
| 0:51:30 | -7.808 | -11.160 | -15.086 | 119.806 | 115.659 | 114.071 | 94.839 | 89.502 | 95.457 |
| 0:52:00 | -7.675 | -12.042 | -15.351 | 120.468 | 115.924 | 114.468 | 96.295 | 90.560 | 95.060 |
| 0:52:30 | -7.852 | -11.998 | -14.865 | 121.835 | 116.056 | 113.145 | 96.603 | 90.781 | 97.442 |
| 0:53:00 | -7.852 | -12.439 | -14.998 | 120.423 | 114.821 | 113.277 | 96.206 | 90.207 | 97.927 |
| 0:53:30 | -7.764 | -11.557 | -15.439 | 121.350 | 116.277 | 114.468 | 95.148 | 91.222 | 96.339 |
| 0:54:00 | -8.028 | -12.131 | -15.351 | 119.144 | 116.586 | 113.983 | 95.545 | 92.192 | 98.059 |
| 0:54:30 | -8.161 | -11.557 | -15.527 | 119.982 | 116.145 | 114.160 | 97.177 | 92.281 | 97.045 |
| 0:55:00 | -8.513 | -12.572 | -15.218 | 119.982 | 115.307 | 113.322 | 98.633 | 92.192 | 96.471 |
| 0:55:30 | -8.425 | -12.748 | -15.792 | 120.026 | 114.380 | 113.674 | 98.412 | 93.295 | 98.324 |
| 0:56:00 | -8.425 | -12.836 | -15.704 | 119.894 | 115.880 | 114.689 | 97.397 | 93.383 | 98.941 |
| 0:56:30 | -8.602 | -12.351 | -15.307 | 118.703 | 115.571 | 113.454 | 95.721 | 93.957 | 98.412 |
| 0:57:00 | -8.558 | -12.086 | -15.748 | 120.335 | 114.689 | 113.366 | 98.809 | 93.207 | 98.059 |
| 0:57:30 | -8.910 | -11.866 | -15.571 | 119.321 | 115.439 | 113.983 | 98.853 | 92.854 | 98.412 |
| 0:58:00 | -8.558 | -12.528 | -15.483 | 120.820 | 115.130 | 113.366 | 98.985 | 93.163 | 98.853 |

| | | | | | | | | | |
|---|---|---|---|---|---|---|---|---|---|
| 0:58:30 | -8.866 | -13.013 | -15.395 | 119.806 | 114.821 | 113.542 | 100.265 | 93.383 | 99.206 |
| 0:59:00 | -8.778 | -13.189 | -15.748 | 119.938 | 115.042 | 112.131 | 97.662 | 92.986 | 100.088 |
| 0:59:30 | -8.999 | -13.189 | -16.012 | 119.100 | 114.865 | 112.660 | 98.456 | 93.736 | 98.985 |
| 1:00:00 | -9.131 | -12.351 | -16.321 | 118.306 | 116.718 | 113.586 | 99.735 | 94.795 | 98.059 |

**MTREC (Abs 360 nm)**

| Time (hr:min:sec) | $H_2O$ Ctrl (Rep A) | $H_2O$ Ctrl (Rep B) | $H_2O$ Ctrl (Rep C) | +RNA (RepA) | +RNA (RepB) | +RNA (RepC) | -RNA (Rep A) | -RNA (Rep B) | -RNA (Rep C) |
|---|---|---|---|---|---|---|---|---|---|
| 0:00:00 | 0.3792 | 0.3757 | 0.3797 | 0.4204 | 0.421 | 0.4174 | 0.4081 | 0.4027 | 0.4024 |
| 0:00:30 | 0.4094 | 0.3761 | 0.3812 | 0.4275 | 0.428 | 0.4255 | 0.41 | 0.402 | 0.4044 |
| 0:01:00 | 0.4076 | 0.3759 | 0.3804 | 0.4319 | 0.4327 | 0.4307 | 0.413 | 0.4053 | 0.4067 |
| 0:01:30 | 0.4077 | 0.374 | 0.378 | 0.4382 | 0.4385 | 0.4332 | 0.4155 | 0.4071 | 0.4078 |
| 0:02:00 | 0.41 | 0.3713 | 0.3767 | 0.446 | 0.4416 | 0.4389 | 0.415 | 0.4062 | 0.4098 |
| 0:02:30 | 0.408 | 0.3733 | 0.3789 | 0.4518 | 0.4488 | 0.4441 | 0.4203 | 0.4076 | 0.411 |
| 0:03:00 | 0.4057 | 0.3729 | 0.378 | 0.4542 | 0.4547 | 0.4507 | 0.4195 | 0.4103 | 0.4129 |
| 0:03:30 | 0.4071 | 0.3732 | 0.376 | 0.4577 | 0.4603 | 0.4519 | 0.4189 | 0.4142 | 0.4111 |
| 0:04:00 | 0.4085 | 0.3727 | 0.3748 | 0.4672 | 0.4641 | 0.4556 | 0.4187 | 0.414 | 0.4152 |
| 0:04:30 | 0.4073 | 0.3704 | 0.3764 | 0.472 | 0.4679 | 0.4637 | 0.419 | 0.4135 | 0.4155 |
| 0:05:00 | 0.4052 | 0.3716 | 0.3779 | 0.4758 | 0.4758 | 0.4681 | 0.4191 | 0.414 | 0.4173 |
| 0:05:30 | 0.4051 | 0.3711 | 0.3759 | 0.4799 | 0.48 | 0.4736 | 0.4217 | 0.4188 | 0.4191 |
| 0:06:00 | 0.4053 | 0.3703 | 0.3741 | 0.4854 | 0.4854 | 0.4773 | 0.4235 | 0.421 | 0.4211 |
| 0:06:30 | 0.4066 | 0.3697 | 0.3752 | 0.493 | 0.4901 | 0.4809 | 0.4249 | 0.4204 | 0.4223 |
| 0:07:00 | 0.4048 | 0.3686 | 0.3753 | 0.4953 | 0.4947 | 0.4872 | 0.4261 | 0.4204 | 0.4225 |
| 0:07:30 | 0.4047 | 0.3687 | 0.3759 | 0.4985 | 0.5014 | 0.4908 | 0.4285 | 0.4249 | 0.4243 |
| 0:08:00 | 0.4056 | 0.3692 | 0.3731 | 0.5057 | 0.5048 | 0.4957 | 0.4305 | 0.4282 | 0.427 |
| 0:08:30 | 0.4075 | 0.3686 | 0.3729 | 0.5115 | 0.5078 | 0.4977 | 0.4313 | 0.4276 | 0.4279 |
| 0:09:00 | 0.4047 | 0.3691 | 0.3739 | 0.5171 | 0.5154 | 0.5055 | 0.4331 | 0.4273 | 0.4295 |
| 0:09:30 | 0.401 | 0.3686 | 0.3746 | 0.52 | 0.5185 | 0.5103 | 0.4345 | 0.4293 | 0.4306 |
| 0:10:00 | 0.4026 | 0.3683 | 0.3727 | 0.5241 | 0.5255 | 0.5125 | 0.4376 | 0.4326 | 0.4328 |
| 0:10:30 | 0.4033 | 0.3675 | 0.3714 | 0.5317 | 0.5292 | 0.5167 | 0.4394 | 0.4326 | 0.4346 |
| 0:11:00 | 0.4042 | 0.3648 | 0.373 | 0.5353 | 0.5327 | 0.5225 | 0.4403 | 0.4344 | 0.4358 |
| 0:11:30 | 0.4006 | 0.3654 | 0.374 | 0.5377 | 0.54 | 0.5266 | 0.4414 | 0.4375 | 0.4371 |
| 0:12:00 | 0.4008 | 0.3669 | 0.3703 | 0.5422 | 0.5428 | 0.5289 | 0.4444 | 0.4382 | 0.4384 |
| 0:12:30 | 0.4008 | 0.3665 | 0.3711 | 0.5457 | 0.5453 | 0.5322 | 0.4468 | 0.441 | 0.4406 |
| 0:13:00 | 0.4023 | 0.3649 | 0.3702 | 0.5526 | 0.5493 | 0.538 | 0.4481 | 0.4413 | 0.4415 |
| 0:13:30 | 0.4022 | 0.3639 | 0.3716 | 0.5576 | 0.5525 | 0.5433 | 0.4488 | 0.4399 | 0.442 |
| 0:14:00 | 0.3985 | 0.3651 | 0.3718 | 0.5593 | 0.5573 | 0.5467 | 0.4505 | 0.4429 | 0.4439 |
| 0:14:30 | 0.3986 | 0.3635 | 0.3708 | 0.5606 | 0.5628 | 0.5493 | 0.4525 | 0.4451 | 0.4431 |
| 0:15:00 | 0.3997 | 0.3634 | 0.3702 | 0.5677 | 0.5658 | 0.5522 | 0.4535 | 0.4462 | 0.4451 |
| 0:15:30 | 0.4009 | 0.3624 | 0.3701 | 0.5728 | 0.5704 | 0.5575 | 0.4546 | 0.4456 | 0.4498 |
| 0:16:00 | 0.3992 | 0.3636 | 0.3716 | 0.5722 | 0.5793 | 0.567 | 0.4566 | 0.4486 | 0.4509 |
| 0:16:30 | 0.3975 | 0.363 | 0.3702 | 0.5796 | 0.5833 | 0.5714 | 0.4587 | 0.4544 | 0.4538 |
| 0:17:00 | 0.3973 | 0.3629 | 0.3686 | 0.586 | 0.5883 | 0.5739 | 0.4607 | 0.4575 | 0.4549 |
| 0:17:30 | 0.4 | 0.3621 | 0.3685 | 0.5953 | 0.5932 | 0.5772 | 0.4624 | 0.4575 | 0.4563 |
| 0:18:00 | 0.3991 | 0.3619 | 0.3696 | 0.6023 | 0.5965 | 0.5848 | 0.4662 | 0.4608 | 0.4588 |
| 0:18:30 | 0.3981 | 0.3601 | 0.3708 | 0.6026 | 0.6007 | 0.5896 | 0.4675 | 0.4607 | 0.4617 |
| 0:19:00 | 0.3965 | 0.3618 | 0.37 | 0.6062 | 0.6024 | 0.5936 | 0.4687 | 0.4646 | 0.4625 |
| 0:19:30 | 0.3939 | 0.3607 | 0.3687 | 0.6094 | 0.606 | 0.5906 | 0.4714 | 0.4669 | 0.4642 |
| 0:20:00 | 0.3965 | 0.3611 | 0.3682 | 0.6125 | 0.606 | 0.5953 | 0.4715 | 0.4693 | 0.4667 |
| 0:20:30 | 0.3985 | 0.3599 | 0.3675 | 0.6191 | 0.6141 | 0.6012 | 0.4752 | 0.467 | 0.4681 |
| 0:21:00 | 0.3974 | 0.3602 | 0.3682 | 0.6194 | 0.6158 | 0.6021 | 0.4829 | 0.469 | 0.4693 |
| 0:21:30 | 0.3959 | 0.36 | 0.367 | 0.6235 | 0.6189 | 0.61 | 0.4895 | 0.4702 | 0.4713 |
| 0:22:00 | 0.3962 | 0.3586 | 0.3689 | 0.6228 | 0.6237 | 0.6108 | 0.4909 | 0.4728 | 0.4713 |
| 0:22:30 | 0.3949 | 0.3594 | 0.367 | 0.625 | 0.627 | 0.6125 | 0.4958 | 0.4765 | 0.4741 |
| 0:23:00 | 0.3944 | 0.3599 | 0.3664 | 0.6309 | 0.627 | 0.6145 | 0.4941 | 0.4766 | 0.4752 |
| 0:23:30 | 0.3956 | 0.3586 | 0.3646 | 0.6334 | 0.6313 | 0.6162 | 0.4924 | 0.4801 | 0.4757 |

| | | | | | | | | | |
|---|---|---|---|---|---|---|---|---|---|
| 0:24:00 | 0.3953 | 0.3576 | 0.3657 | 0.6367 | 0.6311 | 0.6186 | 0.4912 | 0.4787 | 0.4796 |
| 0:24:30 | 0.395 | 0.3583 | 0.3663 | 0.6381 | 0.6334 | 0.6229 | 0.4908 | 0.4806 | 0.4804 |
| 0:25:00 | 0.3946 | 0.3583 | 0.3665 | 0.6422 | 0.6376 | 0.6288 | 0.4909 | 0.4834 | 0.4841 |
| 0:25:30 | 0.3915 | 0.3583 | 0.365 | 0.6388 | 0.6377 | 0.6256 | 0.4935 | 0.487 | 0.4824 |
| 0:26:00 | 0.3917 | 0.3567 | 0.3648 | 0.6406 | 0.6407 | 0.6261 | 0.4938 | 0.4893 | 0.4838 |
| 0:26:30 | 0.3955 | 0.3575 | 0.3652 | 0.6483 | 0.6428 | 0.6288 | 0.4975 | 0.4864 | 0.4863 |
| 0:27:00 | 0.3952 | 0.3562 | 0.3637 | 0.6511 | 0.6446 | 0.6338 | 0.498 | 0.4871 | 0.4882 |
| 0:27:30 | 0.3929 | 0.3568 | 0.3656 | 0.6487 | 0.6467 | 0.6378 | 0.4998 | 0.4888 | 0.4886 |
| 0:28:00 | 0.3923 | 0.3564 | 0.3663 | 0.6507 | 0.6474 | 0.6407 | 0.5005 | 0.4915 | 0.4913 |
| 0:28:30 | 0.3916 | 0.3575 | 0.3641 | 0.6505 | 0.6488 | 0.6385 | 0.5005 | 0.4934 | 0.4917 |
| 0:29:00 | 0.3914 | 0.3563 | 0.3633 | 0.6519 | 0.6522 | 0.6395 | 0.5028 | 0.4969 | 0.4952 |
| 0:29:30 | 0.392 | 0.3564 | 0.3629 | 0.655 | 0.653 | 0.6416 | 0.5044 | 0.4994 | 0.4936 |
| 0:30:00 | 0.3905 | 0.3557 | 0.3626 | 0.6553 | 0.6538 | 0.641 | 0.5062 | 0.5014 | 0.4956 |
| 0:30:30 | 0.3922 | 0.3545 | 0.3622 | 0.6614 | 0.6549 | 0.6426 | 0.509 | 0.5015 | 0.4979 |
| 0:31:00 | 0.393 | 0.3538 | 0.3629 | 0.6636 | 0.6526 | 0.6441 | 0.5089 | 0.5008 | 0.5009 |
| 0:31:30 | 0.3914 | 0.3546 | 0.3636 | 0.6585 | 0.6577 | 0.6468 | 0.5101 | 0.5039 | 0.4984 |
| 0:32:00 | 0.391 | 0.3543 | 0.3635 | 0.659 | 0.6551 | 0.6453 | 0.5135 | 0.5048 | 0.5012 |
| 0:32:30 | 0.3894 | 0.3543 | 0.3624 | 0.6604 | 0.6571 | 0.6477 | 0.5137 | 0.5071 | 0.5031 |
| 0:33:00 | 0.39 | 0.3542 | 0.3612 | 0.6611 | 0.6589 | 0.645 | 0.5144 | 0.5086 | 0.5047 |
| 0:33:30 | 0.3922 | 0.3547 | 0.3599 | 0.6638 | 0.6594 | 0.6462 | 0.5182 | 0.5105 | 0.5075 |
| 0:34:00 | 0.3928 | 0.3515 | 0.362 | 0.6674 | 0.6584 | 0.6501 | 0.5203 | 0.5106 | 0.5083 |
| 0:34:30 | 0.3902 | 0.3516 | 0.3619 | 0.6663 | 0.6595 | 0.6538 | 0.5177 | 0.5132 | 0.5091 |
| 0:35:00 | 0.3904 | 0.3516 | 0.3633 | 0.6643 | 0.6626 | 0.657 | 0.5212 | 0.5127 | 0.5122 |
| 0:35:30 | 0.3887 | 0.3531 | 0.3608 | 0.663 | 0.6626 | 0.6554 | 0.52 | 0.5158 | 0.5133 |
| 0:36:00 | 0.3888 | 0.3537 | 0.3617 | 0.664 | 0.6601 | 0.6535 | 0.5228 | 0.5164 | 0.5126 |
| 0:36:30 | 0.3876 | 0.3537 | 0.3601 | 0.6628 | 0.6627 | 0.6511 | 0.5254 | 0.5187 | 0.5155 |
| 0:37:00 | 0.3903 | 0.3515 | 0.3589 | 0.67 | 0.6585 | 0.6532 | 0.5274 | 0.5201 | 0.5192 |
| 0:37:30 | 0.3885 | 0.3521 | 0.3609 | 0.6677 | 0.6626 | 0.6576 | 0.5289 | 0.5186 | 0.5183 |
| 0:38:00 | 0.3876 | 0.3528 | 0.3606 | 0.6632 | 0.663 | 0.6572 | 0.5302 | 0.5219 | 0.5196 |
| 0:38:30 | 0.3874 | 0.3526 | 0.3599 | 0.6636 | 0.6627 | 0.6559 | 0.5325 | 0.5234 | 0.5204 |
| 0:39:00 | 0.3873 | 0.3519 | 0.3595 | 0.6665 | 0.6662 | 0.6543 | 0.532 | 0.5286 | 0.5208 |
| 0:39:30 | 0.3886 | 0.3505 | 0.3572 | 0.6683 | 0.6616 | 0.6531 | 0.5381 | 0.5299 | 0.5231 |
| 0:40:00 | 0.3904 | 0.3506 | 0.3589 | 0.6696 | 0.6655 | 0.6564 | 0.5341 | 0.5277 | 0.5261 |
| 0:40:30 | 0.3884 | 0.3509 | 0.3592 | 0.6689 | 0.6631 | 0.6576 | 0.5374 | 0.53 | 0.5262 |
| 0:41:00 | 0.3861 | 0.3512 | 0.3614 | 0.667 | 0.6655 | 0.6589 | 0.5409 | 0.5315 | 0.5267 |
| 0:41:30 | 0.3867 | 0.3513 | 0.3596 | 0.6673 | 0.6662 | 0.6553 | 0.5416 | 0.5333 | 0.5284 |
| 0:42:00 | 0.3872 | 0.3505 | 0.3569 | 0.6677 | 0.6651 | 0.6548 | 0.5435 | 0.5347 | 0.529 |
| 0:42:30 | 0.3882 | 0.3494 | 0.358 | 0.671 | 0.665 | 0.6581 | 0.5437 | 0.5348 | 0.5339 |
| 0:43:00 | 0.3873 | 0.3491 | 0.3585 | 0.6702 | 0.6665 | 0.6591 | 0.5438 | 0.5352 | 0.5323 |
| 0:43:30 | 0.3866 | 0.35 | 0.3598 | 0.6661 | 0.6663 | 0.6628 | 0.5476 | 0.5371 | 0.5344 |
| 0:44:00 | 0.3869 | 0.3503 | 0.358 | 0.6664 | 0.6658 | 0.6562 | 0.5483 | 0.5411 | 0.5357 |
| 0:44:30 | 0.3876 | 0.3491 | 0.3564 | 0.6693 | 0.662 | 0.6552 | 0.5485 | 0.5381 | 0.5345 |
| 0:45:00 | 0.387 | 0.3485 | 0.3568 | 0.6707 | 0.6654 | 0.6552 | 0.5501 | 0.5406 | 0.5389 |
| 0:45:30 | 0.3853 | 0.3496 | 0.3587 | 0.667 | 0.6683 | 0.6602 | 0.5504 | 0.5411 | 0.5398 |
| 0:46:00 | 0.3847 | 0.3486 | 0.3581 | 0.6668 | 0.6669 | 0.658 | 0.5536 | 0.5487 | 0.54 |
| 0:46:30 | 0.3845 | 0.3493 | 0.3555 | 0.6688 | 0.6661 | 0.6551 | 0.553 | 0.548 | 0.5418 |
| 0:47:00 | 0.3869 | 0.3499 | 0.3551 | 0.6715 | 0.664 | 0.6535 | 0.5574 | 0.5466 | 0.5446 |
| 0:47:30 | 0.3862 | 0.3468 | 0.3559 | 0.6693 | 0.6654 | 0.6596 | 0.555 | 0.5462 | 0.5463 |
| 0:48:00 | 0.3844 | 0.3483 | 0.3563 | 0.6672 | 0.6664 | 0.661 | 0.5597 | 0.5499 | 0.5457 |
| 0:48:30 | 0.3848 | 0.3484 | 0.3541 | 0.6668 | 0.666 | 0.6559 | 0.5608 | 0.5561 | 0.5467 |
| 0:49:00 | 0.3861 | 0.3465 | 0.3542 | 0.6726 | 0.6638 | 0.6543 | 0.5647 | 0.5523 | 0.5468 |
| 0:49:30 | 0.3839 | 0.3479 | 0.3566 | 0.6706 | 0.6663 | 0.6593 | 0.5621 | 0.5527 | 0.5471 |

| | | | | | | | | | |
|---|---|---|---|---|---|---|---|---|---|
| 0:50:00 | 0.3822 | 0.3484 | 0.3541 | 0.6665 | 0.6655 | 0.6562 | 0.5674 | 0.558 | 0.5496 |
| 0:50:30 | 0.3829 | 0.3466 | 0.3547 | 0.669 | 0.6636 | 0.6592 | 0.5674 | 0.5586 | 0.5507 |
| 0:51:00 | 0.3852 | 0.3473 | 0.3525 | 0.6725 | 0.6652 | 0.6564 | 0.5691 | 0.5573 | 0.5527 |
| 0:51:30 | 0.3842 | 0.3454 | 0.3544 | 0.6669 | 0.6642 | 0.6576 | 0.5689 | 0.5579 | 0.5548 |
| 0:52:00 | 0.383 | 0.3459 | 0.3561 | 0.6676 | 0.6657 | 0.6597 | 0.5676 | 0.5598 | 0.5545 |
| 0:52:30 | 0.3811 | 0.3466 | 0.3532 | 0.6629 | 0.668 | 0.6563 | 0.5692 | 0.564 | 0.5526 |
| 0:53:00 | 0.3818 | 0.3471 | 0.3527 | 0.6653 | 0.6691 | 0.6556 | 0.5709 | 0.5651 | 0.5583 |
| 0:53:30 | 0.3845 | 0.3455 | 0.3519 | 0.6729 | 0.6653 | 0.6529 | 0.5698 | 0.5628 | 0.5573 |
| 0:54:00 | 0.3825 | 0.3452 | 0.3517 | 0.6734 | 0.6659 | 0.6563 | 0.5746 | 0.5634 | 0.5609 |
| 0:54:30 | 0.3827 | 0.3462 | 0.3532 | 0.6708 | 0.6675 | 0.6579 | 0.5729 | 0.5632 | 0.5613 |
| 0:55:00 | 0.381 | 0.346 | 0.3549 | 0.6646 | 0.6671 | 0.6615 | 0.5758 | 0.5667 | 0.5616 |
| 0:55:30 | 0.3809 | 0.3458 | 0.3512 | 0.6648 | 0.6655 | 0.6598 | 0.576 | 0.5717 | 0.5629 |
| 0:56:00 | 0.3821 | 0.3451 | 0.3513 | 0.6666 | 0.6647 | 0.6565 | 0.5791 | 0.5711 | 0.5641 |
| 0:56:30 | 0.3839 | 0.3447 | 0.3512 | 0.6698 | 0.6622 | 0.655 | 0.5773 | 0.5685 | 0.565 |
| 0:57:00 | 0.3813 | 0.3431 | 0.3516 | 0.6686 | 0.6663 | 0.6592 | 0.5792 | 0.5693 | 0.5671 |
| 0:57:30 | 0.3804 | 0.3453 | 0.3524 | 0.6687 | 0.6683 | 0.6598 | 0.5821 | 0.5707 | 0.5668 |
| 0:58:00 | 0.3792 | 0.3449 | 0.3536 | 0.6639 | 0.6658 | 0.6577 | 0.5823 | 0.5727 | 0.5686 |
| 0:58:30 | 0.3797 | 0.3446 | 0.3511 | 0.6638 | 0.6687 | 0.6566 | 0.5823 | 0.5784 | 0.5673 |
| 0:59:00 | 0.38 | 0.3439 | 0.3515 | 0.6684 | 0.6668 | 0.6563 | 0.5863 | 0.5769 | 0.5695 |
| 0:59:30 | 0.3808 | 0.344 | 0.352 | 0.6697 | 0.6657 | 0.6519 | 0.5846 | 0.578 | 0.5704 |
| 1:00:00 | 0.3822 | 0.3434 | 0.349 | 0.6694 | 0.6631 | 0.6555 | 0.5871 | 0.577 | 0.5714 |

**MTREC (µM $P_i$)**

| Time (hr:min:sec) | H2O Ctrl (Rep A) | H2O Ctrl (Rep B) | H2O Ctrl (Rep C) | +RNA (RepA) | +RNA (RepB) | +RNA (RepC) | -RNA (Rep A) | -RNA (Rep B) | -RNA (Rep C) |
|---|---|---|---|---|---|---|---|---|---|
| 0:00:00 | -5.514 | -7.058 | -5.293 | 12.660 | 12.925 | 11.337 | 7.234 | 4.852 | 4.720 |
| 0:00:30 | 7.808 | -6.881 | -4.632 | 15.792 | 16.012 | 14.910 | 8.072 | 4.543 | 5.602 |
| 0:01:00 | 7.014 | -6.970 | -4.985 | 17.733 | 18.086 | 17.203 | 9.396 | 5.999 | 6.617 |
| 0:01:30 | 7.058 | -7.808 | -6.043 | 20.512 | 20.644 | 18.306 | 10.498 | 6.793 | 7.102 |
| 0:02:00 | 8.072 | -8.999 | -6.617 | 23.952 | 22.011 | 20.820 | 10.278 | 6.396 | 7.984 |
| 0:02:30 | 7.190 | -8.116 | -5.646 | 26.511 | 25.187 | 23.114 | 12.616 | 7.014 | 8.513 |
| 0:03:00 | 6.176 | -8.293 | -6.043 | 27.569 | 27.790 | 26.026 | 12.263 | 8.205 | 9.352 |
| 0:03:30 | 6.793 | -8.161 | -6.925 | 29.113 | 30.260 | 26.555 | 11.998 | 9.925 | 8.558 |
| 0:04:00 | 7.411 | -8.381 | -7.455 | 33.304 | 31.936 | 28.187 | 11.910 | 9.837 | 10.366 |
| 0:04:30 | 6.881 | -9.396 | -6.749 | 35.421 | 33.613 | 31.760 | 12.042 | 9.616 | 10.498 |
| 0:05:00 | 5.955 | -8.866 | -6.087 | 37.097 | 37.097 | 33.701 | 12.086 | 9.837 | 11.292 |
| 0:05:30 | 5.911 | -9.087 | -6.970 | 38.906 | 38.950 | 36.127 | 13.233 | 11.954 | 12.086 |
| 0:06:00 | 5.999 | -9.440 | -7.764 | 41.332 | 41.332 | 37.759 | 14.027 | 12.925 | 12.969 |
| 0:06:30 | 6.573 | -9.704 | -7.278 | 44.685 | 43.405 | 39.347 | 14.645 | 12.660 | 13.498 |
| 0:07:00 | 5.779 | -10.190 | -7.234 | 45.699 | 45.434 | 42.126 | 15.174 | 12.660 | 13.586 |
| 0:07:30 | 5.734 | -10.146 | -6.970 | 47.111 | 48.390 | 43.714 | 16.233 | 14.645 | 14.380 |
| 0:08:00 | 6.131 | -9.925 | -8.205 | 50.287 | 49.890 | 45.876 | 17.115 | 16.101 | 15.571 |
| 0:08:30 | 6.970 | -10.190 | -8.293 | 52.845 | 51.213 | 46.758 | 17.468 | 15.836 | 15.968 |
| 0:09:00 | 5.734 | -9.969 | -7.852 | 55.315 | 54.566 | 50.199 | 18.262 | 15.704 | 16.674 |
| 0:09:30 | 4.102 | -10.190 | -7.543 | 56.595 | 55.933 | 52.316 | 18.880 | 16.586 | 17.159 |
| 0:10:00 | 4.808 | -10.322 | -8.381 | 58.403 | 59.021 | 53.286 | 20.247 | 18.041 | 18.130 |
| 0:10:30 | 5.117 | -10.675 | -8.955 | 61.756 | 60.653 | 55.139 | 21.041 | 18.041 | 18.924 |
| 0:11:00 | 5.514 | -11.866 | -8.249 | 63.344 | 62.197 | 57.697 | 21.438 | 18.835 | 19.453 |
| 0:11:30 | 3.926 | -11.601 | -7.808 | 64.402 | 65.417 | 59.506 | 21.923 | 20.203 | 20.026 |
| 0:12:00 | 4.014 | -10.940 | -9.440 | 66.387 | 66.652 | 60.521 | 23.247 | 20.512 | 20.600 |
| 0:12:30 | 4.014 | -11.116 | -9.087 | 67.931 | 67.755 | 61.976 | 24.305 | 21.747 | 21.570 |
| 0:13:00 | 4.676 | -11.822 | -9.484 | 70.975 | 69.519 | 64.535 | 24.879 | 21.879 | 21.967 |
| 0:13:30 | 4.632 | -12.263 | -8.866 | 73.180 | 70.931 | 66.873 | 25.187 | 21.262 | 22.188 |
| 0:14:00 | 3.000 | -11.734 | -8.778 | 73.930 | 73.048 | 68.372 | 25.937 | 22.585 | 23.026 |
| 0:14:30 | 3.044 | -12.439 | -9.219 | 74.504 | 75.474 | 69.519 | 26.820 | 23.555 | 22.673 |
| 0:15:00 | 3.529 | -12.483 | -9.484 | 77.636 | 76.798 | 70.798 | 27.261 | 24.041 | 23.555 |
| 0:15:30 | 4.058 | -12.925 | -9.528 | 79.885 | 78.827 | 73.136 | 27.746 | 23.776 | 25.629 |
| 0:16:00 | 3.308 | -12.395 | -8.866 | 79.621 | 82.753 | 77.327 | 28.628 | 25.099 | 26.114 |
| 0:16:30 | 2.558 | -12.660 | -9.484 | 82.885 | 84.517 | 79.268 | 29.554 | 27.658 | 27.393 |
| 0:17:00 | 2.470 | -12.704 | -10.190 | 85.708 | 86.723 | 80.371 | 30.437 | 29.025 | 27.878 |
| 0:17:30 | 3.661 | -13.057 | -10.234 | 89.810 | 88.884 | 81.826 | 31.187 | 29.025 | 28.496 |
| 0:18:00 | 3.264 | -13.145 | -9.749 | 92.898 | 90.340 | 85.179 | 32.863 | 30.481 | 29.599 |
| 0:18:30 | 2.823 | -13.939 | -9.219 | 93.030 | 92.192 | 87.296 | 33.436 | 30.437 | 30.878 |
| 0:19:00 | 2.117 | -13.189 | -9.572 | 94.618 | 92.942 | 89.060 | 33.966 | 32.157 | 31.231 |
| 0:19:30 | 0.970 | -13.674 | -10.146 | 96.030 | 94.530 | 87.737 | 35.157 | 33.172 | 31.981 |
| 0:20:00 | 2.117 | -13.498 | -10.366 | 97.397 | 94.530 | 89.810 | 35.201 | 34.230 | 33.083 |
| 0:20:30 | 3.000 | -14.027 | -10.675 | 100.309 | 98.103 | 92.413 | 36.833 | 33.216 | 33.701 |
| 0:21:00 | 2.514 | -13.895 | -10.366 | 100.441 | 98.853 | 92.810 | 40.229 | 34.098 | 34.230 |
| 0:21:30 | 1.853 | -13.983 | -10.895 | 102.250 | 100.221 | 96.295 | 43.141 | 34.627 | 35.112 |
| 0:22:00 | 1.985 | -14.601 | -10.057 | 101.941 | 102.338 | 96.648 | 43.758 | 35.774 | 35.112 |
| 0:22:30 | 1.412 | -14.248 | -10.895 | 102.911 | 103.794 | 97.397 | 45.920 | 37.406 | 36.348 |
| 0:23:00 | 1.191 | -14.027 | -11.160 | 105.514 | 103.794 | 98.280 | 45.170 | 37.450 | 36.833 |
| 0:23:30 | 1.720 | -14.601 | -11.954 | 106.617 | 105.690 | 99.030 | 44.420 | 38.994 | 37.053 |
| 0:24:00 | 1.588 | -15.042 | -11.469 | 108.072 | 105.602 | 100.088 | 43.891 | 38.377 | 38.774 |
| 0:24:30 | 1.456 | -14.733 | -11.204 | 108.690 | 106.617 | 101.985 | 43.714 | 39.215 | 39.127 |
| 0:25:00 | 1.279 | -14.733 | -11.116 | 110.498 | 108.469 | 104.588 | 43.758 | 40.450 | 40.759 |
| 0:25:30 | -0.088 | -14.733 | -11.778 | 108.999 | 108.513 | 103.176 | 44.905 | 42.038 | 40.009 |
| 0:26:00 | 0.000 | -15.439 | -11.866 | 109.793 | 109.837 | 103.397 | 45.037 | 43.052 | 40.626 |

| | | | | | | | | | |
|---|---|---|---|---|---|---|---|---|---|
| 0:26:30 | 1.676 | -15.086 | -11.689 | 113.189 | 110.763 | 104.588 | 46.670 | 41.773 | 41.729 |
| 0:27:00 | 1.544 | -15.659 | -12.351 | 114.424 | 111.557 | 106.793 | 46.890 | 42.082 | 42.567 |
| 0:27:30 | 0.529 | -15.395 | -11.513 | 113.366 | 112.483 | 108.558 | 47.684 | 42.832 | 42.744 |
| 0:28:00 | 0.265 | -15.571 | -11.204 | 114.248 | 112.792 | 109.837 | 47.993 | 44.023 | 43.935 |
| 0:28:30 | -0.044 | -15.086 | -12.175 | 114.160 | 113.410 | 108.866 | 47.993 | 44.861 | 44.111 |
| 0:29:00 | -0.132 | -15.615 | -12.528 | 114.777 | 114.910 | 109.307 | 49.007 | 46.405 | 45.655 |
| 0:29:30 | 0.132 | -15.571 | -12.704 | 116.145 | 115.262 | 110.234 | 49.713 | 47.508 | 44.949 |
| 0:30:00 | -0.529 | -15.880 | -12.836 | 116.277 | 115.615 | 109.969 | 50.507 | 48.390 | 45.831 |
| 0:30:30 | 0.221 | -16.409 | -13.013 | 118.968 | 116.101 | 110.675 | 51.742 | 48.434 | 46.846 |
| 0:31:00 | 0.573 | -16.718 | -12.704 | 119.938 | 115.086 | 111.337 | 51.698 | 48.125 | 48.169 |
| 0:31:30 | -0.132 | -16.365 | -12.395 | 117.689 | 117.336 | 112.528 | 52.228 | 49.493 | 47.067 |
| 0:32:00 | -0.309 | -16.498 | -12.439 | 117.909 | 116.189 | 111.866 | 53.727 | 49.890 | 48.302 |
| 0:32:30 | -1.015 | -16.498 | -12.925 | 118.527 | 117.071 | 112.925 | 53.816 | 50.904 | 49.140 |
| 0:33:00 | -0.750 | -16.542 | -13.454 | 118.835 | 117.865 | 111.734 | 54.124 | 51.566 | 49.846 |
| 0:33:30 | 0.221 | -16.321 | -14.027 | 120.026 | 118.086 | 112.263 | 55.801 | 52.404 | 51.081 |
| 0:34:00 | 0.485 | -17.733 | -13.101 | 121.614 | 117.644 | 113.983 | 56.727 | 52.448 | 51.434 |
| 0:34:30 | -0.662 | -17.689 | -13.145 | 121.129 | 118.130 | 115.615 | 55.580 | 53.595 | 51.787 |
| 0:35:00 | -0.573 | -17.689 | -12.528 | 120.247 | 119.497 | 117.027 | 57.124 | 53.375 | 53.154 |
| 0:35:30 | -1.323 | -17.027 | -13.630 | 119.674 | 119.497 | 116.321 | 56.595 | 54.742 | 53.639 |
| 0:36:00 | -1.279 | -16.762 | -13.233 | 120.115 | 118.394 | 115.483 | 57.830 | 55.007 | 53.330 |
| 0:36:30 | -1.809 | -16.762 | -13.939 | 119.585 | 119.541 | 114.424 | 58.977 | 56.021 | 54.610 |
| 0:37:00 | -0.618 | -17.733 | -14.468 | 122.761 | 117.689 | 115.351 | 59.859 | 56.639 | 56.242 |
| 0:37:30 | -1.412 | -17.468 | -13.586 | 121.747 | 119.497 | 117.292 | 60.521 | 55.977 | 55.845 |
| 0:38:00 | -1.809 | -17.159 | -13.719 | 119.762 | 119.674 | 117.115 | 61.094 | 57.433 | 56.418 |
| 0:38:30 | -1.897 | -17.247 | -14.027 | 119.938 | 119.541 | 116.542 | 62.109 | 58.094 | 56.771 |
| 0:39:00 | -1.941 | -17.556 | -14.204 | 121.217 | 121.085 | 115.836 | 61.888 | 60.388 | 56.948 |
| 0:39:30 | -1.367 | -18.174 | -15.218 | 122.011 | 119.056 | 115.307 | 64.579 | 60.962 | 57.962 |
| 0:40:00 | -0.573 | -18.130 | -14.468 | 122.585 | 120.776 | 116.762 | 62.814 | 59.991 | 59.285 |
| 0:40:30 | -1.456 | -17.997 | -14.336 | 122.276 | 119.718 | 117.292 | 64.270 | 61.006 | 59.330 |
| 0:41:00 | -2.470 | -17.865 | -13.366 | 121.438 | 120.776 | 117.865 | 65.814 | 61.667 | 59.550 |
| 0:41:30 | -2.206 | -17.821 | -14.160 | 121.570 | 121.085 | 116.277 | 66.123 | 62.461 | 60.300 |
| 0:42:00 | -1.985 | -18.174 | -15.351 | 121.747 | 120.600 | 116.056 | 66.961 | 63.079 | 60.565 |
| 0:42:30 | -1.544 | -18.659 | -14.865 | 123.202 | 120.556 | 117.512 | 67.049 | 63.123 | 62.726 |
| 0:43:00 | -1.941 | -18.791 | -14.645 | 122.850 | 121.217 | 117.953 | 67.093 | 63.300 | 62.020 |
| 0:43:30 | -2.250 | -18.394 | -14.071 | 121.041 | 121.129 | 119.585 | 68.769 | 64.138 | 62.947 |
| 0:44:00 | -2.117 | -18.262 | -14.865 | 121.173 | 120.909 | 116.674 | 69.078 | 65.902 | 63.520 |
| 0:44:30 | -1.809 | -18.791 | -15.571 | 122.453 | 119.232 | 116.233 | 69.166 | 64.579 | 62.991 |
| 0:45:00 | -2.073 | -19.056 | -15.395 | 123.070 | 120.732 | 116.233 | 69.872 | 65.682 | 64.932 |
| 0:45:30 | -2.823 | -18.571 | -14.557 | 121.438 | 122.011 | 118.438 | 70.004 | 65.902 | 65.329 |
| 0:46:00 | -3.088 | -19.012 | -14.821 | 121.350 | 121.394 | 117.468 | 71.416 | 69.255 | 65.417 |
| 0:46:30 | -3.176 | -18.703 | -15.968 | 122.232 | 121.041 | 116.189 | 71.151 | 68.946 | 66.211 |
| 0:47:00 | -2.117 | -18.438 | -16.145 | 123.423 | 120.115 | 115.483 | 73.092 | 68.328 | 67.446 |
| 0:47:30 | -2.426 | -19.806 | -15.792 | 122.453 | 120.732 | 118.174 | 72.034 | 68.152 | 68.196 |
| 0:48:00 | -3.220 | -19.144 | -15.615 | 121.526 | 121.173 | 118.791 | 74.107 | 69.784 | 67.931 |
| 0:48:30 | -3.044 | -19.100 | -16.586 | 121.350 | 120.997 | 116.542 | 74.592 | 72.519 | 68.372 |
| 0:49:00 | -2.470 | -19.938 | -16.542 | 123.908 | 120.026 | 115.836 | 76.312 | 70.843 | 68.416 |
| 0:49:30 | -3.441 | -19.321 | -15.483 | 123.026 | 121.129 | 118.041 | 75.165 | 71.019 | 68.549 |
| 0:50:00 | -4.191 | -19.100 | -16.586 | 121.217 | 120.776 | 116.674 | 77.503 | 73.357 | 69.652 |
| 0:50:30 | -3.882 | -19.894 | -16.321 | 122.320 | 119.938 | 117.997 | 77.503 | 73.622 | 70.137 |
| 0:51:00 | -2.867 | -19.585 | -17.292 | 123.864 | 120.644 | 116.762 | 78.253 | 73.048 | 71.019 |
| 0:51:30 | -3.308 | -20.423 | -16.453 | 121.394 | 120.203 | 117.292 | 78.165 | 73.313 | 71.945 |
| 0:52:00 | -3.838 | -20.203 | -15.704 | 121.703 | 120.865 | 118.218 | 77.592 | 74.151 | 71.813 |
| 0:52:30 | -4.676 | -19.894 | -16.983 | 119.629 | 121.879 | 116.718 | 78.297 | 76.004 | 70.975 |
| 0:53:00 | -4.367 | -19.674 | -17.203 | 120.688 | 122.364 | 116.409 | 79.047 | 76.489 | 73.489 |
| 0:53:30 | -3.176 | -20.379 | -17.556 | 124.041 | 120.688 | 115.218 | 78.562 | 75.474 | 73.048 |
| 0:54:00 | -4.058 | -20.512 | -17.644 | 124.261 | 120.953 | 116.718 | 80.679 | 75.739 | 74.636 |
| 0:54:30 | -3.970 | -20.071 | -16.983 | 123.114 | 121.659 | 117.424 | 79.929 | 75.651 | 74.813 |

| | | | | | | | | | |
|---|---|---|---|---|---|---|---|---|---|
| 0:55:00 | -4.720 | -20.159 | -16.233 | 120.379 | 121.482 | 119.012 | 81.209 | 77.195 | 74.945 |
| 0:55:30 | -4.764 | -20.247 | -17.865 | 120.468 | 120.776 | 118.262 | 81.297 | 79.400 | 75.518 |
| 0:56:00 | -4.235 | -20.556 | -17.821 | 121.262 | 120.423 | 116.806 | 82.664 | 79.135 | 76.048 |
| 0:56:30 | -3.441 | -20.732 | -17.865 | 122.673 | 119.321 | 116.145 | 81.870 | 77.989 | 76.445 |
| 0:57:00 | -4.588 | -21.438 | -17.689 | 122.144 | 121.129 | 117.997 | 82.708 | 78.341 | 77.371 |
| 0:57:30 | -4.985 | -20.468 | -17.336 | 122.188 | 122.011 | 118.262 | 83.988 | 78.959 | 77.239 |
| 0:58:00 | -5.514 | -20.644 | -16.806 | 120.071 | 120.909 | 117.336 | 84.076 | 79.841 | 78.033 |
| 0:58:30 | -5.293 | -20.776 | -17.909 | 120.026 | 122.188 | 116.850 | 84.076 | 82.356 | 77.459 |
| 0:59:00 | -5.161 | -21.085 | -17.733 | 122.056 | 121.350 | 116.718 | 85.840 | 81.694 | 78.430 |
| 0:59:30 | -4.808 | -21.041 | -17.512 | 122.629 | 120.865 | 114.777 | 85.090 | 82.179 | 78.827 |
| 1:00:00 | -4.191 | -21.306 | -18.835 | 122.497 | 119.718 | 116.365 | 86.193 | 81.738 | 79.268 |

**MTREC[mut] (Abs 360 nm)**

| Time (hr:min:sec) | $H_2O$ Ctrl (Rep A) | $H_2O$ Ctrl (Rep B) | $H_2O$ Ctrl (Rep C) | +RNA (RepA) | +RNA (RepB) | +RNA (RepC) | -RNA (Rep A) | -RNA (Rep B) | -RNA (Rep C) |
|---|---|---|---|---|---|---|---|---|---|
| 0:00:00 | 0.3812 | 0.3834 | 0.3838 | 0.4112 | 0.4123 | 0.4075 | 0.4296 | 0.4339 | 0.4374 |
| 0:00:30 | 0.3828 | 0.384 | 0.3855 | 0.4171 | 0.4168 | 0.4119 | 0.451 | 0.4561 | 0.4568 |
| 0:01:00 | 0.3819 | 0.3829 | 0.3865 | 0.4232 | 0.4214 | 0.4146 | 0.4678 | 0.4748 | 0.4752 |
| 0:01:30 | 0.3802 | 0.3827 | 0.3865 | 0.4235 | 0.4256 | 0.4195 | 0.4836 | 0.4927 | 0.491 |
| 0:02:00 | 0.3811 | 0.382 | 0.3831 | 0.4314 | 0.4291 | 0.4212 | 0.5006 | 0.5108 | 0.5034 |
| 0:02:30 | 0.3805 | 0.3818 | 0.3834 | 0.4357 | 0.4304 | 0.4244 | 0.5164 | 0.526 | 0.5198 |
| 0:03:00 | 0.3814 | 0.3817 | 0.3852 | 0.4378 | 0.4344 | 0.4291 | 0.5293 | 0.5414 | 0.5348 |
| 0:03:30 | 0.3797 | 0.3807 | 0.3846 | 0.4393 | 0.4395 | 0.4326 | 0.5458 | 0.5555 | 0.5505 |
| 0:04:00 | 0.3786 | 0.3805 | 0.3842 | 0.4426 | 0.4404 | 0.4346 | 0.5561 | 0.5709 | 0.5629 |
| 0:04:30 | 0.3787 | 0.3806 | 0.3832 | 0.4471 | 0.4456 | 0.4384 | 0.5725 | 0.5859 | 0.5744 |
| 0:05:00 | 0.379 | 0.3805 | 0.3831 | 0.4481 | 0.4466 | 0.4404 | 0.5832 | 0.5953 | 0.5831 |
| 0:05:30 | 0.3774 | 0.3793 | 0.3835 | 0.4514 | 0.4501 | 0.4444 | 0.5928 | 0.6086 | 0.5951 |
| 0:06:00 | 0.3786 | 0.3795 | 0.3838 | 0.4574 | 0.4555 | 0.4466 | 0.604 | 0.6179 | 0.61 |
| 0:06:30 | 0.3779 | 0.3789 | 0.3849 | 0.4631 | 0.4576 | 0.4494 | 0.6135 | 0.6268 | 0.6174 |
| 0:07:00 | 0.3782 | 0.3784 | 0.3838 | 0.4645 | 0.4622 | 0.4528 | 0.6235 | 0.6348 | 0.6248 |
| 0:07:30 | 0.3782 | 0.3791 | 0.3835 | 0.4688 | 0.4648 | 0.4567 | 0.6342 | 0.6462 | 0.6357 |
| 0:08:00 | 0.3769 | 0.3786 | 0.3817 | 0.4742 | 0.4696 | 0.4591 | 0.6426 | 0.6517 | 0.6454 |
| 0:08:30 | 0.3769 | 0.3785 | 0.3834 | 0.4755 | 0.4729 | 0.4649 | 0.6446 | 0.6594 | 0.651 |
| 0:09:00 | 0.377 | 0.376 | 0.3804 | 0.4785 | 0.477 | 0.4673 | 0.6554 | 0.6635 | 0.6599 |
| 0:09:30 | 0.3774 | 0.3772 | 0.3822 | 0.4829 | 0.4787 | 0.471 | 0.6631 | 0.674 | 0.6623 |
| 0:10:00 | 0.3758 | 0.3757 | 0.3817 | 0.4872 | 0.4822 | 0.4746 | 0.6683 | 0.6777 | 0.6728 |
| 0:10:30 | 0.376 | 0.3752 | 0.3816 | 0.4915 | 0.4858 | 0.4778 | 0.6719 | 0.6822 | 0.6789 |
| 0:11:00 | 0.3756 | 0.3754 | 0.3804 | 0.4957 | 0.4904 | 0.4822 | 0.676 | 0.6851 | 0.682 |
| 0:11:30 | 0.3763 | 0.3755 | 0.3796 | 0.4966 | 0.4913 | 0.483 | 0.6838 | 0.6851 | 0.6869 |
| 0:12:00 | 0.3776 | 0.3741 | 0.3803 | 0.5012 | 0.4933 | 0.4885 | 0.6868 | 0.6891 | 0.6918 |
| 0:12:30 | 0.3752 | 0.3742 | 0.3794 | 0.5037 | 0.497 | 0.491 | 0.6878 | 0.6898 | 0.6966 |
| 0:13:00 | 0.3744 | 0.3744 | 0.3802 | 0.505 | 0.5042 | 0.4941 | 0.693 | 0.6945 | 0.699 |
| 0:13:30 | 0.3745 | 0.374 | 0.3804 | 0.5099 | 0.5113 | 0.4963 | 0.6947 | 0.6938 | 0.6967 |
| 0:14:00 | 0.3766 | 0.3746 | 0.3798 | 0.513 | 0.5157 | 0.5013 | 0.6999 | 0.6955 | 0.7025 |
| 0:14:30 | 0.3753 | 0.3736 | 0.3801 | 0.5218 | 0.521 | 0.5042 | 0.7017 | 0.6965 | 0.7015 |
| 0:15:00 | 0.3742 | 0.3742 | 0.381 | 0.5237 | 0.5236 | 0.5057 | 0.7005 | 0.7004 | 0.7052 |
| 0:15:30 | 0.3738 | 0.374 | 0.3805 | 0.5265 | 0.5272 | 0.5096 | 0.6974 | 0.6994 | 0.7054 |
| 0:16:00 | 0.3736 | 0.3731 | 0.3793 | 0.5317 | 0.5362 | 0.5171 | 0.6981 | 0.6995 | 0.7054 |
| 0:16:30 | 0.3762 | 0.372 | 0.3782 | 0.5333 | 0.5347 | 0.525 | 0.7019 | 0.6973 | 0.7074 |
| 0:17:00 | 0.3774 | 0.3738 | 0.3792 | 0.539 | 0.5385 | 0.5294 | 0.6972 | 0.6993 | 0.708 |
| 0:17:30 | 0.373 | 0.3734 | 0.379 | 0.5445 | 0.5426 | 0.5332 | 0.6997 | 0.7018 | 0.7111 |
| 0:18:00 | 0.3732 | 0.3728 | 0.378 | 0.5463 | 0.5457 | 0.5376 | 0.6976 | 0.7033 | 0.7084 |
| 0:18:30 | 0.3733 | 0.3708 | 0.3777 | 0.5497 | 0.5533 | 0.5394 | 0.6973 | 0.7047 | 0.7102 |
| 0:19:00 | 0.3742 | 0.3719 | 0.3794 | 0.5525 | 0.5589 | 0.5426 | 0.7032 | 0.7077 | 0.7095 |
| 0:19:30 | 0.3743 | 0.3722 | 0.3798 | 0.5564 | 0.561 | 0.5454 | 0.7008 | 0.7038 | 0.7081 |
| 0:20:00 | 0.3736 | 0.3732 | 0.3784 | 0.5596 | 0.5581 | 0.5494 | 0.7019 | 0.7031 | 0.7139 |
| 0:20:30 | 0.3724 | 0.371 | 0.3778 | 0.5632 | 0.5644 | 0.5525 | 0.7033 | 0.704 | 0.7124 |
| 0:21:00 | 0.373 | 0.3712 | 0.3783 | 0.5688 | 0.5681 | 0.5569 | 0.7008 | 0.7055 | 0.7108 |
| 0:21:30 | 0.3719 | 0.373 | 0.3761 | 0.569 | 0.5724 | 0.5583 | 0.7015 | 0.7049 | 0.7109 |
| 0:22:00 | 0.3731 | 0.3718 | 0.3773 | 0.5731 | 0.5694 | 0.5618 | 0.7041 | 0.7072 | 0.7077 |
| 0:22:30 | 0.3746 | 0.3724 | 0.3777 | 0.5737 | 0.5778 | 0.5639 | 0.7046 | 0.7075 | 0.7127 |
| 0:23:00 | 0.3731 | 0.3713 | 0.3787 | 0.5788 | 0.5796 | 0.5685 | 0.7033 | 0.7043 | 0.7101 |
| 0:23:30 | 0.3726 | 0.3717 | 0.377 | 0.5823 | 0.5826 | 0.5702 | 0.7042 | 0.7039 | 0.7141 |

| | | | | | | | | | |
|---|---|---|---|---|---|---|---|---|---|
| 0:24:00 | 0.3724 | 0.3722 | 0.3791 | 0.5841 | 0.5857 | 0.5692 | 0.7041 | 0.7037 | 0.71 |
| 0:24:30 | 0.3733 | 0.372 | 0.3778 | 0.5877 | 0.5895 | 0.574 | 0.7022 | 0.7053 | 0.7102 |
| 0:25:00 | 0.3712 | 0.3712 | 0.379 | 0.5893 | 0.5931 | 0.5769 | 0.7061 | 0.7049 | 0.712 |
| 0:25:30 | 0.3728 | 0.3711 | 0.3783 | 0.5926 | 0.5928 | 0.5814 | 0.7073 | 0.707 | 0.7118 |
| 0:26:00 | 0.3724 | 0.3715 | 0.3786 | 0.5957 | 0.5986 | 0.5853 | 0.7046 | 0.7044 | 0.7155 |
| 0:26:30 | 0.3703 | 0.3714 | 0.3789 | 0.5982 | 0.5986 | 0.5876 | 0.7026 | 0.7054 | 0.7144 |
| 0:27:00 | 0.3698 | 0.3724 | 0.3772 | 0.6028 | 0.6007 | 0.5893 | 0.7028 | 0.7048 | 0.7143 |
| 0:27:30 | 0.371 | 0.3703 | 0.3766 | 0.6026 | 0.6047 | 0.5924 | 0.7026 | 0.7066 | 0.7121 |
| 0:28:00 | 0.3707 | 0.371 | 0.3776 | 0.6072 | 0.6048 | 0.5946 | 0.7039 | 0.7062 | 0.7128 |
| 0:28:30 | 0.3725 | 0.3704 | 0.3763 | 0.6086 | 0.6075 | 0.5996 | 0.707 | 0.7038 | 0.7113 |
| 0:29:00 | 0.3716 | 0.3712 | 0.3763 | 0.6129 | 0.6104 | 0.6013 | 0.7045 | 0.7068 | 0.7141 |
| 0:29:30 | 0.3713 | 0.3714 | 0.3766 | 0.6137 | 0.6127 | 0.6029 | 0.7032 | 0.7054 | 0.7132 |
| 0:30:00 | 0.3706 | 0.3702 | 0.3787 | 0.6159 | 0.6156 | 0.6038 | 0.7039 | 0.7041 | 0.711 |
| 0:30:30 | 0.3704 | 0.3712 | 0.3772 | 0.6166 | 0.6165 | 0.605 | 0.702 | 0.7047 | 0.7152 |
| 0:31:00 | 0.3698 | 0.371 | 0.3772 | 0.6173 | 0.6196 | 0.6059 | 0.7006 | 0.7057 | 0.7133 |
| 0:31:30 | 0.3686 | 0.3703 | 0.3754 | 0.6203 | 0.6229 | 0.6109 | 0.6998 | 0.7051 | 0.7109 |
| 0:32:00 | 0.37 | 0.3711 | 0.3743 | 0.6205 | 0.6233 | 0.6124 | 0.7008 | 0.7025 | 0.71 |
| 0:32:30 | 0.3714 | 0.3714 | 0.3758 | 0.6241 | 0.6248 | 0.6137 | 0.7049 | 0.7055 | 0.7106 |
| 0:33:00 | 0.3705 | 0.3696 | 0.3773 | 0.6275 | 0.6269 | 0.6159 | 0.7015 | 0.7038 | 0.7117 |
| 0:33:30 | 0.3692 | 0.3708 | 0.3772 | 0.6287 | 0.6271 | 0.616 | 0.699 | 0.7038 | 0.7163 |
| 0:34:00 | 0.3691 | 0.3704 | 0.3753 | 0.6267 | 0.6304 | 0.6192 | 0.7037 | 0.7035 | 0.712 |
| 0:34:30 | 0.3689 | 0.3693 | 0.3745 | 0.6312 | 0.6338 | 0.6212 | 0.7036 | 0.7021 | 0.7093 |
| 0:35:00 | 0.3685 | 0.3687 | 0.3734 | 0.6335 | 0.6358 | 0.6222 | 0.7019 | 0.7038 | 0.71 |
| 0:35:30 | 0.3688 | 0.3696 | 0.375 | 0.6303 | 0.6335 | 0.6254 | 0.7035 | 0.7033 | 0.709 |
| 0:36:00 | 0.3706 | 0.3683 | 0.3744 | 0.6362 | 0.6339 | 0.6244 | 0.7044 | 0.7009 | 0.7096 |
| 0:36:30 | 0.3704 | 0.3694 | 0.3743 | 0.6378 | 0.6378 | 0.6253 | 0.7038 | 0.7029 | 0.7144 |
| 0:37:00 | 0.3684 | 0.3693 | 0.3753 | 0.637 | 0.6408 | 0.6296 | 0.7004 | 0.702 | 0.7131 |
| 0:37:30 | 0.3675 | 0.3696 | 0.3737 | 0.638 | 0.6424 | 0.632 | 0.6996 | 0.7022 | 0.7127 |
| 0:38:00 | 0.3688 | 0.3686 | 0.3726 | 0.639 | 0.6411 | 0.6314 | 0.7006 | 0.7024 | 0.711 |
| 0:38:30 | 0.369 | 0.3688 | 0.3733 | 0.6412 | 0.6427 | 0.6328 | 0.7053 | 0.7003 | 0.7115 |
| 0:39:00 | 0.3692 | 0.3682 | 0.3741 | 0.6429 | 0.6416 | 0.6318 | 0.7023 | 0.7009 | 0.7111 |
| 0:39:30 | 0.3691 | 0.3689 | 0.3765 | 0.6431 | 0.6431 | 0.6335 | 0.7029 | 0.702 | 0.7118 |
| 0:40:00 | 0.3678 | 0.3682 | 0.3756 | 0.6444 | 0.6458 | 0.6341 | 0.7001 | 0.7013 | 0.7118 |
| 0:40:30 | 0.3671 | 0.3681 | 0.3737 | 0.647 | 0.6491 | 0.6357 | 0.7004 | 0.7034 | 0.7127 |
| 0:41:00 | 0.3684 | 0.3683 | 0.3726 | 0.6465 | 0.6482 | 0.6378 | 0.7001 | 0.7009 | 0.7133 |
| 0:41:30 | 0.3709 | 0.3682 | 0.3729 | 0.6452 | 0.6483 | 0.6379 | 0.7007 | 0.6993 | 0.7098 |
| 0:42:00 | 0.3695 | 0.3683 | 0.3727 | 0.647 | 0.6495 | 0.6373 | 0.7 | 0.7017 | 0.7144 |
| 0:42:30 | 0.3671 | 0.3672 | 0.374 | 0.6486 | 0.6476 | 0.6404 | 0.6972 | 0.7018 | 0.7115 |
| 0:43:00 | 0.3689 | 0.3668 | 0.3724 | 0.6495 | 0.6529 | 0.6404 | 0.6986 | 0.7005 | 0.706 |
| 0:43:30 | 0.3671 | 0.3655 | 0.3724 | 0.6487 | 0.6516 | 0.6402 | 0.6998 | 0.7017 | 0.7066 |
| 0:44:00 | 0.3696 | 0.3658 | 0.3719 | 0.6501 | 0.6523 | 0.6392 | 0.7037 | 0.7018 | 0.7082 |
| 0:44:30 | 0.3681 | 0.367 | 0.3734 | 0.6493 | 0.6531 | 0.6424 | 0.6999 | 0.7017 | 0.7103 |
| 0:45:00 | 0.3675 | 0.3667 | 0.3706 | 0.6525 | 0.6554 | 0.6437 | 0.7003 | 0.6998 | 0.7118 |
| 0:45:30 | 0.3672 | 0.3704 | 0.3707 | 0.6509 | 0.6563 | 0.6434 | 0.6972 | 0.7005 | 0.706 |
| 0:46:00 | 0.3688 | 0.3668 | 0.3718 | 0.6515 | 0.6505 | 0.642 | 0.6997 | 0.6998 | 0.7087 |
| 0:46:30 | 0.3678 | 0.3665 | 0.3703 | 0.6509 | 0.6537 | 0.6433 | 0.6993 | 0.6988 | 0.7095 |
| 0:47:00 | 0.3667 | 0.3666 | 0.3723 | 0.6524 | 0.6559 | 0.6434 | 0.699 | 0.6989 | 0.7102 |
| 0:47:30 | 0.3658 | 0.3665 | 0.3706 | 0.6502 | 0.6544 | 0.6487 | 0.6955 | 0.6985 | 0.7093 |
| 0:48:00 | 0.3668 | 0.3666 | 0.3698 | 0.6531 | 0.661 | 0.6468 | 0.6986 | 0.6979 | 0.7064 |
| 0:48:30 | 0.367 | 0.3659 | 0.3697 | 0.6526 | 0.6563 | 0.6442 | 0.702 | 0.6987 | 0.7045 |
| 0:49:00 | 0.3651 | 0.3637 | 0.3704 | 0.6517 | 0.6566 | 0.6439 | 0.6961 | 0.7 | 0.7085 |
| 0:49:30 | 0.3659 | 0.3663 | 0.3698 | 0.6533 | 0.6617 | 0.647 | 0.6968 | 0.7019 | 0.7095 |

| | | | | | | | | | |
|---|---|---|---|---|---|---|---|---|---|
| 0:50:00 | 0.3669 | 0.3639 | 0.3724 | 0.6533 | 0.6549 | 0.6473 | 0.6977 | 0.7017 | 0.7057 |
| 0:50:30 | 0.3663 | 0.3641 | 0.3692 | 0.6524 | 0.6569 | 0.6467 | 0.6972 | 0.6997 | 0.7098 |
| 0:51:00 | 0.365 | 0.3663 | 0.3702 | 0.6537 | 0.6559 | 0.6486 | 0.6983 | 0.7014 | 0.708 |
| 0:51:30 | 0.3643 | 0.3644 | 0.3699 | 0.6534 | 0.6603 | 0.6476 | 0.6979 | 0.697 | 0.7079 |
| 0:52:00 | 0.365 | 0.3639 | 0.37 | 0.6535 | 0.6575 | 0.6456 | 0.6964 | 0.698 | 0.7035 |
| 0:52:30 | 0.3669 | 0.3654 | 0.3685 | 0.6523 | 0.6541 | 0.6494 | 0.7004 | 0.7011 | 0.7072 |
| 0:53:00 | 0.3642 | 0.3644 | 0.368 | 0.653 | 0.658 | 0.6431 | 0.6978 | 0.6991 | 0.7088 |
| 0:53:30 | 0.3646 | 0.3636 | 0.3701 | 0.6567 | 0.6576 | 0.6453 | 0.6965 | 0.6962 | 0.7116 |
| 0:54:00 | 0.3622 | 0.3629 | 0.3681 | 0.6547 | 0.6596 | 0.6489 | 0.6942 | 0.6996 | 0.7063 |
| 0:54:30 | 0.3645 | 0.3635 | 0.3679 | 0.6566 | 0.6576 | 0.6458 | 0.6939 | 0.6971 | 0.7035 |
| 0:55:00 | 0.364 | 0.3645 | 0.3667 | 0.654 | 0.6573 | 0.6472 | 0.698 | 0.6977 | 0.7026 |
| 0:55:30 | 0.3648 | 0.3628 | 0.3675 | 0.6559 | 0.6573 | 0.6502 | 0.7013 | 0.6985 | 0.7065 |
| 0:56:00 | 0.3633 | 0.362 | 0.3677 | 0.654 | 0.6586 | 0.6463 | 0.6943 | 0.6967 | 0.7115 |
| 0:56:30 | 0.3634 | 0.3634 | 0.368 | 0.6554 | 0.6589 | 0.6468 | 0.6956 | 0.6971 | 0.7067 |
| 0:57:00 | 0.3627 | 0.3631 | 0.3665 | 0.6563 | 0.6583 | 0.6496 | 0.6958 | 0.6991 | 0.7055 |
| 0:57:30 | 0.3631 | 0.362 | 0.3673 | 0.6571 | 0.6601 | 0.6469 | 0.6954 | 0.698 | 0.7073 |
| 0:58:00 | 0.3635 | 0.3626 | 0.3654 | 0.6559 | 0.6574 | 0.6471 | 0.6969 | 0.6968 | 0.7063 |
| 0:58:30 | 0.3642 | 0.3623 | 0.366 | 0.6518 | 0.6564 | 0.6502 | 0.6974 | 0.6976 | 0.7028 |
| 0:59:00 | 0.3636 | 0.3621 | 0.3662 | 0.6556 | 0.66 | 0.6482 | 0.699 | 0.6949 | 0.7051 |
| 0:59:30 | 0.362 | 0.3617 | 0.3672 | 0.6547 | 0.6559 | 0.6487 | 0.6931 | 0.6965 | 0.7092 |
| 1:00:00 | 0.361 | 0.3616 | 0.3656 | 0.6577 | 0.6578 | 0.6476 | 0.6939 | 0.6955 | 0.7059 |

**MTREC$^{mut}$ (μM $P_i$)**

| Time (hr:min:sec) | H2O Ctrl (Rep A) | H2O Ctrl (Rep B) | H2O Ctrl (Rep C) | +RNA (RepA) | +RNA (RepB) | +RNA (RepC) | -RNA (Rep A) | -RNA (Rep B) | -RNA (Rep C) |
|---|---|---|---|---|---|---|---|---|---|
| 0:00:00 | -4.632 | -3.661 | -3.485 | 8.602 | 9.087 | 6.970 | 16.718 | 18.615 | 20.159 |
| 0:00:30 | -3.926 | -3.397 | -2.735 | 11.204 | 11.072 | 8.910 | 26.158 | 28.408 | 28.716 |
| 0:01:00 | -4.323 | -3.882 | -2.294 | 13.895 | 13.101 | 10.101 | 33.569 | 36.656 | 36.833 |
| 0:01:30 | -5.073 | -3.970 | -2.294 | 14.027 | 14.954 | 12.263 | 40.538 | 44.552 | 43.802 |
| 0:02:00 | -4.676 | -4.279 | -3.794 | 17.512 | 16.498 | 13.013 | 48.037 | 52.536 | 49.272 |
| 0:02:30 | -4.940 | -4.367 | -3.661 | 19.409 | 17.071 | 14.424 | 55.007 | 59.241 | 56.506 |
| 0:03:00 | -4.543 | -4.411 | -2.867 | 20.335 | 18.835 | 16.498 | 60.697 | 66.034 | 63.123 |
| 0:03:30 | -5.293 | -4.852 | -3.132 | 20.997 | 21.085 | 18.041 | 67.975 | 72.254 | 70.049 |
| 0:04:00 | -5.779 | -4.940 | -3.308 | 22.453 | 21.482 | 18.924 | 72.519 | 79.047 | 75.518 |
| 0:04:30 | -5.734 | -4.896 | -3.749 | 24.438 | 23.776 | 20.600 | 79.753 | 85.664 | 80.591 |
| 0:05:00 | -5.602 | -4.940 | -3.794 | 24.879 | 24.217 | 21.482 | 84.473 | 89.810 | 84.429 |
| 0:05:30 | -6.308 | -5.470 | -3.617 | 26.334 | 25.761 | 23.247 | 88.708 | 95.677 | 89.722 |
| 0:06:00 | -5.779 | -5.382 | -3.485 | 28.981 | 28.143 | 24.217 | 93.648 | 99.779 | 96.295 |
| 0:06:30 | -6.087 | -5.646 | -3.000 | 31.495 | 29.069 | 25.452 | 97.839 | 103.705 | 99.559 |
| 0:07:00 | -5.955 | -5.867 | -3.485 | 32.113 | 31.098 | 26.952 | 102.250 | 107.234 | 102.823 |
| 0:07:30 | -5.955 | -5.558 | -3.617 | 34.010 | 32.245 | 28.672 | 106.970 | 112.263 | 107.631 |
| 0:08:00 | -6.528 | -5.779 | -4.411 | 36.392 | 34.363 | 29.731 | 110.675 | 114.689 | 111.910 |
| 0:08:30 | -6.528 | -5.823 | -3.661 | 36.965 | 35.818 | 32.289 | 111.557 | 118.086 | 114.380 |
| 0:09:00 | -6.484 | -6.925 | -4.985 | 38.288 | 37.627 | 33.348 | 116.321 | 119.894 | 118.306 |
| 0:09:30 | -6.308 | -6.396 | -4.191 | 40.229 | 38.377 | 34.980 | 119.718 | 124.526 | 119.365 |
| 0:10:00 | -7.014 | -7.058 | -4.411 | 42.126 | 39.921 | 36.568 | 122.011 | 126.158 | 123.996 |
| 0:10:30 | -6.925 | -7.278 | -4.455 | 44.023 | 41.509 | 37.980 | 123.599 | 128.143 | 126.687 |
| 0:11:00 | -7.102 | -7.190 | -4.985 | 45.876 | 43.538 | 39.921 | 125.408 | 129.422 | 128.055 |
| 0:11:30 | -6.793 | -7.146 | -5.337 | 46.273 | 43.935 | 40.273 | 128.849 | 129.422 | 130.216 |
| 0:12:00 | -6.220 | -7.764 | -5.029 | 48.302 | 44.817 | 42.700 | 130.172 | 131.187 | 132.378 |
| 0:12:30 | -7.278 | -7.719 | -5.426 | 49.404 | 46.449 | 43.802 | 130.613 | 131.495 | 134.495 |
| 0:13:00 | -7.631 | -7.631 | -5.073 | 49.978 | 49.625 | 45.170 | 132.907 | 133.569 | 135.554 |
| 0:13:30 | -7.587 | -7.808 | -4.985 | 52.139 | 52.757 | 46.140 | 133.657 | 133.260 | 134.539 |
| 0:14:00 | -6.661 | -7.543 | -5.249 | 53.507 | 54.698 | 48.346 | 135.951 | 134.010 | 137.097 |
| 0:14:30 | -7.234 | -7.984 | -5.117 | 57.389 | 57.036 | 49.625 | 136.745 | 134.451 | 136.656 |
| 0:15:00 | -7.719 | -7.719 | -4.720 | 58.227 | 58.183 | 50.287 | 136.215 | 136.171 | 138.288 |
| 0:15:30 | -7.896 | -7.808 | -4.940 | 59.462 | 59.771 | 52.007 | 134.848 | 135.730 | 138.377 |
| 0:16:00 | -7.984 | -8.205 | -5.470 | 61.756 | 63.741 | 55.315 | 135.157 | 135.774 | 138.377 |
| 0:16:30 | -6.837 | -8.690 | -5.955 | 62.461 | 63.079 | 58.800 | 136.833 | 134.804 | 139.259 |
| 0:17:00 | -6.308 | -7.896 | -5.514 | 64.976 | 64.755 | 60.741 | 134.760 | 135.686 | 139.524 |
| 0:17:30 | -8.249 | -8.072 | -5.602 | 67.402 | 66.564 | 62.417 | 135.862 | 136.789 | 140.891 |
| 0:18:00 | -8.161 | -8.337 | -6.043 | 68.196 | 67.931 | 64.358 | 134.936 | 137.450 | 139.700 |
| 0:18:30 | -8.116 | -9.219 | -6.176 | 69.696 | 71.284 | 65.152 | 134.804 | 138.068 | 140.494 |
| 0:19:00 | -7.719 | -8.734 | -5.426 | 70.931 | 73.754 | 66.564 | 137.406 | 139.391 | 140.185 |
| 0:19:30 | -7.675 | -8.602 | -5.249 | 72.651 | 74.680 | 67.799 | 136.348 | 137.671 | 139.568 |
| 0:20:00 | -7.984 | -8.161 | -5.867 | 74.063 | 73.401 | 69.563 | 136.833 | 137.362 | 142.126 |
| 0:20:30 | -8.513 | -9.131 | -6.131 | 75.651 | 76.180 | 70.931 | 137.450 | 137.759 | 141.464 |
| 0:21:00 | -8.249 | -9.043 | -5.911 | 78.121 | 77.812 | 72.872 | 136.348 | 138.421 | 140.759 |
| 0:21:30 | -8.734 | -8.249 | -6.881 | 78.209 | 79.709 | 73.489 | 136.656 | 138.156 | 140.803 |
| 0:22:00 | -8.205 | -8.778 | -6.352 | 80.018 | 78.386 | 75.033 | 137.803 | 139.171 | 139.391 |
| 0:22:30 | -7.543 | -8.513 | -6.176 | 80.282 | 82.091 | 75.959 | 138.024 | 139.303 | 141.597 |
| 0:23:00 | -8.205 | -8.999 | -5.734 | 82.532 | 82.885 | 77.989 | 137.450 | 137.891 | 140.450 |
| 0:23:30 | -8.425 | -8.822 | -6.484 | 84.076 | 84.208 | 78.738 | 137.847 | 137.715 | 142.214 |
| 0:24:00 | -8.513 | -8.602 | -5.558 | 84.870 | 85.576 | 78.297 | 137.803 | 137.627 | 140.406 |
| 0:24:30 | -8.116 | -8.690 | -6.131 | 86.458 | 87.252 | 80.415 | 136.965 | 138.333 | 140.494 |
| 0:25:00 | -9.043 | -9.043 | -5.602 | 87.164 | 88.840 | 81.694 | 138.685 | 138.156 | 141.288 |
| 0:25:30 | -8.337 | -9.087 | -5.911 | 88.619 | 88.708 | 83.679 | 139.215 | 139.082 | 141.200 |
| 0:26:00 | -8.513 | -8.910 | -5.779 | 89.987 | 91.266 | 85.399 | 138.024 | 137.936 | 142.832 |

| | | | | | | | | | |
|---|---|---|---|---|---|---|---|---|---|
| 0:26:30 | -9.440 | -8.955 | -5.646 | 91.090 | 91.266 | 86.414 | 137.142 | 138.377 | 142.347 |
| 0:27:00 | -9.660 | -8.513 | -6.396 | 93.119 | 92.192 | 87.164 | 137.230 | 138.112 | 142.303 |
| 0:27:30 | -9.131 | -9.440 | -6.661 | 93.030 | 93.957 | 88.531 | 137.142 | 138.906 | 141.332 |
| 0:28:00 | -9.263 | -9.131 | -6.220 | 95.060 | 94.001 | 89.502 | 137.715 | 138.730 | 141.641 |
| 0:28:30 | -8.469 | -9.396 | -6.793 | 95.677 | 95.192 | 91.707 | 139.082 | 137.671 | 140.979 |
| 0:29:00 | -8.866 | -9.043 | -6.793 | 97.574 | 96.471 | 92.457 | 137.980 | 138.994 | 142.214 |
| 0:29:30 | -8.999 | -8.955 | -6.661 | 97.927 | 97.486 | 93.163 | 137.406 | 138.377 | 141.817 |
| 0:30:00 | -9.307 | -9.484 | -5.734 | 98.897 | 98.765 | 93.560 | 137.715 | 137.803 | 140.847 |
| 0:30:30 | -9.396 | -9.043 | -6.396 | 99.206 | 99.162 | 94.089 | 136.877 | 138.068 | 142.700 |
| 0:31:00 | -9.660 | -9.131 | -6.396 | 99.515 | 100.529 | 94.486 | 136.259 | 138.509 | 141.861 |
| 0:31:30 | -10.190 | -9.440 | -7.190 | 100.838 | 101.985 | 96.692 | 135.906 | 138.244 | 140.803 |
| 0:32:00 | -9.572 | -9.087 | -7.675 | 100.926 | 102.161 | 97.353 | 136.348 | 137.097 | 140.406 |
| 0:32:30 | -8.955 | -8.955 | -7.014 | 102.514 | 102.823 | 97.927 | 138.156 | 138.421 | 140.670 |
| 0:33:00 | -9.352 | -9.749 | -6.352 | 104.014 | 103.749 | 98.897 | 136.656 | 137.671 | 141.156 |
| 0:33:30 | -9.925 | -9.219 | -6.396 | 104.543 | 103.838 | 98.941 | 135.554 | 137.671 | 143.185 |
| 0:34:00 | -9.969 | -9.396 | -7.234 | 103.661 | 105.293 | 100.353 | 137.627 | 137.539 | 141.288 |
| 0:34:30 | -10.057 | -9.881 | -7.587 | 105.646 | 106.793 | 101.235 | 137.583 | 136.921 | 140.097 |
| 0:35:00 | -10.234 | -10.146 | -8.072 | 106.661 | 107.675 | 101.676 | 136.833 | 137.671 | 140.406 |
| 0:35:30 | -10.101 | -9.749 | -7.367 | 105.249 | 106.661 | 103.088 | 137.539 | 137.450 | 139.965 |
| 0:36:00 | -9.307 | -10.322 | -7.631 | 107.852 | 106.837 | 102.647 | 137.936 | 136.392 | 140.229 |
| 0:36:30 | -9.396 | -9.837 | -7.675 | 108.558 | 108.558 | 103.044 | 137.671 | 137.274 | 142.347 |
| 0:37:00 | -10.278 | -9.881 | -7.234 | 108.205 | 109.881 | 104.940 | 136.171 | 136.877 | 141.773 |
| 0:37:30 | -10.675 | -9.749 | -7.940 | 108.646 | 110.587 | 105.999 | 135.818 | 136.965 | 141.597 |
| 0:38:00 | -10.101 | -10.190 | -8.425 | 109.087 | 110.013 | 105.734 | 136.259 | 137.053 | 140.847 |
| 0:38:30 | -10.013 | -10.101 | -8.116 | 110.057 | 110.719 | 106.352 | 138.333 | 136.127 | 141.067 |
| 0:39:00 | -9.925 | -10.366 | -7.764 | 110.807 | 110.234 | 105.911 | 137.009 | 136.392 | 140.891 |
| 0:39:30 | -9.969 | -10.057 | -6.705 | 110.895 | 110.895 | 106.661 | 137.274 | 136.877 | 141.200 |
| 0:40:00 | -10.543 | -10.366 | -7.102 | 111.469 | 112.086 | 106.925 | 136.039 | 136.568 | 141.200 |
| 0:40:30 | -10.851 | -10.410 | -7.940 | 112.616 | 113.542 | 107.631 | 136.171 | 137.494 | 141.597 |
| 0:41:00 | -10.278 | -10.322 | -8.425 | 112.395 | 113.145 | 108.558 | 136.039 | 136.392 | 141.861 |
| 0:41:30 | -9.175 | -10.366 | -8.293 | 111.822 | 113.189 | 108.602 | 136.303 | 135.686 | 140.318 |
| 0:42:00 | -9.793 | -10.322 | -8.381 | 112.616 | 113.719 | 108.337 | 135.995 | 136.745 | 142.347 |
| 0:42:30 | -10.851 | -10.807 | -7.808 | 113.322 | 112.880 | 109.704 | 134.760 | 136.789 | 141.067 |
| 0:43:00 | -10.057 | -10.984 | -8.513 | 113.719 | 115.218 | 109.704 | 135.377 | 136.215 | 138.641 |
| 0:43:30 | -10.851 | -11.557 | -8.513 | 113.366 | 114.645 | 109.616 | 135.906 | 136.745 | 138.906 |
| 0:44:00 | -9.749 | -11.425 | -8.734 | 113.983 | 114.954 | 109.175 | 137.627 | 136.789 | 139.612 |
| 0:44:30 | -10.410 | -10.895 | -8.072 | 113.630 | 115.307 | 110.587 | 135.951 | 136.745 | 140.538 |
| 0:45:00 | -10.675 | -11.028 | -9.307 | 115.042 | 116.321 | 111.160 | 136.127 | 135.906 | 141.200 |
| 0:45:30 | -10.807 | -9.396 | -9.263 | 114.336 | 116.718 | 111.028 | 134.760 | 136.215 | 138.641 |
| 0:46:00 | -10.101 | -10.984 | -8.778 | 114.601 | 114.160 | 110.410 | 135.862 | 135.906 | 139.832 |
| 0:46:30 | -10.543 | -11.116 | -9.440 | 114.336 | 115.571 | 110.984 | 135.686 | 135.465 | 140.185 |
| 0:47:00 | -11.028 | -11.072 | -8.558 | 114.998 | 116.542 | 111.028 | 135.554 | 135.509 | 140.494 |
| 0:47:30 | -11.425 | -11.116 | -9.307 | 114.027 | 115.880 | 113.366 | 134.010 | 135.333 | 140.097 |
| 0:48:00 | -10.984 | -11.072 | -9.660 | 115.307 | 118.791 | 112.528 | 135.377 | 135.068 | 138.818 |
| 0:48:30 | -10.895 | -11.381 | -9.704 | 115.086 | 116.718 | 111.381 | 136.877 | 135.421 | 137.980 |
| 0:49:00 | -11.734 | -12.351 | -9.396 | 114.689 | 116.850 | 111.248 | 134.274 | 135.995 | 139.744 |
| 0:49:30 | -11.381 | -11.204 | -9.660 | 115.395 | 119.100 | 112.616 | 134.583 | 136.833 | 140.185 |
| 0:50:00 | -10.940 | -12.263 | -8.513 | 115.395 | 116.101 | 112.748 | 134.980 | 136.745 | 138.509 |
| 0:50:30 | -11.204 | -12.175 | -9.925 | 114.998 | 116.983 | 112.483 | 134.760 | 135.862 | 140.318 |
| 0:51:00 | -11.778 | -11.204 | -9.484 | 115.571 | 116.542 | 113.322 | 135.245 | 136.612 | 139.524 |
| 0:51:30 | -12.086 | -12.042 | -9.616 | 115.439 | 118.483 | 112.880 | 135.068 | 134.671 | 139.479 |
| 0:52:00 | -11.778 | -12.263 | -9.572 | 115.483 | 117.247 | 111.998 | 134.407 | 135.112 | 137.539 |
| 0:52:30 | -10.940 | -11.601 | -10.234 | 114.954 | 115.748 | 113.674 | 136.171 | 136.480 | 139.171 |
| 0:53:00 | -12.131 | -12.042 | -10.454 | 115.262 | 117.468 | 110.895 | 135.024 | 135.598 | 139.876 |
| 0:53:30 | -11.954 | -12.395 | -9.528 | 116.895 | 117.292 | 111.866 | 134.451 | 134.318 | 141.112 |
| 0:54:00 | -13.013 | -12.704 | -10.410 | 116.012 | 118.174 | 113.454 | 133.436 | 135.818 | 138.774 |
| 0:54:30 | -11.998 | -12.439 | -10.498 | 116.850 | 117.292 | 112.086 | 133.304 | 134.715 | 137.539 |

| | | | | | | | | | |
|---|---|---|---|---|---|---|---|---|---|
| 0:55:00 | -12.219 | -11.998 | -11.028 | 115.704 | 117.159 | 112.704 | 135.112 | 134.980 | 137.142 |
| 0:55:30 | -11.866 | -12.748 | -10.675 | 116.542 | 117.159 | 114.027 | 136.568 | 135.333 | 138.862 |
| 0:56:00 | -12.528 | -13.101 | -10.587 | 115.704 | 117.733 | 112.307 | 133.480 | 134.539 | 141.067 |
| 0:56:30 | -12.483 | -12.483 | -10.454 | 116.321 | 117.865 | 112.528 | 134.054 | 134.715 | 138.950 |
| 0:57:00 | -12.792 | -12.616 | -11.116 | 116.718 | 117.600 | 113.763 | 134.142 | 135.598 | 138.421 |
| 0:57:30 | -12.616 | -13.101 | -10.763 | 117.071 | 118.394 | 112.572 | 133.966 | 135.112 | 139.215 |
| 0:58:00 | -12.439 | -12.836 | -11.601 | 116.542 | 117.203 | 112.660 | 134.627 | 134.583 | 138.774 |
| 0:58:30 | -12.131 | -12.969 | -11.337 | 114.733 | 116.762 | 114.027 | 134.848 | 134.936 | 137.230 |
| 0:59:00 | -12.395 | -13.057 | -11.248 | 116.409 | 118.350 | 113.145 | 135.554 | 133.745 | 138.244 |
| 0:59:30 | -13.101 | -13.233 | -10.807 | 116.012 | 116.542 | 113.366 | 132.951 | 134.451 | 140.053 |
| 1:00:00 | -13.542 | -13.277 | -11.513 | 117.336 | 117.380 | 112.880 | 133.304 | 134.010 | 138.597 |

**Table S6. | ATPase assay raw data and calculations.**
